# Un1Cas12f1 and Cas9 gene drive in HSV1: viruses that ‘infect’ viruses

**DOI:** 10.1101/2023.12.04.569968

**Authors:** Qiaorui Yao, Zhuangjie Lin, Keyuan Lai, Xianying Zeng, Guanxiong Lei, Tongwen Zhang, Hongsheng Dai

## Abstract

Synthetic CRISPR-Cas9 gene drive has been developed as a potential tool to control harmful species. However, Cas9 gene drive faces high resistance rate and mitigation strategies developed so far are difficult to implement. Furthermore, studying the resistance to gene drive is time consuming and challenging in higher organisms. We here tackled these two challenges simultaneously by generating Cas9 and Un1Cas12f1 gene drive in a fast-replicating DNA virus, HSV1. We assessed the transmission dynamics and resistance formation through phenotypical staining and next-generation sequencing, and demonstrated that HSV1 supported fast and effective transmission of gene drives, and the Un1Cas12f1 gene drives yielded greater conversion and lower resistance than did the Cas9 gene drives. This positions the Un1Cas12f1 gene drive as a promising alternative, and HSV1 emerges as a dependable and swift platform for gene drive assessment. The gene drive viruses function like pathogens that specifically infect viruses, offering potential applications in attenuating viral infections.

## Introduction

Gene drive is a super mendelian inheritance in which particular genetic sequences are preferentially passed to the next generation, resulting in the dominant spread of corresponding traits within a target population^1^. Natural gene drives are mainly “selfish genes”, such as transposons, homing endonuclease genes, and meiotic drive genes, and they have been modified to reduce the survival or reproduction of pests^2^. However, these agents were not broadly adopted due to their low efficiency, poor operationality, and, most importantly, inevitable emergence of resistance^3^.

The CRISPR[Cas system has galvanized a new wave of interest in developing CRISPR-based synthetic gene drives as a tool for population control^2^. CRISPR[Cas gene-editing technique involves a Cas nuclease and a short guide RNA (sgRNA) ^4–6^. Cas9 is a widely used Cas nuclease and serves as a model for other Cas nucleases. The sgRNA has two components: a 20-nucleotide region that matches the target DNA and a scaffold that interacts with Cas9. The sgRNA guides Cas9 to a specific location in the genome by forming base pairs with the target DNA. Cas9 is then allosterically activated by the protospacer adjacent motif (PAM), a 2–6 bp sequence immediately following the target DNA sequence, and cut inside the DNA target 3-4 bp upstream of the PAM, creating a double-strand break (DSB)^7,8^.

The CRISPR-Cas9 gene drive in general are composed of: (1) Cas9 nuclease and sgRNA coding sequences and (2) adjacent DNA segments (∼1 kb in length on each side) that match the locus targeted by sgRNA ^9,10^. The CRISPR[Cas9 gene drive specifically induces DSB on the targeted allele and subsequently guides homology-directed repair (HDR) by using itself as the template, resulting in the integration and spread of the gene drive over generations. Gene drives based on CRISPR[Cas9 have been established in yeast^11,12^, Drosophila^9,10^, mosquitoes^13,14^, mice^15^ and a human DNA virus^16^.

The DSBs induced by gene drive may also be repaired via nonhomologous end joining (NHEJ), which directly reconnects broken ends of sgRNA target DNA and often introduces small insertions or deletions (indels). For this reason, CRISPR[Cas9 gene drives would eventually be stopped by resistance alleles, which mainly arises from mutations in the target sequence ^17,18^. Several methods have been tested to overcome resistance to Cas9 gene drive, such as targeting multiple regions that are conserved or essential for the organism^19–22^. However, implementing these approaches is cumbersome and not necessarily effective. Another option is to explore Cas nucleases that would mechanistically induce fewer mutations at the target site after DSB repair. Cas12a is the only other Cas nuclease implemented for gene drive thus far and has been shown to propagate efficiently in yeast and fruit flies^23^ ^24^. However, Cas12a cuts DNA within the sgRNA target sequence ^25^ and is very sensitive to mismatches between the sgRNA and the target DNA ^26^. In line with these properties of Cas12a, Víctor López Del Amo *et al* reported that the Cas12a gene drive produced resistant alleles in 5-8% of the target population^24^.

The Cas9 nuclease is a large protein of 1368 amino acids that poses challenges for viral vector delivery^27^. To overcome this limitation, smaller Cas nucleases have been searched for or engineered, resulting in a variety of alternatives ^28–34^. Un1Cas12f1 is a type-V CRISPR nuclease from archaea, although being 1/3 the size of Cas9, it achieved efficient and specific genome editing with optimized sgRNA^28,31,33^. Unlike Cas9, Un1Cas12f1 cleaves dsDNA outside the sgRNA target and PAM, so the target site is likely preserved even after indel mutations by NHEJ^28^. Because of its smaller size and different DNA cleavage pattern, Un1Cas12f1 may offer some advantages over Cas9. However, Un1Cas12f1-based gene drive has not been reported thus far, and its performance remains unknown.

Gene drive resistance has been studied with modeling ^19,20^ or experimentally by tracking gene drive resistance in insects ^17,18,35,36^. The relatively slow pace of sexual reproduction in insects (i.e., one generation needs 10 days for flies housed at 25°C), however, necessitates a long time for genetic changes to spread and accumulate within a population, making such experiments very time-consuming and difficult to scale up. A new gene drive carrier that can be conveniently manipulated and support the rapid spread of a gene drive would be highly helpful.

Human herpes simplex virus 1 (HSV1), a member of the family Herpesviridae, has a linear double-stranded DNA (dsDNA) genome containing 74 open reading frames^39^. The dsDNA genome of HSV1 can be edited by CRISPR[Cas9 ^37,38^. Like many other dsDNA viruses, HSV1 viruses are considered monoploid because each virion contains only one copy of the genome. However, HSV1 viruses usually produce many copies of the genome that coexist in the nucleus before being packaged into new virions, and one host cell can be infected simultaneously by multiple HSV1 virions. These biological features of HSV1 thus create opportunities for genetic materials to be transferred horizontally between viral genomes (Supp Fig 1). The HSV1 dsDNA genome (∼150 kb) contains many dispensable regions that can potentially be replaced with gene drive elements^39^. Furthermore, HSV1 infects a broader range of host cells, produces thousands of genome copies in one cell, and finishes an infection/replication cycle in 12 hours. These features make HSV1 an promising vector for testing gene drive and monitoring resistance.

**Fig. 1.**
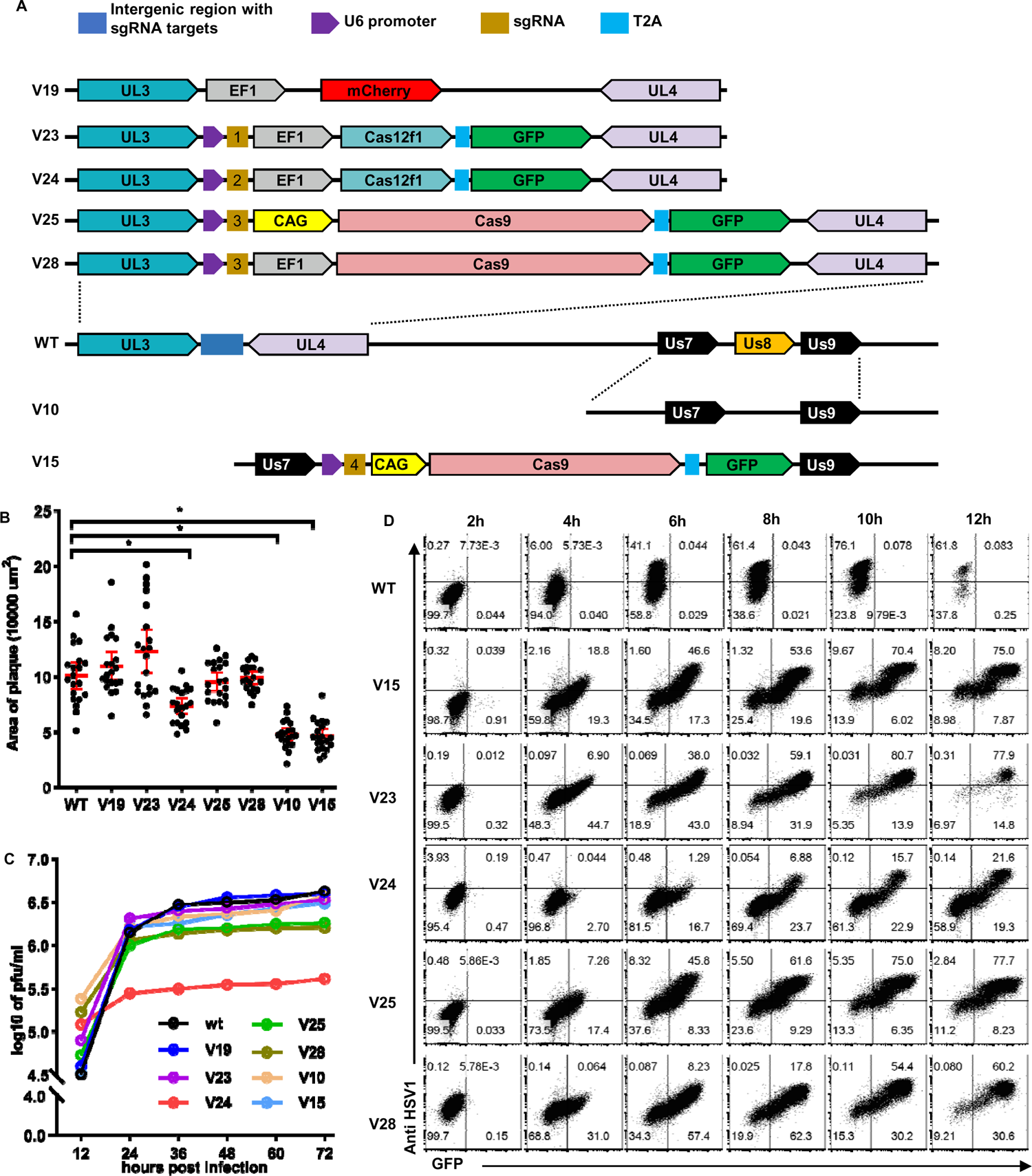
Generation and basic properties of gene drive viruses. A, Schematic representation of the genetic organization of plasmids and the resultant recombinant HSV1 viruses. B, Plaque sizes of HSV-1 viruses on Vero cells at 48 hours post infection. Twenty plaques from each virus were randomly picked, and the area for each plaque was calculated and plotted. The mean with 95% CI was marked for each virus. * P<0.05. C, One-step growth curve of HSV-1 viruses. D, Expression of GFP and HSV1 antigens at different time points after BHK cells were infected with HSV1 viruses (MOI=2).

In this study, we set to create the first Un1Cas12f1 gene drive and test the performance of HSV1 as the carrier of gene drive. As the result, we created HSV1 viruses that carried either Cas9 or Un1Cas12f1-based gene drive, evaluated their transmission dynamics, and investigated the difference between Cas9 and the Un1Cas12f1 gene drive in generating resistance for the first time.

## Results

### Generation of HSV1 gene drive viruses

The HSV1 genes UL3 and UL4 play no essential role in HSV1 replication, and the UL3-UL4 intergenic region has been reported to be a safe harbor that tolerates the insertion of large DNA fragments^40^. To generate gene drive viruses with uncompromised fitness, we created gene drive cassettes to target the UL3-UL4 intergenic region (Fig 1A). The two Un1Cas12f1 genes drive V23 and V24 produce sgRNAs targeting different sites within the UL3-UL4 intergenic region (Supp. Fig 2). The two Cas9 genes drive V25 and V28 target the same site, but the expression of Cas9 is driven by the CAG and EF1 promoters, respectively (Fig 1A, Supp. Fig 2). HSV1 Us8 gene encodes glycoprotein E (gE), although not essential for viral replication, its deficiency impairs the transmission of HSV1 between cells^39,41^. To make gene drive viruses with defined attenuation, we constructed gene drive cassette V15 to target HSV1 Us8 gene. We accordingly made two control cassettes: V19 having the UL3-UL4 intergenic region replaced with mCherry, and V10 with large deletion within Us8 coding sequences (Fig 1A).

Viruses V19 and V10 were generated by CRISPR-Cas9 assisted recombination (Supp Fig. 3A), and the five gene drive viruses were generated by transfecting 293T cells with gene drive plasmids described above and subsequently infecting with the HSV1 strain F (wildtype, WT hereafter) (Supp Fig. 3B). Although autonomous homology recombination (in the absence of Cas9 and sgRNA) has long been used to modify the genome of herpesviruses ^42^, it occurred at very low frequency as reported in many other studies ^37,38^ and hardly contributed to the generation of recombinant viruses in our reverse genetics system (Supp Fig 2C). Single clones of all engineered viruses were easily purified by 3-4 rounds of plaque assay (Supp Fig 3D) and confirmed by junctional PCR and sequencing (Supp Figure 3E). The successful generation of gene drive viruses at high efficiency attested that the gene drive cassettes converted the target loci in the HSV1 genome as designed.

**Fig. 2.**
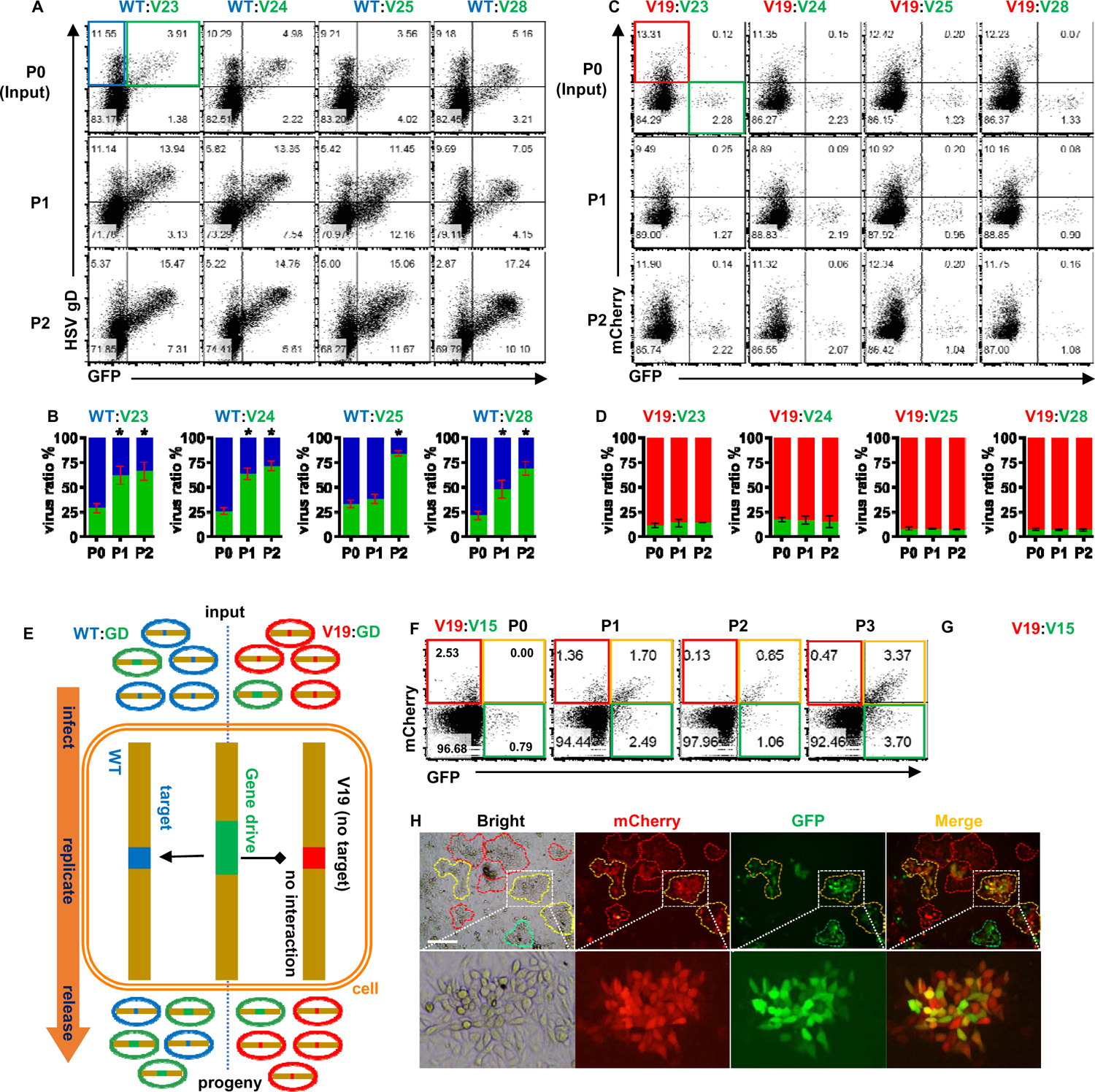
Spread of gene drive in HSV1 viruses. A and B, WT (GFP-, gD+) and gene drive viruses (GFP+, gD+) coinfection of vero cells were performed at MOI=5 with an initial ratio of WT: gene drive viruses close to 4:1, and passaged for 2 generations. The ratio of each virus was measured before coinfection (P0) and after each passage by infecting fresh BHK cells for 12 hours and labeling them with antibodies. Representative flow cytometry graphs (A) and statistical summary (B) of 3 experiments are shown. * p<0.05 compared with P0. C and D, V19 (mCherry+) and gene drive virus (GFP+) coinfection of Vero cells were conducted as in A and B. Representative flow cytometry graphs (C) and statistical summary (D) of 3 experiments are shown. E, Schematic diagram showing the process and consequence of the spread of gene drive viruses with WT or V19 viruses. F-H, V19 (mCherry+) and V15 (GFP+) coinfection of Vero cells were conducted as in A and B, but for 3 passages. Flow cytometry graphs (F) and the average ratio of each strain (H) from two independent experiments was shown. H, P1 supernatant of V19:V15 coinfection was diluted and plated on Vero cells for 48 hours. Plaques were imaged for the expression of fluorescent proteins. Scale bar, 500 µM.

**Fig. 3.**
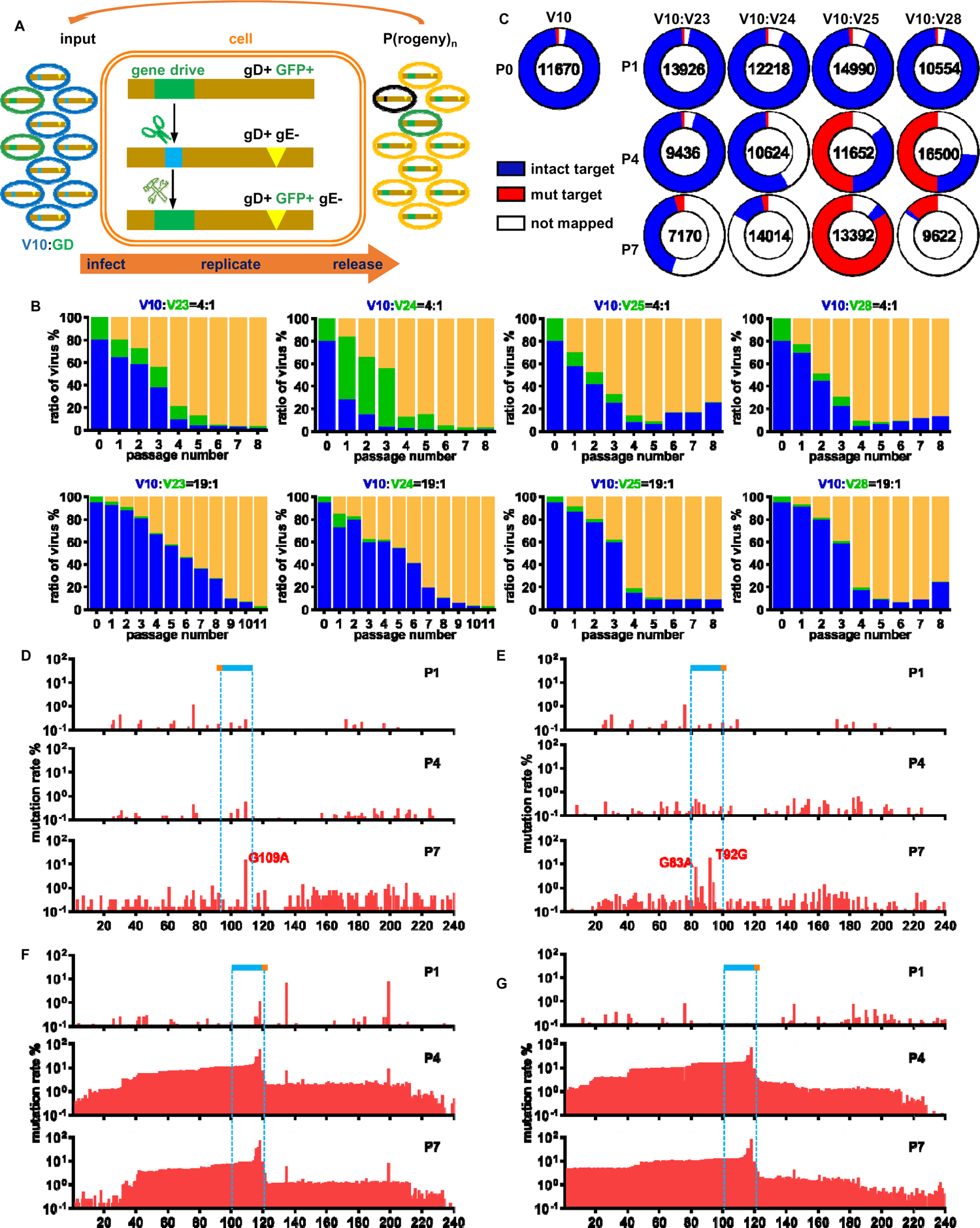
The transmission dynamics of the HSV1 gene drive viruses and the rise of resistant clones. A, Schematic diagram showing the procedure and possible consequences of V10 (gD^+^ gE^-^) and gene drive viruses (gD^+^ GFP^+^ gE^+^) coinfection. V10 can be converted by newly generated gene drive viruses (gD^+^ GFP^+^gE^-^) at late stage of coinfection, and may produce variants resistant (gD^+^ gE^-^) to Cas nuclease. B, V10 and gene drive coinfection of Vero cells was conducted at MOI=5 with the initial ratio of V10: gene drive =4:1 or 19:1. The ratio of each virus after each passage was determined by infecting and staining BHK cells, and summarized here. The blue, green and yellow bars represent V10, the input gene drive and the new gene drive viruses, respectively. C, NGS sequencing results for the UL3-UL4 intergenic region amplified from input viruses (P0) and coinfection supernatants. The scaled cycle showed the relative ratio for three types of reads, and inside each cycle were the total number of amplicon reads for each sample. D-G. Distribution and frequency of mutations accumulated at the UL3-UL4 intergenic region for coinfection V10:V23 (D), V10:V24 (E), V10:V25 (F) and V10:V28 (G). The locations of the sgRNA target sequences and the PAM are marked with blue‒red bars.

### Basic properties of HSV1 gene drive viruses

Virus V19 and V23 infected and replicated like WT (Fig. 1B, C), confirming that the UL3-UL4 junction is suitable for editing. Gene drive viruses V25 and V28 produced plaques similar in size to those of the WT, but their titers were slightly lower than those of the WT (Fig. 1B, C). V24 purified in this study produced smaller plaques and a 10-fold lower titer. V23 and V24 had nearly identical gene drives, therefore, the decreased replication of V24 was unlikely caused by the gene drive cassette itself. Due to the large size of the HSV1 genome, no attempt was made to identify the cause of the attenuated phenotype of V24, or to purify other clones of V24 with more favorable replication. Instead, this V24 clone provided an opportunity to test if an attenuated virus could spread gene drive. HSV1 viruses V10 and V15 lost the expression of gE as expected (Supp Fig. 4). Although V10 and V15 produced viral titers similar to those of the WT, they both formed much smaller plaques (Fig. 1B, C). These results suggested that the V15 gene drive cassette disrupted gE and contributed to the reduced fitness of gene drive viruses.

The five gene drive viruses started to express GFP in the early stage of infection, and GFP increased in proportion to the level of HSV1 antigens as infection proceeded (Fig. 1D). GFP is linked downstream of Cas 9 or Un1Cas12f1 through T2A, and thus reports the production of Cas9 or Un1Cas12f1 (Fig. 1A). Although Un1Cas12f1 is approximately 1/3 of the size of Cas9, the additional length of Cas9 seemed to have little effect on the expression of GFP, as the levels and kinetics of GFP for all the gene drive viruses did not significantly differ (Fig. 1D).

### Spread of gene drive in HSV1 viruses

Multiple HSV1 virions can infect the same host cell and initiate their own replication, providing the opportunity for viruses with different genomes to interact (Supp Fig 1). We therefore co-infected vero cells with WT virus (gD^+^ GFP-) and each strain of gene drive viruses (gD^+^, GFP^+^), respectively, and determined the composition of progeny viruses after each cycle of coinfection via fluorescence staining (Fig. 2A). Although the gene drive viruses V23, V24, V25, and V28 replicated at a level close to or even lower than that of the WT (Fig. 1C), their proportion in the viral population increased steadily from ∼25% at P0 to ∼70% at P2 (Fig. 2B). To validate if such a biased expansion of gene drive viruses is related to the UL3-UL4 intergenic region, for which the four gene drives targeted, we coinfected each of the four gene drive viruses (GFP^+^ and mCherry^-^) with V19 (GFP^-^ and mCherry^+^) (Fig. 2C). V19 had the UL3-UL4 intergenic region replaced with mCherry (Fig. 1A). Under this condition, the percentage of each strain of gene drive virus (including the weakened V24) after each passage was within a range close to their initial proportion in the viral population (Fig 2C, D), and no viruses expressing both mCherry and GFP were observed (Figure 2C, D, Supp Fig 5). The biased expansion of gene drive viruses during coinfection with WT is the result of the UL3-UL4 intergenic of WT being converted to gene drive (Fig 2E).

However, using WT as the recipient of gene drive produces progeny gene drive viruses indistinguishable from the input gene drive virus V23, V24, V25 and V28. We thus performed coinfection of gene drive virus V15 (mCherry-GFP+) with V19 (mCherry+GFP-), which had intact Us8 gene to be targeted by V15 (Fig 1A). This time, transmission of gene drive V15 was directly observed as the rise of viruses expressing both GFP and mCherry (Fig. 2F-H). Gene drive viruses, including GFP^+^ mCherry^-^ and GFP^+^ mCherry^+^ populations, comprised 90% of all viruses after two passages (Fig. 5H), an efficiency not inferior to that of gene drive viruses with normal phenotype (Fig. 2B, D).

### Transmission dynamics of the Un1Cas12f1 and Cas9 gene drives

Thus far, we have demonstrated that the two Un1Cas12f1 gene drive viruses (V23 and V24) and the two Cas9 gene drive viruses (V25, V28) carried functional gene drive targeting UL3-UL4 intergenic region. To sort out differences between Un1Cas12f1 and Cas9 gene drives, we coinfected V10 with each gene drive virus at initial ratios of 4:1 and 19:1 to simulate different levels of transmission threshold, and tracked the conversion of V10 to gene drive after each passage (Fig. 3A). Because these four gene drives had the same homologous arms, any discrepancy shown in their transmission should originate from Cas9 and Un1Cas12f1.

The new gene drive viruses were gD+ GFP+ gE- and readily distinguishable from the input gene drive viruses, which were gD+ GFP+ gE+ (Supp Figs. 6-9). At the initial ratio of 4:1, all four gene drive viruses converted V10 steadily and became dominant after 4 passages (Fig 3B, Supp Fig 10). The Un1Cas12f1 and Cas9 gene drives had similar conversion rates until P5, when the Cas9 gene drives encountered resistance and some V10 rebounded. At the initial ratio of 19:1, the Un1Cas12f1 gene drives spread slower than the Cas9 gene drives, and converted almost all V10 by P11. The lower starting ratio did not affect the two Cas9 gene drivers very much. They still reached equilibrium with V10 at P5, leaving a large fraction of V10 (up to 25%) unconverted that may be resistant variants to Cas9-gRNA (Fig. 3B, Supp Fig 10).

### Genetic changes introduced by Cas9 and Un1Cas12f1 gene drives

To examine how the spread of gene drive could change the genome of V10 viruses and make them resistant, we performed NGS amplicon sequencing of the UL3-UL4 intergenic region from the supernatants of the 4:1 coinfection at P0 (input), P1, P4, and P7 (Supp. Fig 2, and Supp Fig 11). We used a short PCR extension time (20 seconds) to avoid amplifying gene drive viruses, however, they could attract the primers and cause false priming or primer dimers. The NGS data showed how different gene drives affected V10 viruses over time (Fig. 3C). At P1, V10 viruses were abundant and mostly unchanged from P0. At P4 and P7, gene drive viruses took over and interfered with PCR, causing many unmapped NGS reads. The mapped reads either had intact or mutated sgRNA targets, depending on the gene drive. Un1Cas12f1 gene drives (V23 and V24) preserved most of the sgRNA targets in V10, while Cas9 gene drives (V25 and V28) disrupted most sgRNA targets in V10.

We examined the UL3-UL4 intergenic region for mutations in all the mapped reads and calculated the cumulative mutation rate for each base (Fig 3D-G). Mutations can arise naturally during genome replication or from error-prone NHEJ repair of a DSB. NHEJ often produces a high frequency of short indels at DSB junctions. Surprisingly, almost no indels were found in V10 viruses from the six V10:V23 and V10:V24 coinfection pools, instead, point mutations were enriched within the Un1Cas12f1 sgRNA target in P4 and P7(Fig. 3D, E). The G109A mutation was on the V23 sgRNA target, and its frequency in mapped reads (0.3% of P1, 0.47% of P4, and 8.3% of P7) increased as V10:V23 coinfection progressed (Fig. 3D, Supp. Fig. 1). The G83A and T92G mutations, which disrupt the V24 sgRNA target, were the two main mutations that accumulated during V10:V24 coinfections (Fig. 3E, Supp. Fig. 1). The two Cas9 gene drive V25 and V28 introduced many short indels to the V10 genome, and these indels were centered on the third nucleotide upstream of the PAM (Fig. 3F, G). Large deletions (loss >50 nt) appeared in 3-7% of the reads in the P4 and P7 pools. However large deletions were not equally distributed around the cut site of Cas9. The 5’ side of the breakpoint was truncated more frequently than the 3’ side, and there were almost no reads with large deletions across both sides of the breakpoint (Supp Fig. 12). NGS sequencing results were generally consistent with the dynamic change of V10 viruses (Fig 3B), and collectively confirmed that Un1Cas12f1 gene drives showed high conversion rate and low occurrence of resistance, and that Cas9 gene drive caused high rate of resistance.

## Discussion

In this study, we generated for the first time HSV1 viruses that carried functional Cas9 and Un1Cas12f1 gene drives. Due to the rapid replication of HSV1, the spread of gene drive proceeded at a speed not achieved in any other organisms tested thus far ^12,15,16,24^. Both Cas9 and Un1Cas12f1 gene drives were highly effective and capable of converting targets even when their initial presence was low. The Cas9 gene drives produced a high percentage of resistant viruses by introducing mutations at the target sites, the Un1Cas12f1 gene drives instead were able to convert almost all targets without causing many resistant clones.

Un1Cas12f1 cut the target DNA at 22 nt and 24 nt upstream of the PAM ^43^, intrinsically preserves the sgRNA target sites and allows multiple editing at the same locus ^28,44^. Previous studies showed that Un1Cas12f1 caused fewer mutations than Cas9 while editing the genome of 293T cells and the resultant mutations were mainly deletions around the target^28,44^. We in the first Un1Cas12f1 viral gene drive study barely observed any deletional mutations near the target sites of Un1Cas12f1 gene drive. It is unlikely that we failed to detect such indels, because our amplicon NGS sequencing protocol reliably detected a wide range of indels caused by Cas9 gene drives. Un1Cas12f1 cuts dsDNA and leave sticky ends ^43^, which had been reported to help with HDR when there is a matching template ^45^. HSV1 viruses are essentially multiploid when they replicate inside cells. This means that there are many templates available for HDR, both from the wild-type and the gene drive versions of HSV1. The sticky ends generated by Un1Cas12f1 and the availability of ample repair templates may contribute to nearly 100% of the conversion rate of the Un1Cas12f1 gene drives in HSV1. It would be worthwhile to investigate whether Un1Cas12f1 gene drive could achieve such a high conversion rate in other organisms.

We noticed that Cas9 gene drive V25 and V28 had produced some resistant V10 clones that had large deletions in their genomes. These large deletions occurred dominantly at the 5’ side of the break point of the target sequences. Unidirectional editing is a property of type I-E CRISPR nucleases Cas3, which induces large deletions mostly upstream of the PAM^46^. However, this property has not been previously reported for Cas9. Since V25 and V28 targeted the same sequences, we could not rule out that such a deletion pattern is specific to this target site. Testing Cas9 gene drives targeting other regions of the HSV1 genome may help answer if Cas9 preferentially introduce large deletions to the 5’ side of DSB.

The fitness costs of gene drives in insects and other higher organisms often hinder the spread of gene drives^47^. Gene drive viruses V24 and V15 had reduced fitness, however, they efficiently spread the gene drive at a rate not inferior to that of V23. HSV1 viruses reproduce thousands of copies of genome in host cells. The presence of coexisting WT viruses may compensate for the defects of the gene drive viruses and explain why mild fitness loss did not affected the spread of gene drive in viral population.

Cas9-based gene drive has been reported for HCMV^16^. Although both HCMV and HSV1 are herpesviruses, HSV1 replicates much faster and at higher titers than HCMV^48^. We initially suspected that the short replication cycle of HSV1 might pose a challenge for the transmission of gene drive, as it might be too short to complete a multistep process involving the production of sgRNA and Cas nuclease, cleavage of the target, and DSB repair by HDR. Both the Cas9 and Cas12f1 gene drives spread very efficiently among HSV1 viruses, suggesting that such viral gene drives would also work for other herpesviruses.

Overall, our study presented the first Un1Cas12f1-based gene drive and established HSV1 as a useful vector for studying horizontal gene transfer. The Un1Cas12f1 gene drive demonstrated high efficiency and low resistance and could have potential in controlling pathogens and pests. HSV1 is a safe and versatile vector, as engineered HSV1 viruses have been approved for treating cancer^49^ and genetic diseases^50^. Our study opens new possibilities for using the HSV1 gene drive for the prevention and treatment of diseases.

**Notes:** Immediately after we posted our manuscript on bioRxiv, Dr Marius Walter et al posted their development and characterization of Cas9-based HSV1 gene drive viruses (https://www.biorxiv.org/content/10.1101/2023.12.07.570711v3). While targeting at a different locus of HSV1 genome, they reported that “20% of the remaining target sites had been mutated by NHEJ” after being edited by Cas9 gene drive. It is in line with our findings regarding HSV1 Cas9 gene drives.

## MATERIALS AND METHODS

### Cells and viruses

HEK293T, BHK-21, and Vero cells and the HSV-1 F strain were obtained from American Type Culture Collection (ATCC). All cells were cultured in DMEM supplemented with 10% fetal bovine serum (FSP500, ExCell Bio) and 10000 U/ml penicillin/streptomycin (BL505A, Biosharp) and maintained at 37°C with 5% CO2.

### Construction of gene drive cassettes

sgRNA targets were selected using the online tool CRISPOR (http://crispor.tefor.net/), and all plasmids were designed using SnapGene® software (Dotmatics; available at snapgene.com). Cas9 was amplified from px330 (Addgene), and Un1Cas12f1 was derived from the optimized Un1Cas12f1 sequence^28^ and synthesized (General Biol). The vector backbone and insert fragments were amplified with Q5 high-fidelity DNA polymerase (M0491L, NEB) and ligated together with the ClonExpress® Ultra One Step Cloning Kit (C112-02, Vyzame). Ligation products were subsequently transformed with DH10B competent bacteria, after which single clones were picked and validated via Sanger sequencing (Tsingke). The sequences and annotations of all the plasmids used in this study are provided in the supplemental material.

### Generation of the HSV1 gene drive virus

First, 293T cells were transfected with gene drive plasmids precomplexed with PEI (408727-100ML; Sigma[Aldrich), cultured for 12 hours, and subsequently infected with the HSV-1 F strain (MOI =0.1). The recombination of plasmids and viruses was allowed to continue for 48 h, after which the virus-containing supernatants were harvested. To isolate recombinant viruses, a single layer of Vero cells was infected with sequentially diluted supernatant for 1 hour and overlaid with 1% methylcellulose (C6333-250 g, Macklin) solubilized in DMEM. Individual plaques expressing the reporter gene were picked at 48 hours and propagated and purified by an additional 3 to 4 rounds of plaque assay. The genomic DNA of individual clones was extracted with a DNA extraction kit (DC102, Vazyme), amplified with primers (Supplemental Table 1), and validated by Sanger sequencing (Tsingke).

### Titration of Viruses

Viral titration was conducted as previously described ^39^. Briefly, 2×10^4^ Vero cells were incubated with sequentially diluted viruses at 37°C for 4 hours. Human gamma globulin, which contains polyclonal antibodies against HSV1, was added to the culture to a final concentration of 0.5% (v/v). After 48 hours, the plaques were counted under an inverted microscope.

### Imaging

Pictures of viral plaques were taken at 48 hours post infection with a Nikon Eclipse Ts2-FL inverted microscope. To compare the plaque sizes of the different viruses, 20 plaques were randomly picked from each virus, and the area of each plaque was calculated using Nikon acquisition software NIS-Elements BR 5.30.0364.

### Purification of viruses

To prepare virus stocks, BHK-21 or Vero cells were infected with the HSV1 F strain or its derivatives at an MOI=0.1. Both cells and supernatants were harvested when cytopathic effects were observed in all cells. The cells were subjected to three freeze[thaw cycles to release the viruses. Cell debris was removed by centrifuging at 6000 × g for 20 minutes. The cleared supernatant was layered on 30% sucrose and further purified by centrifugation at 12,000 ×g for 2 hours. The viruses were resuspended in PBS and subsequently aliquoted, titrated, and stored at −80°C until use.

### Growth kinetics of viruses

Vero cells seeded in 24-well plates were infected with viruses (MOI = 0.1) for 1 hour at room temperature. After washing with PBS, the cells were further cultured in DMEM supplemented with 5% FBS. Supernatants were collected at 12, 24, 36, 48, 60 and 72 hours after infection. Viral titers were determined by plaque assay.

### Transmission of gene drive

Vero cells were initially infected with the target viruses or the gene drive viruses (total MOI=5, target: gene drive = 4:1) for 24 hours. According to the Poisson distribution, such coinfection conditions meant that a host cell had a 63% probability of being simultaneously infected by the WT strain and at least one gene drive virion. In some cases, additional coinfection with a target: gene drive=19:1 was also conducted. Progeny viruses were harvested and passaged into fresh Vero cells for new cycles of coinfection as needed. To quantify the composition of the viruses, supernatants harvested from each passage were diluted and used to infect fresh BHK cells at a low MOI (∼0.1) for 12 hours. The infected cells were stained with monoclonal antibodies against HSV1 gD (sc-21719, Santa Cruz) or gE (sc-69803, Santa Cruz) and a matching fluorescent secondary antibody (A0473, Beyotime). Samples were collected via flow cytometry (NovoCyte, Agilent). The flow cytometry results were analyzed using FlowJo™ v10.8 Software (BD Life Sciences). The ratio of viruses was calculated as the percentage of gD^+^ cells.

### NGS Amplicon sequencing and analysis

PCR fragments were amplified by specific primers with adaptors (Supp. Fig. 2) and sequenced with a read length of 150 nt from both sides with the Illumina MiSeq PE150 platform (Tsingke). The paired-end sequencing raw data were first checked for quality by FastQC v0.11.9 (https://www.bioinformatics.babraham.ac.uk/projects/fastqc/) and subsequently analyzed with the commercial software Geneoious Prime 2023.2.1 (https://www.geneious.com). The reference sequences were provided in Supp. Fig. 1. The data were trimmed and mapped to reference sequences. After removing primer dimers, a neighbor-joining tree was constructed for all the mapped reads using the Tamura–Nei model. The mutation rate at each nucleotide of the reference sequences was determined from the phylogenetic tree. Indels were defined as small indels (<50 bp) or large indels (50≥[bp).

### Statistics and reproducibility

The percentage, mean, CI and standard deviation were calculated using GraphPad Prism 9.5. The sample size was not predetermined, and no data were excluded from the analyses. The experiments were not randomized, and the investigators were not blinded to the allocation of the studies during the experiments or outcome assessment.

## Supporting information

sequences

## Data availability

The amplicon sequencing data have been deposited in the NCBI Gene Expression Omnibus database (BioProject ID: PRJNA1077879). Request for reagents will be fulfilled by the corresponding author H. D after receiving the appropriate MTA.

## Acknowledgments

We would like to thank Michael Caligiuri (City of Hope, USA) for his support and guidance. This study was supported by startup funds from Southern Medical University and the Natural Science Foundation of Guangdong Province (2022A1515010421). H. D is a Pearl River Young Scholar.

## Contributions

H.D. and T.Z. conceived and designed the study; Q.Y., Z.L. and K.L. performed the experiments with the help of X.Z.; G.L. provided critical reagents; H.D. and T.Z. analyzed the data and wrote the manuscript.

