## Supplementary material for "Un1Cas12f1 and Cas9 gene drive in HSV1: viruses that ‘infect’ viruses": Supp fig

### Slide 1
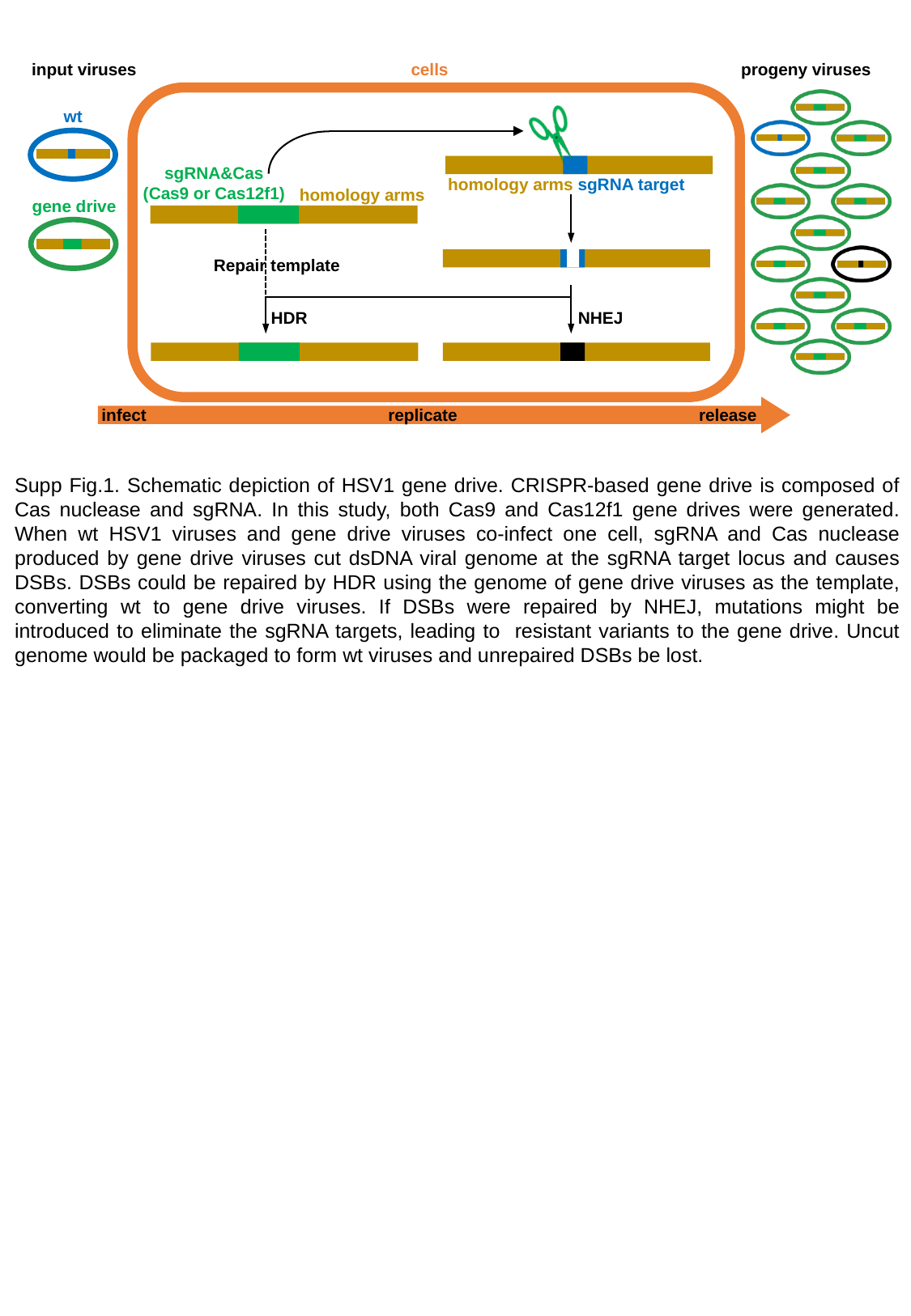

input viruses cells progeny viruses
wt
sgRNA&Cas
(Cas9 or Cas12f1)
homology arms sgRNA target
homology arms
gene drive
HDR
NHEJ
infect replicate release
Repair template
Supp Fig.1. Schematic depiction of HSV1 gene drive. CRISPR-based gene drive is composed of Cas nuclease and sgRNA. In this study, both Cas9 and Cas12f1 gene drives were generated. When wt HSV1 viruses and gene drive viruses co-infect one cell, sgRNA and Cas nuclease produced by gene drive viruses cut dsDNA viral genome at the sgRNA target locus and causes DSBs. DSBs could be repaired by HDR using the genome of gene drive viruses as the template, converting wt to gene drive viruses. If DSBs were repaired by NHEJ, mutations might be introduced to eliminate the sgRNA targets, leading to resistant variants to the gene drive. Uncut genome would be packaged to form wt viruses and unrepaired DSBs be lost.

### Slide 2
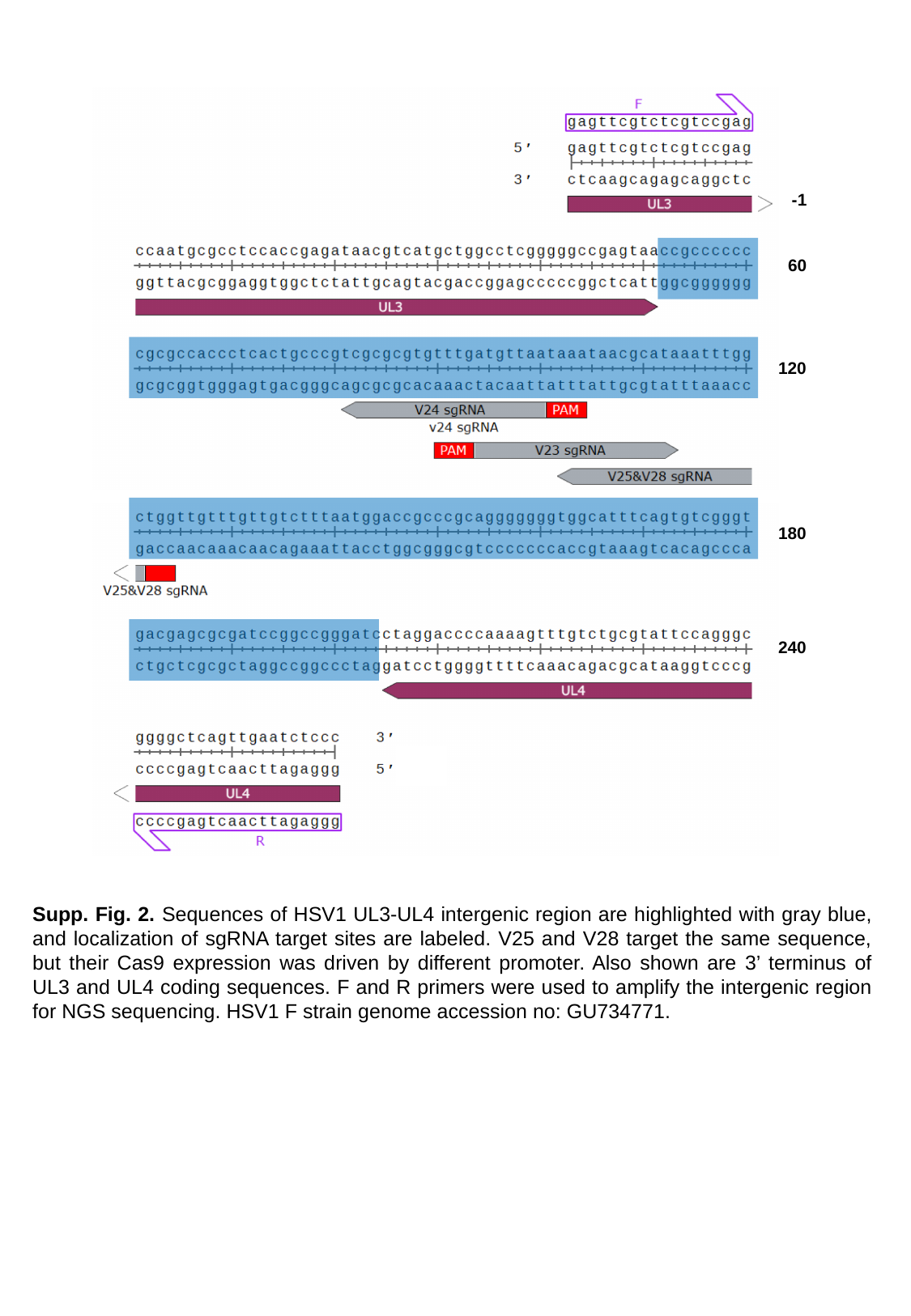

-1
60
120
180
240
Supp. Fig. 2. Sequences of HSV1 UL3-UL4 intergenic region are highlighted with gray blue, and localization of sgRNA target sites are labeled. V25 and V28 target the same sequence, but their Cas9 expression was driven by different promoter. Also shown are 3’ terminus of UL3 and UL4 coding sequences. F and R primers were used to amplify the intergenic region for NGS sequencing. HSV1 F strain genome accession no: GU734771.

### Slide 3
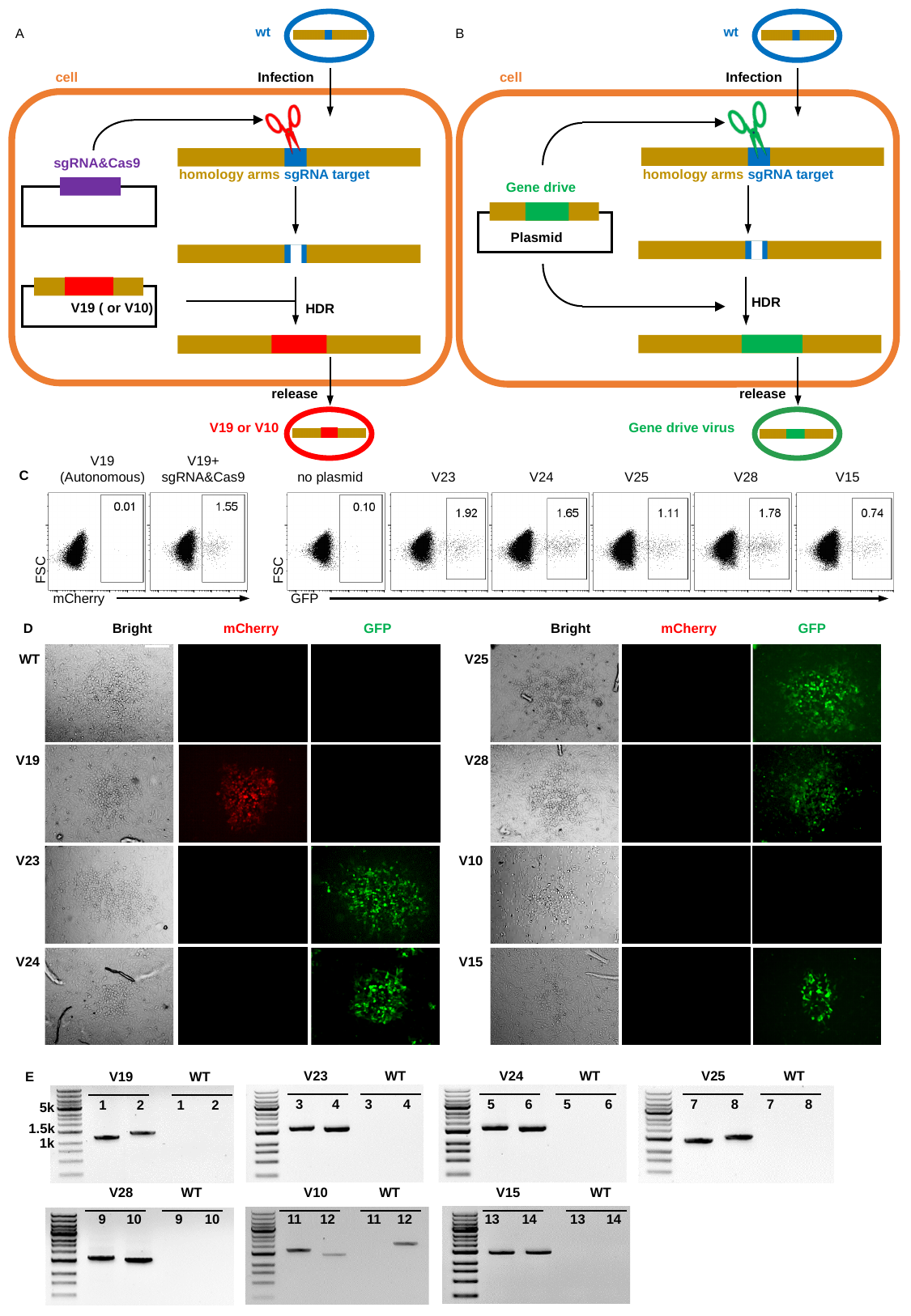

wt
wt
A
B
cell
 Infection
cell
 Infection
sgRNA&Cas9
homology arms sgRNA target
homology arms sgRNA target
Gene drive
Plasmid
HDR
V19 ( or V10)
HDR
 release
 release
V19 or V10
Gene drive virus
V19
(Autonomous)
V19+
sgRNA&Cas9
C
no plasmid
V23
V24
V25
V28
V15
0.03
FSC
FSC
mCherry
GFP
D
Bright
mCherry
GFP
Bright
mCherry
GFP
WT
V25
V19
V28
V23
V10
V24
V15
V23
WT
3
4
3
4
V24
WT
5
6
5
6
V25
WT
7
8
7
8
V19
WT
1
2
1
2
5k
1.5k
1k
V28
WT
V15
WT
V10
WT
9
10
9
10
11
12
11
12
13
14
13
14
E

### Slide 4
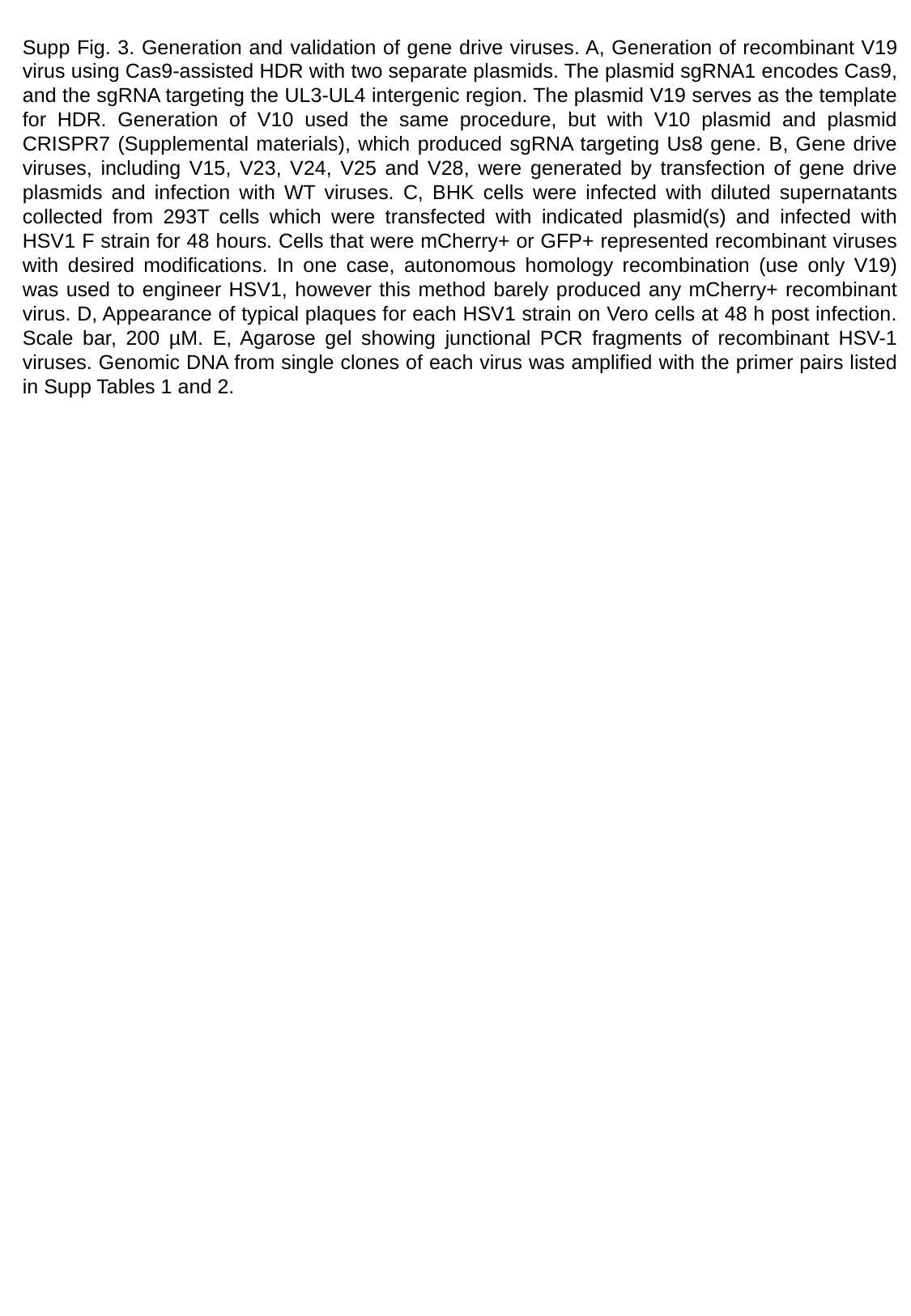

Supp Fig. 3. Generation and validation of gene drive viruses. A, Generation of recombinant V19 virus using Cas9-assisted HDR with two separate plasmids. The plasmid sgRNA1 encodes Cas9, and the sgRNA targeting the UL3-UL4 intergenic region. The plasmid V19 serves as the template for HDR. Generation of V10 used the same procedure, but with V10 plasmid and plasmid CRISPR7 (Supplemental materials), which produced sgRNA targeting Us8 gene. B, Gene drive viruses, including V15, V23, V24, V25 and V28, were generated by transfection of gene drive plasmids and infection with WT viruses. C, BHK cells were infected with diluted supernatants collected from 293T cells which were transfected with indicated plasmid(s) and infected with HSV1 F strain for 48 hours. Cells that were mCherry+ or GFP+ represented recombinant viruses with desired modifications. In one case, autonomous homology recombination (use only V19) was used to engineer HSV1, however this method barely produced any mCherry+ recombinant virus. D, Appearance of typical plaques for each HSV1 strain on Vero cells at 48 h post infection. Scale bar, 200 µM. E, Agarose gel showing junctional PCR fragments of recombinant HSV-1 viruses. Genomic DNA from single clones of each virus was amplified with the primer pairs listed in Supp Tables 1 and 2.

### Slide 5
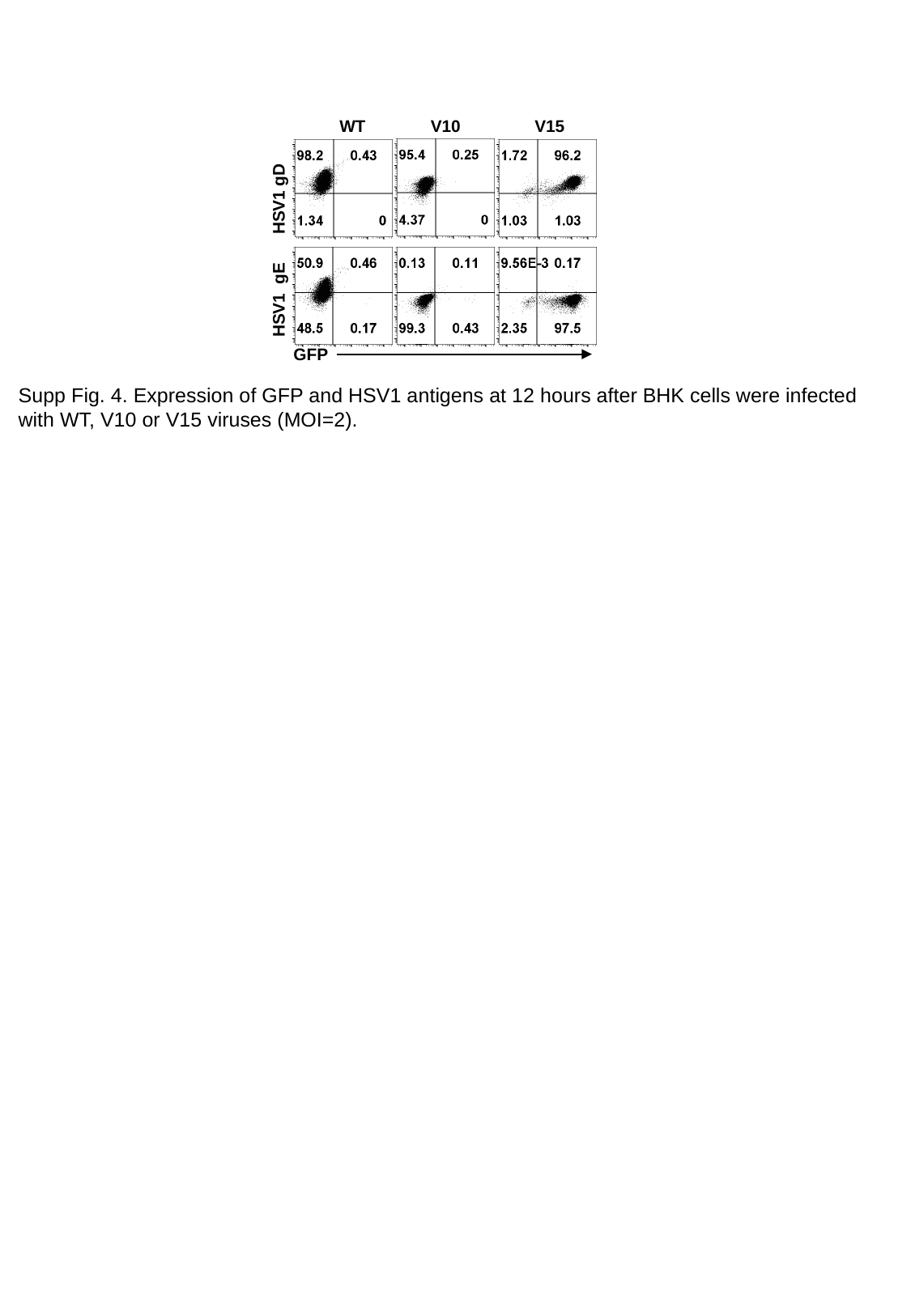

WT
V10
V15
HSV1 gD
HSV1 gE
GFP
Supp Fig. 4. Expression of GFP and HSV1 antigens at 12 hours after BHK cells were infected with WT, V10 or V15 viruses (MOI=2).

### Slide 6
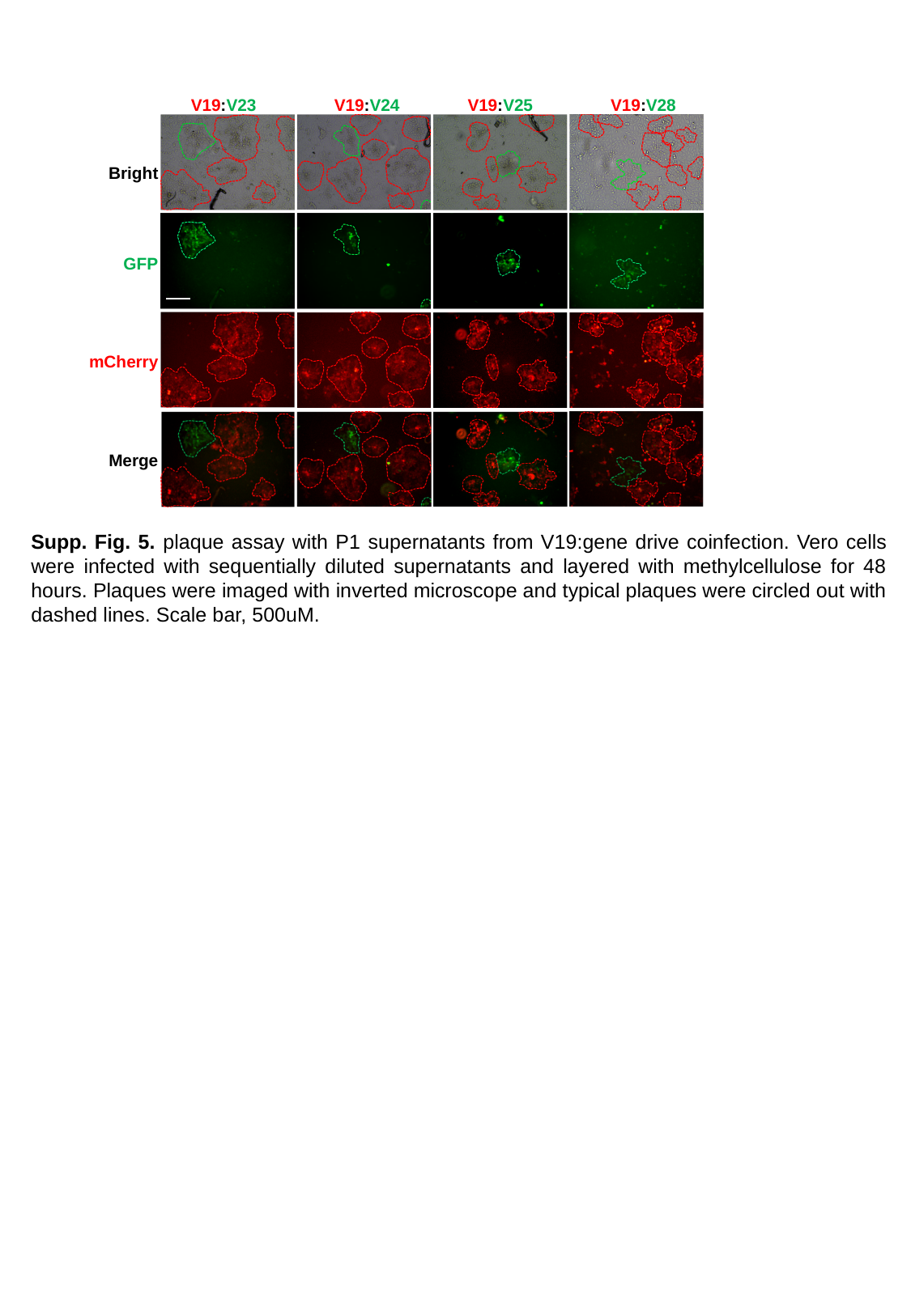

V19:V23
V19:V24
V19:V25
V19:V28
Bright
GFP
mCherry
Merge
Supp. Fig. 5. plaque assay with P1 supernatants from V19:gene drive coinfection. Vero cells were infected with sequentially diluted supernatants and layered with methylcellulose for 48 hours. Plaques were imaged with inverted microscope and typical plaques were circled out with dashed lines. Scale bar, 500uM.

### Slide 7
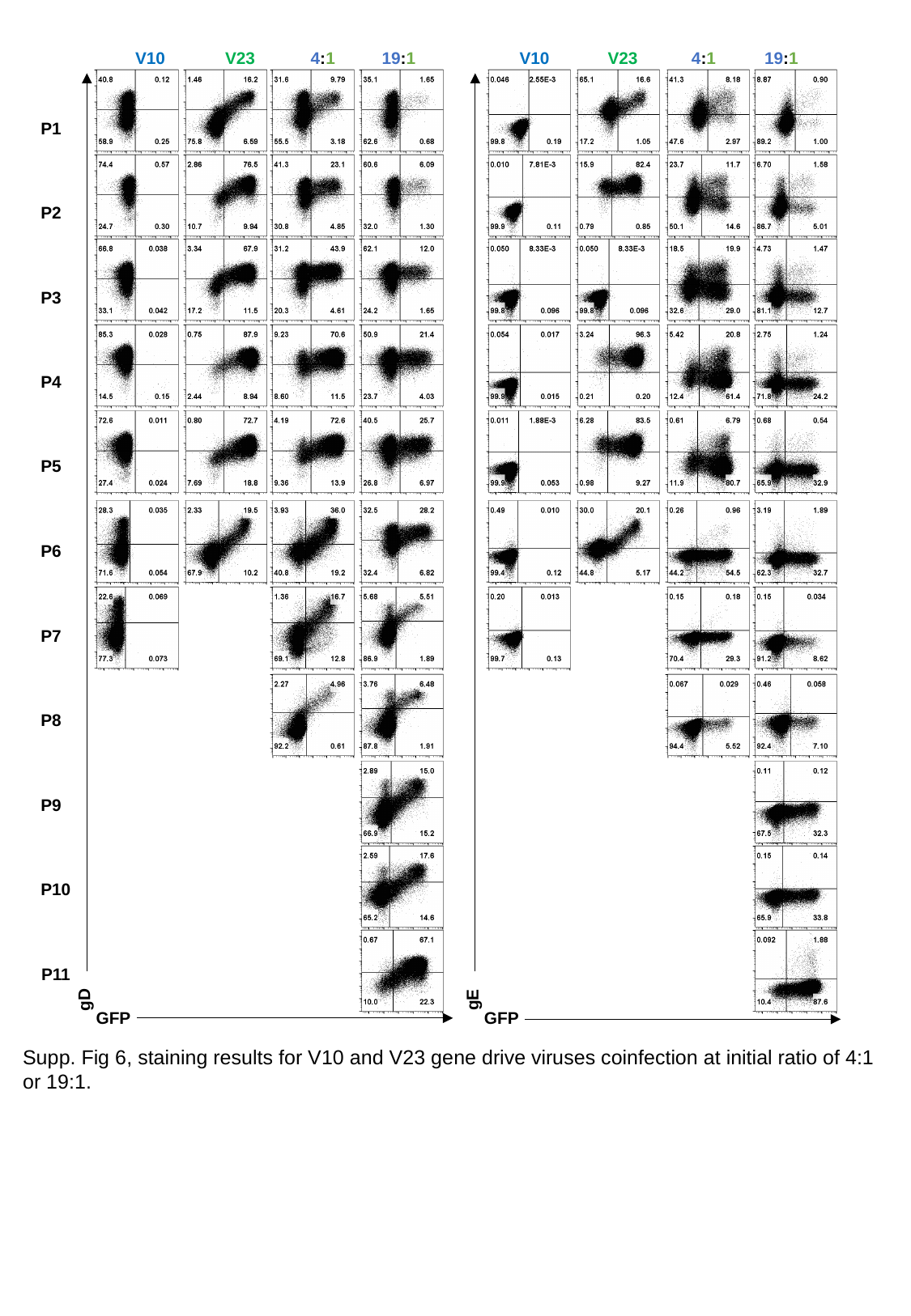

V10
V23
4:1
19:1
V10
V23
4:1
19:1
P1
P2
P3
P4
P5
P6
P7
P8
P9
P10
P11
gD
gE
GFP
GFP
Supp. Fig 6, staining results for V10 and V23 gene drive viruses coinfection at initial ratio of 4:1 or 19:1.

### Slide 8
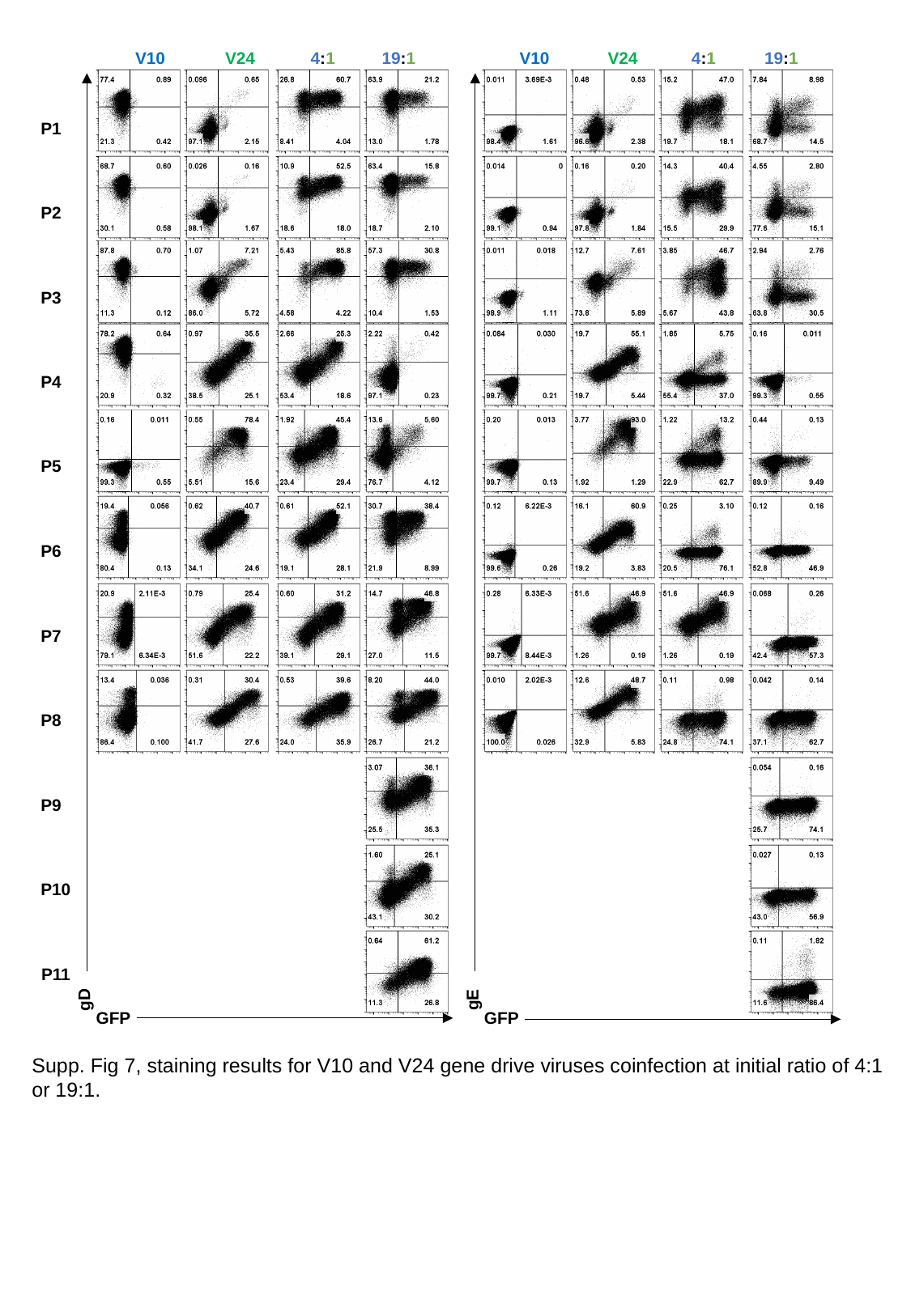

V10
V24
4:1
19:1
V10
V24
4:1
19:1
P1
P2
P3
P4
P5
P6
P7
P8
P9
P10
P11
gD
gE
GFP
GFP
Supp. Fig 7, staining results for V10 and V24 gene drive viruses coinfection at initial ratio of 4:1 or 19:1.

### Slide 9
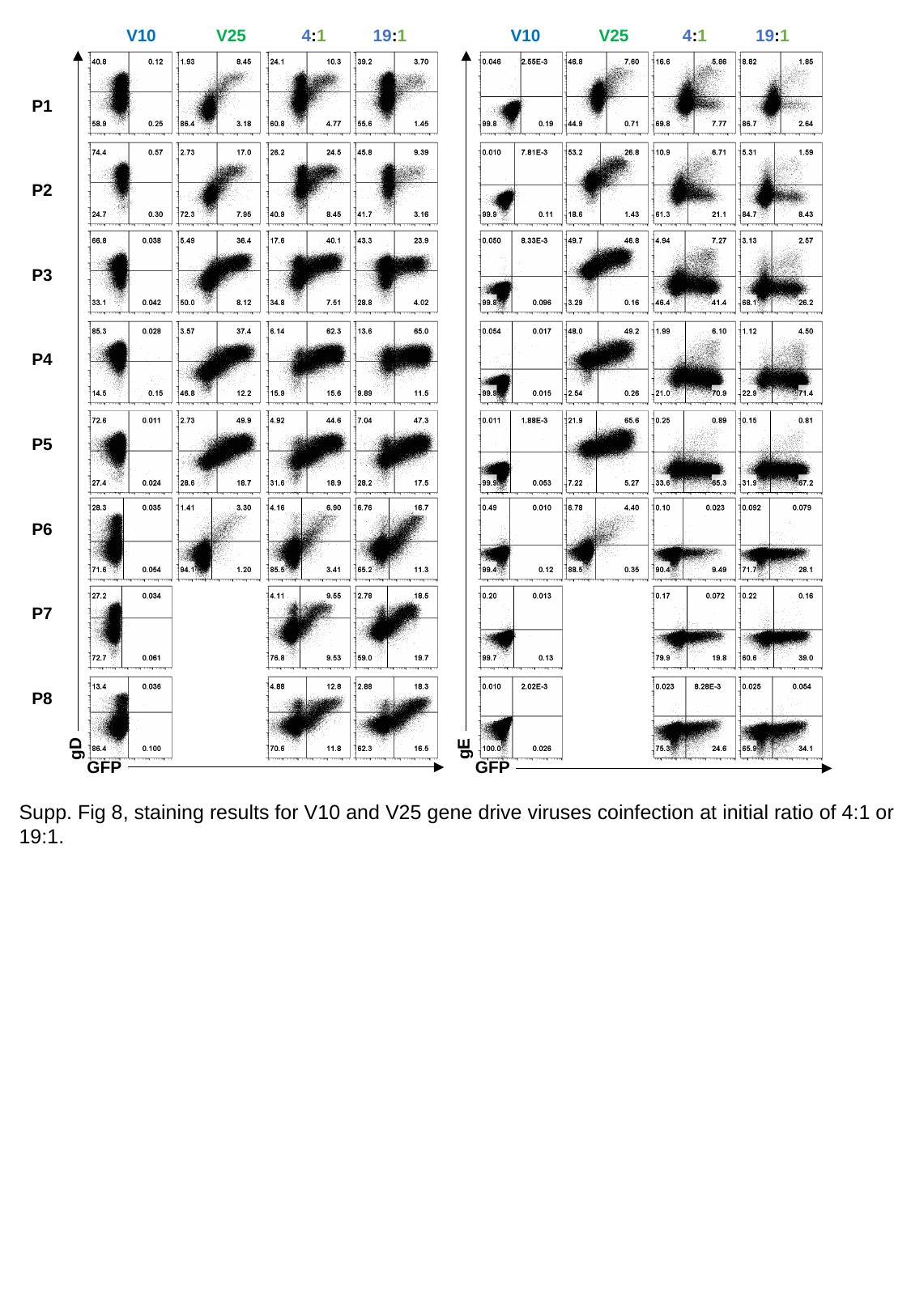

V10
V25
4:1
19:1
V10
V25
4:1
19:1
P1
P2
P3
P4
P5
P6
P7
P8
gD
gE
GFP
GFP
Supp. Fig 8, staining results for V10 and V25 gene drive viruses coinfection at initial ratio of 4:1 or 19:1.

### Slide 10
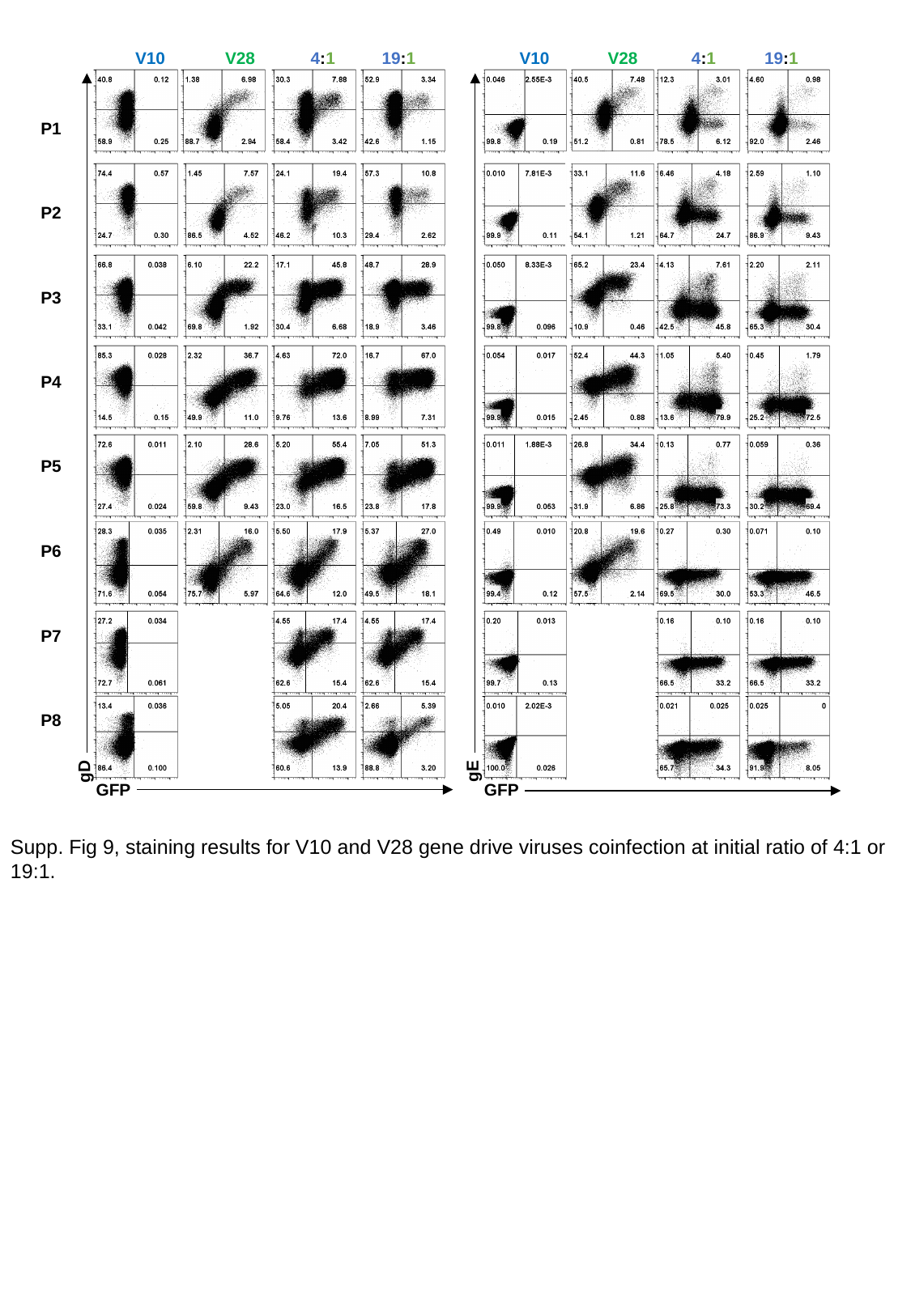

V10
V28
4:1
19:1
V10
V28
4:1
19:1
P1
P2
P3
P4
P5
P6
P7
P8
gD
gE
GFP
GFP
Supp. Fig 9, staining results for V10 and V28 gene drive viruses coinfection at initial ratio of 4:1 or 19:1.

### Slide 11
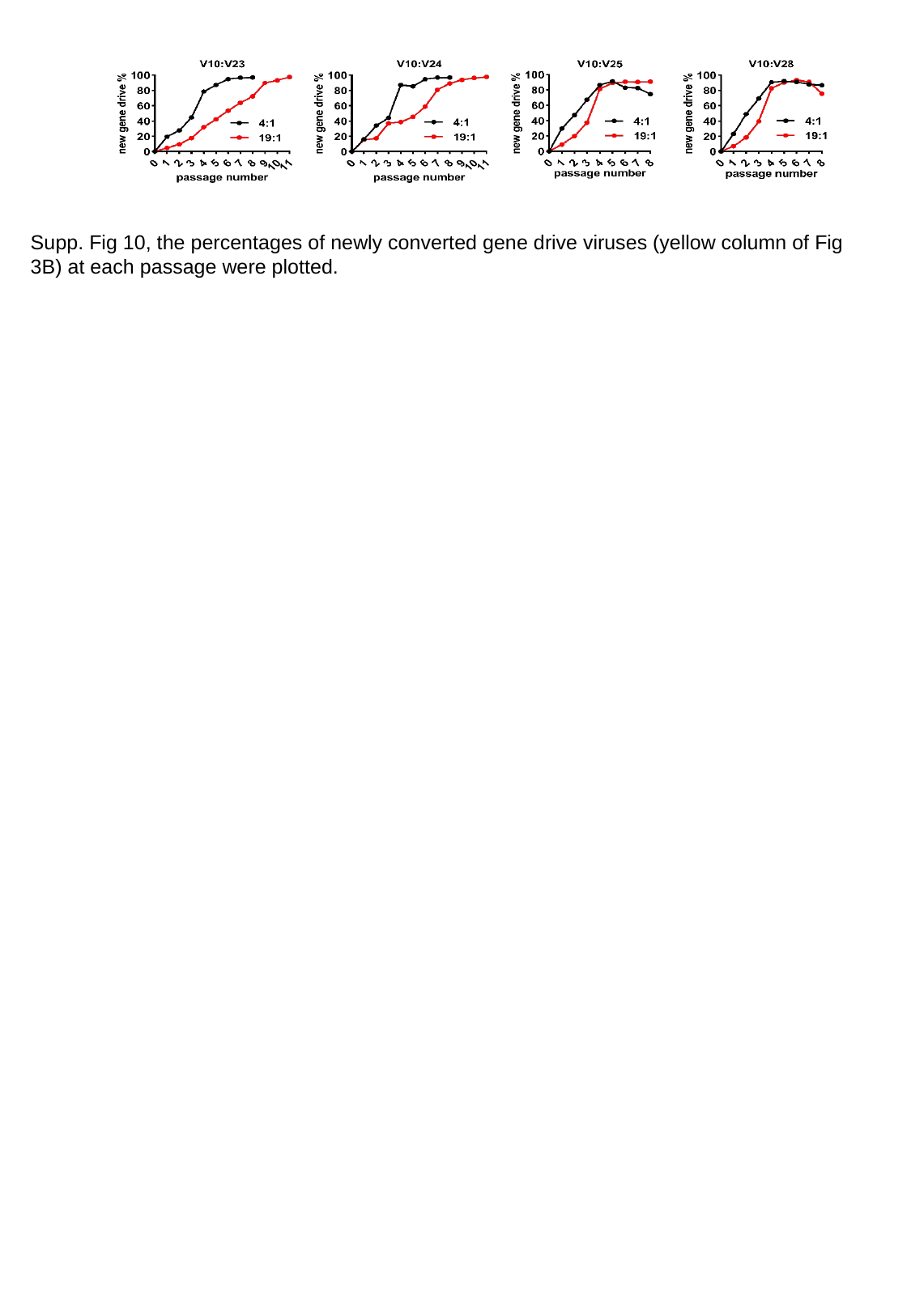

Supp. Fig 10, the percentages of newly converted gene drive viruses (yellow column of Fig 3B) at each passage were plotted.

### Slide 12
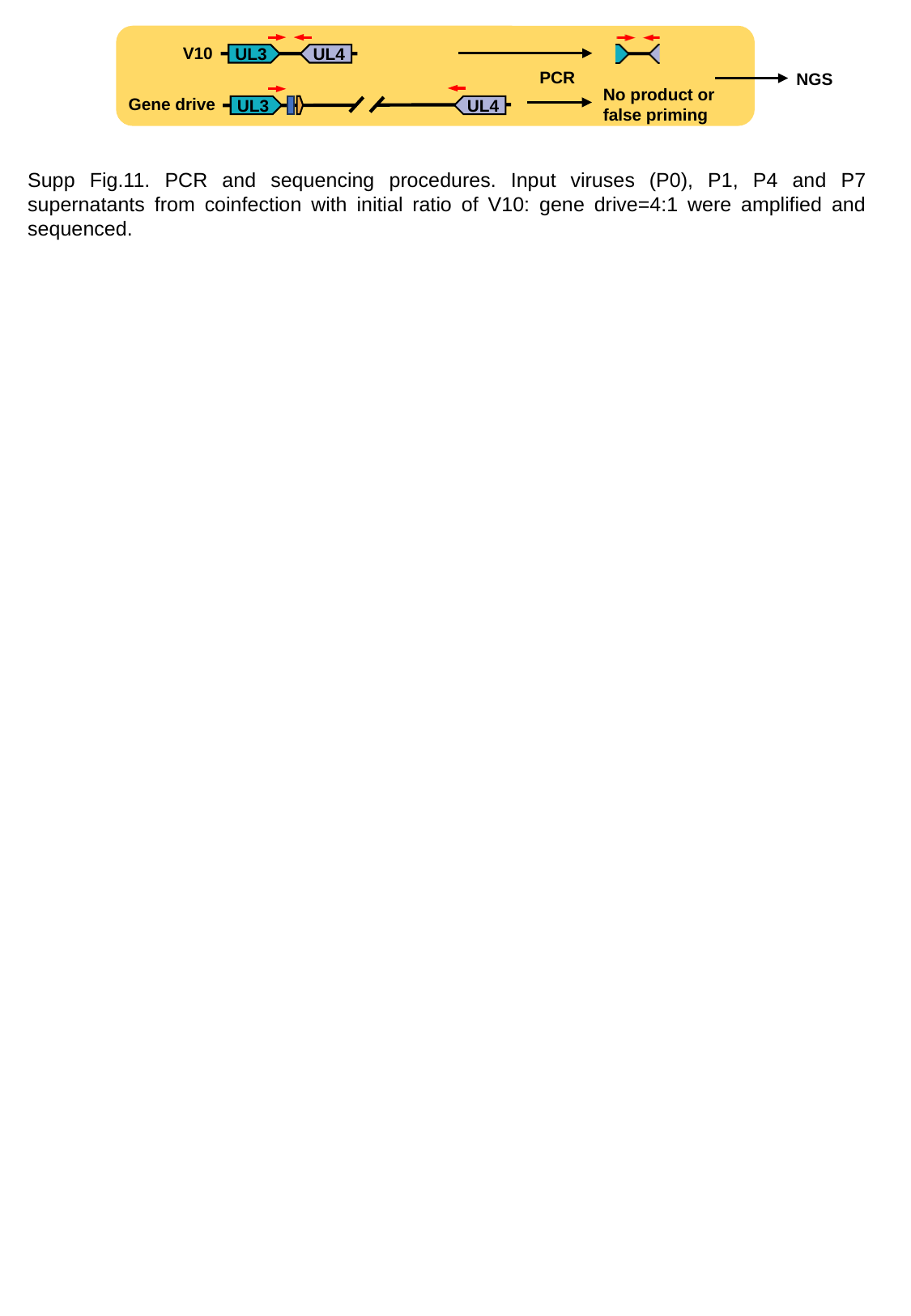

V10
UL3
UL4
PCR
NGS
No product or false priming
Gene drive
UL3
UL4
Supp Fig.11. PCR and sequencing procedures. Input viruses (P0), P1, P4 and P7 supernatants from coinfection with initial ratio of V10: gene drive=4:1 were amplified and sequenced.

### Slide 13
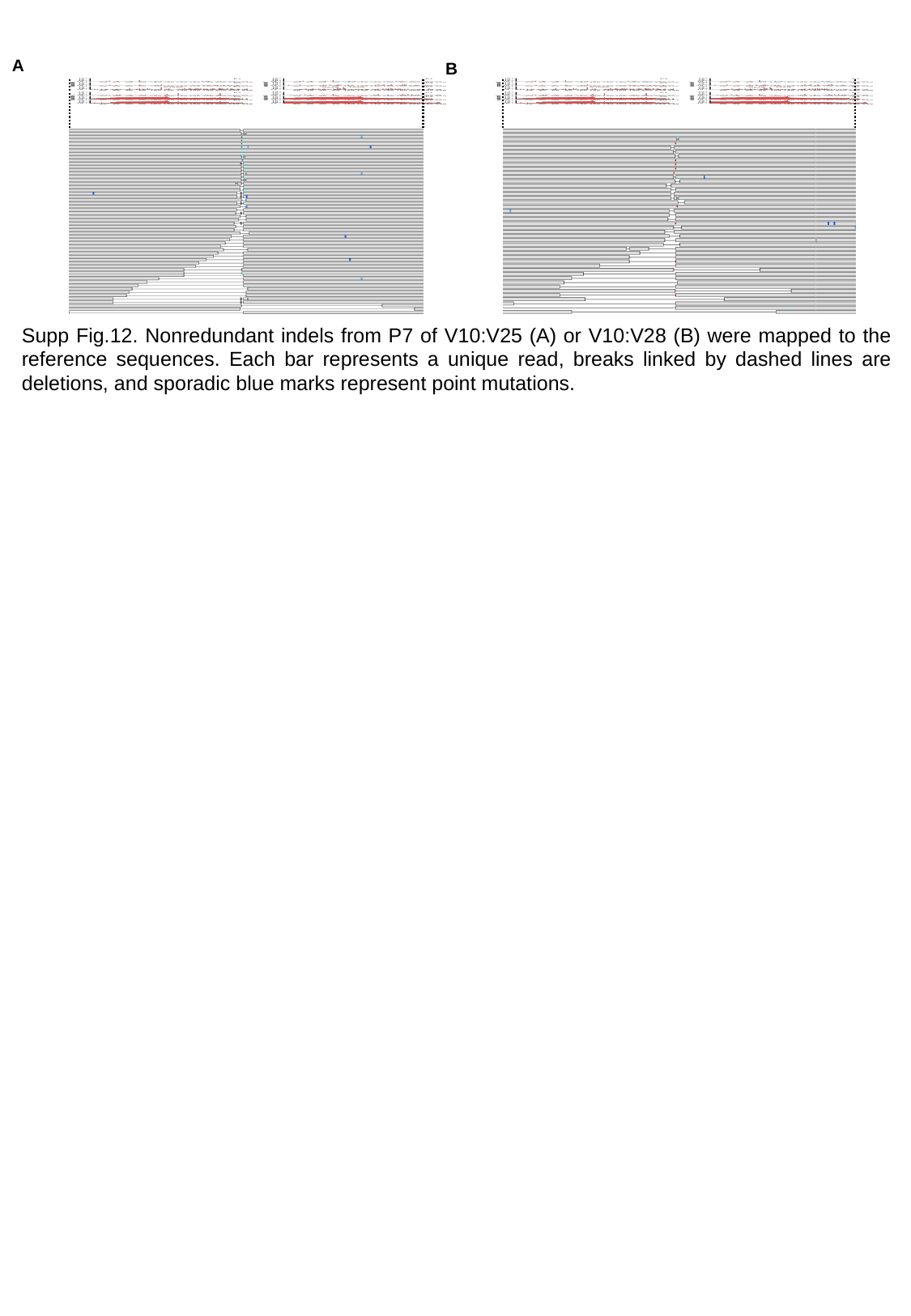

A
B
Supp Fig.12. Nonredundant indels from P7 of V10:V25 (A) or V10:V28 (B) were mapped to the reference sequences. Each bar represents a unique read, breaks linked by dashed lines are deletions, and sporadic blue marks represent point mutations.
