## Supplementary material for "Un1Cas12f1 and Cas9 gene drive in HSV1: viruses that ‘infect’ viruses": table

Supp table 1, primers’ sequences and their binding sites

| Oligo # | Sequences (5'-3') | Binding site |
| --- | --- | --- |
| HD2004 | ggtgtttcgtcctttccaca | U6 promoter |
| HD2029 | tcaacctctggattacaaaatttgtg | WPRE |
| HD2036 | actagtgaaggagagatgcgagcc | EF1 |
| HD2051 | ccaagtaggaaagtcccataa | CMV |
| HD2093 | ccggtattatttacctatatacgtg | Us7 |
| HD2096 | gcgttcgtcggaaatcgcga | Us8 |
| HD2303 | aacttgtttattgcagcttata | polyA |
| HD2355 | cgaccctgaccgtcaag | UL2 |
| HD2356 | ctcatgggatacacgtacg | UL5 |
| HD2602 | tcgtgcatgtcgtcacggtg | Us7 |
| HD2603 | gcttgtttcttccttgccac | Us9 |
| HD2619 | acccccatcgccttcgcc | EGFP |

Supp table 2, primer pairs used for amplifying recombinant HSV1 viruses

| Primer pair # | Primers | Product size |
| --- | --- | --- |
| 1 | HD2303, HD2356 | 1.3k |
| 2 | HD2355, HD2051 | 1.4k |
| 3 | HD2355, HD2036 | 1.7k |
| 4 | HD2619, HD2356 | 1.5k |
| 5 | HD2355, HD2036 | 1.7k |
| 6 | HD2619, HD2356 | 1.5k |
| 7 | HD2355, HD2004 | 1.3k |
| 8 | HD2619, HD2356 | 1.6k |
| 9 | HD2355, HD2036 | 1.7k |
| 10 | HD2619, HD2356 | 1.5k |
| 11 | HD2029, HD2603 | 1.8k |
| 12 | HD2093, HD2096 | 1.3k |
| 13 | HD2602, HD2004 | 1.5k |
| 14 | HD2619, HD2603 | 1.5k |
