## Supplementary material for "Un1Cas12f1 and Cas9 gene drive in HSV1: viruses that ‘infect’ viruses": sequences

```
Supplemental materials: plasmid sequences with annotation

plasmid： sgRNA1                6499 bp DNA
FEATURES             Location/Qualifiers
     source          1..6499
                     /mol_type="other DNA"
                     /organism="synthetic DNA construct"
     misc_feature    1..249
                     /label=U6
                     /note="Unknown feature type:regulatory  human U6 promoter"
     misc_feature    250
                     /label=Addtional G for enhancing efficacy
     misc_feature    251..270
                     /label=gRNA
     misc_feature    271..346
                     /label=gRNA scaffold
                     /note="guide RNA scaffold for the Streptococcus pyogenes
                     CRISPR/Cas9 system; gRNA scaffold"
     terminator      347..352
                     /label=polIII terminator
                     /note="Unknown feature type:regulatory  RNA polymerase III
                     transcription terminator; polIII terminator"
     misc_feature    366..651
                     /label=CMV enhancer
                     /note="Unknown feature type:regulatory  human
                     cytomegalovirus immediate early gene enhancer; CMV
                     enhancer"
     misc_feature    653..931
                     /label=CBP
                     /note="Unknown feature type:regulatory  chicken beta-actin
                     promoter"
     CDS             1180..5448
                     /label=NLS Cas9 NLS
     CDS             5503..6261
                     /label=copGFP
     polyA_signal    6292..6499
                     /label=bGH polyA
                     /note="Unknown feature type:regulatory  bovine growth
                     hormone gene polyadenylation signal; bGH polyA"
ORIGIN
        1 gagggcctat ttcccatgat tccttcatat ttgcatatac gatacaaggc tgttagagag
       61 ataattggaa ttaatttgac tgtaaacaca aagatattag tacaaaatac gtgacgtaga
      121 aagtaataat ttcttgggta gtttgcagtt ttaaaattat gttttaaaat ggactatcat
      181 atgcttaccg taacttgaaa gtatttcgat ttcttggctt tatatatctt gtggaaagga
      241 cgaaacaccg aaataacgca taaatttggc gttttagagc tagaaatagc aagttaaaat
      301 aaggctagtc cgttatcaac ttgaaaaagt ggcaccgagt cggtgctttt ttgtctagag
      361 gtacccgtta cataacttac ggtaaatggc ccgcctggct gaccgcccaa cgacccccgc
      421 ccattgacgt caatagtaac gccaataggg actttccatt gacgtcaatg ggtggagtat
      481 ttacggtaaa ctgcccactt ggcagtacat caagtgtatc atatgccaag tacgccccct
      541 attgacgtca atgacggtaa atggcccgcc tggcattgtg cccagtacat gaccttatgg
      601 gactttccta cttggcagta catctacgta ttagtcatcg ctattaccat ggtcgaggtg
      661 agccccacgt tctgcttcac tctccccatc tcccccccct ccccaccccc aattttgtat
      721 ttatttattt tttaattatt ttgtgcagcg atgggggcgg gggggggggg ggggcgcgcg
      781 ccaggcgggg cggggcgggg gcgaggggcg gggcggggcg aggcggagag gtgcggcggc
      841 agccaatcag agcggcgcgc tccgaaagtt tccttttatg gcgaggcggc ggcggcggcg
      901 gccctataaa aagcgaagcg cgcggcgggc gggagtcgct gcgcgctgcc ttcgccccgt
      961 gccccgctcc gccgccgcct cgcgccgccc gccccggctc tgactgaccg cgttactccc
     1021 acaggtgagc gggcgggacg gcccttctcc tccgggctgt aattagctga gcaagaggta
     1081 agggtttaag ggatggttgg ttggtggggt attaatgttt aattacctgg agcacctgcc
     1141 tgaaatcact ttttttcagg ttggaccggt gccaccatga tggactataa ggaccacgac
     1201 ggagactaca aggatcatga tattgattac aaagacgatg acgataagat ggccccaaag
     1261 aagaagcgga aggtcggtat ccacggagtc ccagcagccg acaagaagta cagcatcggc
     1321 ctggacatcg gcaccaactc tgtgggctgg gccgtgatca ccgacgagta caaggtgccc
     1381 agcaagaaat tcaaggtgct gggcaacacc gaccggcaca gcatcaagaa gaacctgatc
     1441 ggagccctgc tgttcgacag cggcgaaaca gccgaggcca cccggctgaa gagaaccgcc
     1501 agaagaagat acaccagacg gaagaaccgg atctgctatc tgcaagagat cttcagcaac
     1561 gagatggcca aggtggacga cagcttcttc cacagactgg aagagtcctt cctggtggaa
     1621 gaggataaga agcacgagcg gcaccccatc ttcggcaaca tcgtggacga ggtggcctac
     1681 cacgagaagt accccaccat ctaccacctg agaaagaaac tggtggacag caccgacaag
     1741 gccgacctgc ggctgatcta tctggccctg gcccacatga tcaagttccg gggccacttc
     1801 ctgatcgagg gcgacctgaa ccccgacaac agcgacgtgg acaagctgtt catccagctg
     1861 gtgcagacct acaaccagct gttcgaggaa aaccccatca acgccagcgg cgtggacgcc
     1921 aaggccatcc tgtctgccag actgagcaag agcagacggc tggaaaatct gatcgcccag
     1981 ctgcccggcg agaagaagaa tggcctgttc ggaaacctga ttgccctgag cctgggcctg
     2041 acccccaact tcaagagcaa cttcgacctg gccgaggatg ccaaactgca gctgagcaag
     2101 gacacctacg acgacgacct ggacaacctg ctggcccaga tcggcgacca gtacgccgac
     2161 ctgtttctgg ccgccaagaa cctgtccgac gccatcctgc tgagcgacat cctgagagtg
     2221 aacaccgaga tcaccaaggc ccccctgagc gcctctatga tcaagagata cgacgagcac
     2281 caccaggacc tgaccctgct gaaagctctc gtgcggcagc agctgcctga gaagtacaaa
     2341 gagattttct tcgaccagag caagaacggc tacgccggct acattgacgg cggagccagc
     2401 caggaagagt tctacaagtt catcaagccc atcctggaaa agatggacgg caccgaggaa
     2461 ctgctcgtga agctgaacag agaggacctg ctgcggaagc agcggacctt cgacaacggc
     2521 agcatccccc accagatcca cctgggagag ctgcacgcca ttctgcggcg gcaggaagat
     2581 ttttacccat tcctgaagga caaccgggaa aagatcgaga agatcctgac cttccgcatc
     2641 ccctactacg tgggccctct ggccagggga aacagcagat tcgcctggat gaccagaaag
     2701 agcgaggaaa ccatcacccc ctggaacttc gaggaagtgg tggacaaggg cgcttccgcc
     2761 cagagcttca tcgagcggat gaccaacttc gataagaacc tgcccaacga gaaggtgctg
     2821 cccaagcaca gcctgctgta cgagtacttc accgtgtata acgagctgac caaagtgaaa
     2881 tacgtgaccg agggaatgag aaagcccgcc ttcctgagcg gcgagcagaa aaaggccatc
     2941 gtggacctgc tgttcaagac caaccggaaa gtgaccgtga agcagctgaa agaggactac
     3001 ttcaagaaaa tcgagtgctt cgactccgtg gaaatctccg gcgtggaaga tcggttcaac
     3061 gcctccctgg gcacatacca cgatctgctg aaaattatca aggacaagga cttcctggac
     3121 aatgaggaaa acgaggacat tctggaagat atcgtgctga ccctgacact gtttgaggac
     3181 agagagatga tcgaggaacg gctgaaaacc tatgcccacc tgttcgacga caaagtgatg
     3241 aagcagctga agcggcggag atacaccggc tggggcaggc tgagccggaa gctgatcaac
     3301 ggcatccggg acaagcagtc cggcaagaca atcctggatt tcctgaagtc cgacggcttc
     3361 gccaacagaa acttcatgca gctgatccac gacgacagcc tgacctttaa agaggacatc
     3421 cagaaagccc aggtgtccgg ccagggcgat agcctgcacg agcacattgc caatctggcc
     3481 ggcagccccg ccattaagaa gggcatcctg cagacagtga aggtggtgga cgagctcgtg
     3541 aaagtgatgg gccggcacaa gcccgagaac atcgtgatcg aaatggccag agagaaccag
     3601 accacccaga agggacagaa gaacagccgc gagagaatga agcggatcga agagggcatc
     3661 aaagagctgg gcagccagat cctgaaagaa caccccgtgg aaaacaccca gctgcagaac
     3721 gagaagctgt acctgtacta cctgcagaat gggcgggata tgtacgtgga ccaggaactg
     3781 gacatcaacc ggctgtccga ctacgatgtg gaccatatcg tgcctcagag ctttctgaag
     3841 gacgactcca tcgacaacaa ggtgctgacc agaagcgaca agaaccgggg caagagcgac
     3901 aacgtgccct ccgaagaggt cgtgaagaag atgaagaact actggcggca gctgctgaac
     3961 gccaagctga ttacccagag aaagttcgac aatctgacca aggccgagag aggcggcctg
     4021 agcgaactgg ataaggccgg cttcatcaag agacagctgg tggaaacccg gcagatcaca
     4081 aagcacgtgg cacagatcct ggactcccgg atgaacacta agtacgacga gaatgacaag
     4141 ctgatccggg aagtgaaagt gatcaccctg aagtccaagc tggtgtccga tttccggaag
     4201 gatttccagt tttacaaagt gcgcgagatc aacaactacc accacgccca cgacgcctac
     4261 ctgaacgccg tcgtgggaac cgccctgatc aaaaagtacc ctaagctgga aagcgagttc
     4321 gtgtacggcg actacaaggt gtacgacgtg cggaagatga tcgccaagag cgagcaggaa
     4381 atcggcaagg ctaccgccaa gtacttcttc tacagcaaca tcatgaactt tttcaagacc
     4441 gagattaccc tggccaacgg cgagatccgg aagcggcctc tgatcgagac aaacggcgaa
     4501 accggggaga tcgtgtggga taagggccgg gattttgcca ccgtgcggaa agtgctgagc
     4561 atgccccaag tgaatatcgt gaaaaagacc gaggtgcaga caggcggctt cagcaaagag
     4621 tctatcctgc ccaagaggaa cagcgataag ctgatcgcca gaaagaagga ctgggaccct
     4681 aagaagtacg gcggcttcga cagccccacc gtggcctatt ctgtgctggt ggtggccaaa
     4741 gtggaaaagg gcaagtccaa gaaactgaag agtgtgaaag agctgctggg gatcaccatc
     4801 atggaaagaa gcagcttcga gaagaatccc atcgactttc tggaagccaa gggctacaaa
     4861 gaagtgaaaa aggacctgat catcaagctg cctaagtact ccctgttcga gctggaaaac
     4921 ggccggaaga gaatgctggc ctctgccggc gaactgcaga agggaaacga actggccctg
     4981 ccctccaaat atgtgaactt cctgtacctg gccagccact atgagaagct gaagggctcc
     5041 cccgaggata atgagcagaa acagctgttt gtggaacagc acaagcacta cctggacgag
     5101 atcatcgagc agatcagcga gttctccaag agagtgatcc tggccgacgc taatctggac
     5161 aaagtgctgt ccgcctacaa caagcaccgg gataagccca tcagagagca ggccgagaat
     5221 atcatccacc tgtttaccct gaccaatctg ggagcccctg ccgccttcaa gtactttgac
     5281 accaccatcg accggaagag gtacaccagc accaaagagg tgctggacgc caccctgatc
     5341 caccagagca tcaccggcct gtacgagaca cggatcgacc tgtctcagct gggaggcgac
     5401 aaaaggccgg cggccacgaa aaaggccggc caggcaaaaa agaaaaagga gggcagagga
     5461 agtctactaa catgcggtga cgtggaggag aatcccggcc ctatggagag cgacgagagc
     5521 ggcctgcccg ccatggagat cgagtgccgc atcaccggca ccctgaacgg cgtggagttc
     5581 gagctggtgg gcggcggaga gggcaccccc aagcagggcc gcatgaccaa caagatgaag
     5641 agcaccaaag gcgccctgac cttcagcccc tacctgctga gccacgtgat gggctacggc
     5701 ttctaccact tcggcaccta ccccagcggc tacgagaacc ccttcctgca cgccatcaac
     5761 aacggcggct acaccaacac ccgcatcgag aagtacgagg acggcggcgt gctgcacgtg
     5821 agcttcagct accgctacga ggccggccgc gtgatcggcg acttcaaggt ggtgggcacc
     5881 ggcttccccg aggacagcgt gatcttcacc gacaagatca tccgcagcaa cgccaccgtg
     5941 gagcacctgc accccatggg cgataacgtg ctggtgggca gcttcgcccg caccttcagc
     6001 ctgcgcgacg gcggctacta cagcttcgtg gtggacagcc acatgcactt caagagcgcc
     6061 atccacccca gcatcctgca gaacgggggc cccatgttcg ccttccgccg cgtggaggag
     6121 ctgcacagca acaccgagct gggcatcgtg gagtaccagc acgccttcaa gacccccatc
     6181 gccttcgcca gatcccgcgc tcagtcgtcc aattctgccg tggacggcac cgccggaccc
     6241 ggctccaccg gatctcgcta agaattccta gagctcgctg atcagcctcg actgtgcctt
     6301 ctagttgcca gccatctgtt gtttgcccct cccccgtgcc ttccttgacc ctggaaggtg
     6361 ccactcccac tgtcctttcc taataaaatg aggaaattgc atcgcattgt ctgagtaggt
     6421 gtcattctat tctggggggt ggggtggggc aggacagcaa gggggaggat tgggaagaga
     6481 atagcaggca tgctgggga
//

plasmid： sgRNA7                6498 bp DNA
FEATURES             Location/Qualifiers
     source          1..6498
                     /mol_type="other DNA"
                     /organism="synthetic DNA construct"
     misc_feature    1..249
                     /label=U6
                     /note="Unknown feature type:regulatory  human U6 promoter"
     misc_feature    250..269
                     /label=gRNA
     misc_feature    270..345
                     /label=gRNA scaffold
                     /note="guide RNA scaffold for the Streptococcus pyogenes
                     CRISPR/Cas9 system; gRNA scaffold"
     terminator      346..351
                     /label=polIII terminator
                     /note="Unknown feature type:regulatory  RNA polymerase III
                     transcription terminator; polIII terminator"
     misc_feature    365..650
                     /label=CMV enhancer
                     /note="Unknown feature type:regulatory  human
                     cytomegalovirus immediate early gene enhancer; CMV
                     enhancer"
     misc_feature    652..930
                     /label=CBP
                     /note="Unknown feature type:regulatory  chicken beta-actin
                     promoter"
     CDS             1179..5447
                     /label=NLS Cas9 NLS
     CDS             5502..6260
                     /label=copGFP
     polyA_signal    6291..6498
                     /label=bGH polyA
                     /note="Unknown feature type:regulatory  bovine growth
                     hormone gene polyadenylation signal; bGH polyA"
ORIGIN
        1 gagggcctat ttcccatgat tccttcatat ttgcatatac gatacaaggc tgttagagag
       61 ataattggaa ttaatttgac tgtaaacaca aagatattag tacaaaatac gtgacgtaga
      121 aagtaataat ttcttgggta gtttgcagtt ttaaaattat gttttaaaat ggactatcat
      181 atgcttaccg taacttgaaa gtatttcgat ttcttggctt tatatatctt gtggaaagga
      241 cgaaacaccg accgtgcgta tggagactcg ttttagagct agaaatagca agttaaaata
      301 aggctagtcc gttatcaact tgaaaaagtg gcaccgagtc ggtgcttttt tgtctagagg
      361 tacccgttac ataacttacg gtaaatggcc cgcctggctg accgcccaac gacccccgcc
      421 cattgacgtc aatagtaacg ccaataggga ctttccattg acgtcaatgg gtggagtatt
      481 tacggtaaac tgcccacttg gcagtacatc aagtgtatca tatgccaagt acgcccccta
      541 ttgacgtcaa tgacggtaaa tggcccgcct ggcattgtgc ccagtacatg accttatggg
      601 actttcctac ttggcagtac atctacgtat tagtcatcgc tattaccatg gtcgaggtga
      661 gccccacgtt ctgcttcact ctccccatct cccccccctc cccaccccca attttgtatt
      721 tatttatttt ttaattattt tgtgcagcga tgggggcggg gggggggggg gggcgcgcgc
      781 caggcggggc ggggcggggg cgaggggcgg ggcggggcga ggcggagagg tgcggcggca
      841 gccaatcaga gcggcgcgct ccgaaagttt ccttttatgg cgaggcggcg gcggcggcgg
      901 ccctataaaa agcgaagcgc gcggcgggcg ggagtcgctg cgcgctgcct tcgccccgtg
      961 ccccgctccg ccgccgcctc gcgccgcccg ccccggctct gactgaccgc gttactccca
     1021 caggtgagcg ggcgggacgg cccttctcct ccgggctgta attagctgag caagaggtaa
     1081 gggtttaagg gatggttggt tggtggggta ttaatgttta attacctgga gcacctgcct
     1141 gaaatcactt tttttcaggt tggaccggtg ccaccatgat ggactataag gaccacgacg
     1201 gagactacaa ggatcatgat attgattaca aagacgatga cgataagatg gccccaaaga
     1261 agaagcggaa ggtcggtatc cacggagtcc cagcagccga caagaagtac agcatcggcc
     1321 tggacatcgg caccaactct gtgggctggg ccgtgatcac cgacgagtac aaggtgccca
     1381 gcaagaaatt caaggtgctg ggcaacaccg accggcacag catcaagaag aacctgatcg
     1441 gagccctgct gttcgacagc ggcgaaacag ccgaggccac ccggctgaag agaaccgcca
     1501 gaagaagata caccagacgg aagaaccgga tctgctatct gcaagagatc ttcagcaacg
     1561 agatggccaa ggtggacgac agcttcttcc acagactgga agagtccttc ctggtggaag
     1621 aggataagaa gcacgagcgg caccccatct tcggcaacat cgtggacgag gtggcctacc
     1681 acgagaagta ccccaccatc taccacctga gaaagaaact ggtggacagc accgacaagg
     1741 ccgacctgcg gctgatctat ctggccctgg cccacatgat caagttccgg ggccacttcc
     1801 tgatcgaggg cgacctgaac cccgacaaca gcgacgtgga caagctgttc atccagctgg
     1861 tgcagaccta caaccagctg ttcgaggaaa accccatcaa cgccagcggc gtggacgcca
     1921 aggccatcct gtctgccaga ctgagcaaga gcagacggct ggaaaatctg atcgcccagc
     1981 tgcccggcga gaagaagaat ggcctgttcg gaaacctgat tgccctgagc ctgggcctga
     2041 cccccaactt caagagcaac ttcgacctgg ccgaggatgc caaactgcag ctgagcaagg
     2101 acacctacga cgacgacctg gacaacctgc tggcccagat cggcgaccag tacgccgacc
     2161 tgtttctggc cgccaagaac ctgtccgacg ccatcctgct gagcgacatc ctgagagtga
     2221 acaccgagat caccaaggcc cccctgagcg cctctatgat caagagatac gacgagcacc
     2281 accaggacct gaccctgctg aaagctctcg tgcggcagca gctgcctgag aagtacaaag
     2341 agattttctt cgaccagagc aagaacggct acgccggcta cattgacggc ggagccagcc
     2401 aggaagagtt ctacaagttc atcaagccca tcctggaaaa gatggacggc accgaggaac
     2461 tgctcgtgaa gctgaacaga gaggacctgc tgcggaagca gcggaccttc gacaacggca
     2521 gcatccccca ccagatccac ctgggagagc tgcacgccat tctgcggcgg caggaagatt
     2581 tttacccatt cctgaaggac aaccgggaaa agatcgagaa gatcctgacc ttccgcatcc
     2641 cctactacgt gggccctctg gccaggggaa acagcagatt cgcctggatg accagaaaga
     2701 gcgaggaaac catcaccccc tggaacttcg aggaagtggt ggacaagggc gcttccgccc
     2761 agagcttcat cgagcggatg accaacttcg ataagaacct gcccaacgag aaggtgctgc
     2821 ccaagcacag cctgctgtac gagtacttca ccgtgtataa cgagctgacc aaagtgaaat
     2881 acgtgaccga gggaatgaga aagcccgcct tcctgagcgg cgagcagaaa aaggccatcg
     2941 tggacctgct gttcaagacc aaccggaaag tgaccgtgaa gcagctgaaa gaggactact
     3001 tcaagaaaat cgagtgcttc gactccgtgg aaatctccgg cgtggaagat cggttcaacg
     3061 cctccctggg cacataccac gatctgctga aaattatcaa ggacaaggac ttcctggaca
     3121 atgaggaaaa cgaggacatt ctggaagata tcgtgctgac cctgacactg tttgaggaca
     3181 gagagatgat cgaggaacgg ctgaaaacct atgcccacct gttcgacgac aaagtgatga
     3241 agcagctgaa gcggcggaga tacaccggct ggggcaggct gagccggaag ctgatcaacg
     3301 gcatccggga caagcagtcc ggcaagacaa tcctggattt cctgaagtcc gacggcttcg
     3361 ccaacagaaa cttcatgcag ctgatccacg acgacagcct gacctttaaa gaggacatcc
     3421 agaaagccca ggtgtccggc cagggcgata gcctgcacga gcacattgcc aatctggccg
     3481 gcagccccgc cattaagaag ggcatcctgc agacagtgaa ggtggtggac gagctcgtga
     3541 aagtgatggg ccggcacaag cccgagaaca tcgtgatcga aatggccaga gagaaccaga
     3601 ccacccagaa gggacagaag aacagccgcg agagaatgaa gcggatcgaa gagggcatca
     3661 aagagctggg cagccagatc ctgaaagaac accccgtgga aaacacccag ctgcagaacg
     3721 agaagctgta cctgtactac ctgcagaatg ggcgggatat gtacgtggac caggaactgg
     3781 acatcaaccg gctgtccgac tacgatgtgg accatatcgt gcctcagagc tttctgaagg
     3841 acgactccat cgacaacaag gtgctgacca gaagcgacaa gaaccggggc aagagcgaca
     3901 acgtgccctc cgaagaggtc gtgaagaaga tgaagaacta ctggcggcag ctgctgaacg
     3961 ccaagctgat tacccagaga aagttcgaca atctgaccaa ggccgagaga ggcggcctga
     4021 gcgaactgga taaggccggc ttcatcaaga gacagctggt ggaaacccgg cagatcacaa
     4081 agcacgtggc acagatcctg gactcccgga tgaacactaa gtacgacgag aatgacaagc
     4141 tgatccggga agtgaaagtg atcaccctga agtccaagct ggtgtccgat ttccggaagg
     4201 atttccagtt ttacaaagtg cgcgagatca acaactacca ccacgcccac gacgcctacc
     4261 tgaacgccgt cgtgggaacc gccctgatca aaaagtaccc taagctggaa agcgagttcg
     4321 tgtacggcga ctacaaggtg tacgacgtgc ggaagatgat cgccaagagc gagcaggaaa
     4381 tcggcaaggc taccgccaag tacttcttct acagcaacat catgaacttt ttcaagaccg
     4441 agattaccct ggccaacggc gagatccgga agcggcctct gatcgagaca aacggcgaaa
     4501 ccggggagat cgtgtgggat aagggccggg attttgccac cgtgcggaaa gtgctgagca
     4561 tgccccaagt gaatatcgtg aaaaagaccg aggtgcagac aggcggcttc agcaaagagt
     4621 ctatcctgcc caagaggaac agcgataagc tgatcgccag aaagaaggac tgggacccta
     4681 agaagtacgg cggcttcgac agccccaccg tggcctattc tgtgctggtg gtggccaaag
     4741 tggaaaaggg caagtccaag aaactgaaga gtgtgaaaga gctgctgggg atcaccatca
     4801 tggaaagaag cagcttcgag aagaatccca tcgactttct ggaagccaag ggctacaaag
     4861 aagtgaaaaa ggacctgatc atcaagctgc ctaagtactc cctgttcgag ctggaaaacg
     4921 gccggaagag aatgctggcc tctgccggcg aactgcagaa gggaaacgaa ctggccctgc
     4981 cctccaaata tgtgaacttc ctgtacctgg ccagccacta tgagaagctg aagggctccc
     5041 ccgaggataa tgagcagaaa cagctgtttg tggaacagca caagcactac ctggacgaga
     5101 tcatcgagca gatcagcgag ttctccaaga gagtgatcct ggccgacgct aatctggaca
     5161 aagtgctgtc cgcctacaac aagcaccggg ataagcccat cagagagcag gccgagaata
     5221 tcatccacct gtttaccctg accaatctgg gagcccctgc cgccttcaag tactttgaca
     5281 ccaccatcga ccggaagagg tacaccagca ccaaagaggt gctggacgcc accctgatcc
     5341 accagagcat caccggcctg tacgagacac ggatcgacct gtctcagctg ggaggcgaca
     5401 aaaggccggc ggccacgaaa aaggccggcc aggcaaaaaa gaaaaaggag ggcagaggaa
     5461 gtctactaac atgcggtgac gtggaggaga atcccggccc tatggagagc gacgagagcg
     5521 gcctgcccgc catggagatc gagtgccgca tcaccggcac cctgaacggc gtggagttcg
     5581 agctggtggg cggcggagag ggcaccccca agcagggccg catgaccaac aagatgaaga
     5641 gcaccaaagg cgccctgacc ttcagcccct acctgctgag ccacgtgatg ggctacggct
     5701 tctaccactt cggcacctac cccagcggct acgagaaccc cttcctgcac gccatcaaca
     5761 acggcggcta caccaacacc cgcatcgaga agtacgagga cggcggcgtg ctgcacgtga
     5821 gcttcagcta ccgctacgag gccggccgcg tgatcggcga cttcaaggtg gtgggcaccg
     5881 gcttccccga ggacagcgtg atcttcaccg acaagatcat ccgcagcaac gccaccgtgg
     5941 agcacctgca ccccatgggc gataacgtgc tggtgggcag cttcgcccgc accttcagcc
     6001 tgcgcgacgg cggctactac agcttcgtgg tggacagcca catgcacttc aagagcgcca
     6061 tccaccccag catcctgcag aacgggggcc ccatgttcgc cttccgccgc gtggaggagc
     6121 tgcacagcaa caccgagctg ggcatcgtgg agtaccagca cgccttcaag acccccatcg
     6181 ccttcgccag atcccgcgct cagtcgtcca attctgccgt ggacggcacc gccggacccg
     6241 gctccaccgg atctcgctaa gaattcctag agctcgctga tcagcctcga ctgtgccttc
     6301 tagttgccag ccatctgttg tttgcccctc ccccgtgcct tccttgaccc tggaaggtgc
     6361 cactcccact gtcctttcct aataaaatga ggaaattgca tcgcattgtc tgagtaggtg
     6421 tcattctatt ctggggggtg gggtggggca ggacagcaag ggggaggatt gggaagagaa
     6481 tagcaggcat gctgggga
//

plasmid： V10(Us8-)                4292 bp DNA
FEATURES             Location/Qualifiers
     source          1..4292
                     /mol_type="other DNA"
                     /organism="synthetic DNA construct"
     source          3399..4292
                     /label=Source_1
                     /note="parental virus stock; acronym: HHV-1; acronym:
                     HSV-1"
                     /organism="Human herpesvirus 1"
     CDS             9..863
                     /codon_start=1
                     /label=Us7
                     /translation="LCRATDSTHSPAYPTLELNLAQQPLLRVRRATRDYAGVYVLRVWV
                     GDAPNASLFVLGMAIAAEGTLAYNGSAHGSCDPKLLPYSAPRLAPASVYQPAPNPASTP
                     STTTSTPSTTTSTPSTTIPAPQASTTPFPTGDPKPQPHGVNHEPPSNATRATRDSRYAL
                     TVTQIIQIAIPASIIALVFLGSCICFIHRCQRRYRRSRRPIYNPQIPTGISCAVNEAAM
                     ARLGAELKSHPSTPPKSRRRSSRTPMPSLTAIAEESEPAGAAGLPTPPVDPTTSTPTPP
                     LLV"
     misc_feature    1149..1182
                     /label=frt
     protein_bind    1149..1182
                     /label=FRT (minimal)
                     /bound_moiety="FLP recombinase from the Saccharomyces
                     cerevisiae 2u plasmid"
                     /note="supports FLP-mediated excision but not integration
                     (Turan and Bode, 2011)"
     misc_feature    1183..1728
                     /label=EF1
     promoter        1215..1426
                     /label=EF-1-alpha core promoter
                     /note="core promoter for human elongation factor
                     EF-1-alpha"
     LTR             1439..1707
                     /label=5' LTR (truncated)
                     /note="truncated 5' long terminal repeat (LTR) from human
                     T-cell leukemia virus (HTLV) type 1"
     regulatory      1736..1745
                     /label=Kozak sequence
                     /note="vertebrate consensus sequence for strong initiation
                     of translation (Kozak, 1987)"
                     /regulatory_class="other"
     CDS             1742..2497
                     /label=copGFP
     CDS             1766..2428
                     /codon_start=1
                     /product="green fluorescent protein 2 from Pontellina
                     plumata, also known as ppluGFP2 (Shagin et al., 2004)"
                     /label=CopGFP
                     /translation="PAMEIECRITGTLNGVEFELVGGGEGTPKQGRMTNKMKSTKGALT
                     FSPYLLSHVMGYGFYHFGTYPSGYENPFLHAINNGGYTNTRIEKYEDGGVLHVSFSYRY
                     EAGRVIGDFKVVGTGFPEDSVIFTDKIIRSNATVEHLHPMGDNVLVGSFARTFSLRDGG
                     YYSFVVDSHMHFKSAIHPSILQNGGPMFAFRRVEELHSNTELGIVEYQHAFKTPIAFA"
     misc_feature    2501..2534
                     /label=frt
     protein_bind    2501..2534
                     /label=FRT (minimal)
                     /bound_moiety="FLP recombinase from the Saccharomyces
                     cerevisiae 2u plasmid"
                     /note="supports FLP-mediated excision but not integration
                     (Turan and Bode, 2011)"
     misc_feature    2547..3135
                     /label=WPRE
                     /note="woodchuck hepatitis virus posttranscriptional
                     regulatory element"
     CDS             complement(3018..3029)
                     /codon_start=1
                     /product="Factor Xa recognition and cleavage site"
                     /label=Factor Xa site
                     /translation="IEGR"
     polyA_signal    3138..3267
                     /label=sv40 polyA
     polyA_signal    3138..3259
                     /label=SV40 poly(A) signal
                     /note="SV40 polyadenylation signal"
     CDS             3399..3555
                     /codon_start=1
                     /gene="US8"
                     /label=US8
                     /translation="TNGSGFEILSPTAPSVYPRSDGHQSRRQLTTFGSGRPDRRYSQAS
                     DSSVFW"
     CDS             3404..3883
                     /codon_start=1
                     /label=Us8A
                     /translation="MDPALRSYHQRLRLYTPVAMGINLAASSQPLDPEGPIAVTPRPPI
                     RPSSGKAPHPEAPRRSPNWATAGEVDVGDELIAISDERGPPRHDRPPLATSTAPSPHPR
                     PPGYTAVVSPMALQAVDAPSLFVAWLAARWLRGASGLGAVLCGIAWYVTSIARGA"
     CDS             3973..4245
                     /codon_start=1
                     /label=Us9
                     /translation="MTSRLSDPNSSARSDMSVPLYPTASPVSVEAYYSESEDEAANDFL
                     VRMGRQQSVLRRRRRRTRCVGMVIACLLVAVLSGGFGALLMWLLR"
ORIGIN
        1 actagtccct gtgtcgcgca accgacagca ctcacagccc cgcatatccc accctggagc
       61 tgaatctggc ccaacagccg cttttgcggg tccggagggc gacgcgtgac tatgccgggg
      121 tgtacgtgtt acgcgtatgg gtcggggacg caccaaacgc cagcctgttt gtcctgggga
      181 tggccatagc cgccgaaggg actctggcgt acaacggctc ggcccatggc tcctgcgacc
      241 cgaaactgct tccgtattcg gccccgcgtc tggccccggc gagcgtatac caacccgccc
      301 ctaacccggc ctccaccccc tcgaccacca cctccacccc ctcgaccacc acctccaccc
      361 cctcgaccac catccccgct ccccaagcat cgaccacacc cttccccacg ggagacccaa
      421 aaccccaacc tcacggggtc aaccacgaac ccccatcgaa tgccacgcga gcgacccgcg
      481 actcgcgata cgcgctaacg gtgacccaga taatccagat agccatcccc gcgtccatta
      541 tagccctggt gtttctgggg agctgtattt gctttataca cagatgtcaa cgccgctacc
      601 gacgctcccg ccgcccgatt tacaaccccc agatacccac tggcatctca tgcgcggtga
      661 acgaagcggc catggcccgc ctcggagccg agctcaaatc gcatccgagc acccccccca
      721 aatcccggcg ccggtcgtca cgcacaccaa tgccctccct gacggccatc gccgaagagt
      781 cggagcccgc gggggcggct gggcttccga cgccccccgt ggaccccacg acatccaccc
      841 caacgcctcc cctgttggta taggtccacg gccactggcc gggggcacca cataaccgac
      901 cgcagtcact gagttgggaa taaaccggta ttatttacct atatacgtgt atgtccattt
      961 cttccccccc ccccccggaa accaaagaag gaaacaaaga atggatggga ggagttcagg
     1021 aagccgggga gagggcccgc ggcgcattta aggcgttgtt gtgttgactt tggctcttct
     1081 ggcgggttgg tgcggtgctg tttgttgggc tcccatttta cccgaagatc ggctgctatc
     1141 cccgggacga agttcctatt ctctagaaag tataggaact tcaaggatct gcgatcgctc
     1201 cggtgcccgt cagtgggcag agcgcacatc gcccacagtc cccgagaagt tggggggagg
     1261 ggtcggcaat tgaacgggtg cctagagaag gtggcgcggg gtaaactggg aaagtgatgt
     1321 cgtgtactgg ctccgccttt ttcccgaggg tgggggagaa ccgtatataa gtgcagtagt
     1381 cgccgtgaac gttctttttc gcaacgggtt tgccgccaga acacagctga agcttcgagg
     1441 ggctcgcatc tctccttcac gcgcccgccg ccctacctga ggccgccatc cacgccggtt
     1501 gagtcgcgtt ctgccgcctc ccgcctgtgg tgcctcctga actgcgtccg ccgtctaggt
     1561 aagtttaaag ctcaggtcga gaccgggcct ttgtccggcg ctcccttgga gcctacctag
     1621 actcagccgg ctctccacgc tttgcctgac cctgcttgct caactctacg tctttgtttc
     1681 gttttctgtt ctgcgccgtt acagatccaa gctgtgaccg gcgcctacgc tagacgccac
     1741 catggagagc gacgagagcg gcctgcccgc catggagatc gagtgccgca tcaccggcac
     1801 cctgaacggc gtggagttcg agctggtggg cggcggagag ggcaccccca agcagggccg
     1861 catgaccaac aagatgaaga gcaccaaagg cgccctgacc ttcagcccct acctgctgag
     1921 ccacgtgatg ggctacggct tctaccactt cggcacctac cccagcggct acgagaaccc
     1981 cttcctgcac gccatcaaca acggcggcta caccaacacc cgcatcgaga agtacgagga
     2041 cggcggcgtg ctgcacgtga gcttcagcta ccgctacgag gccggccgcg tgatcggcga
     2101 cttcaaggtg gtgggcaccg gcttccccga ggacagcgtg atcttcaccg acaagatcat
     2161 ccgcagcaac gccaccgtgg agcacctgca ccccatgggc gataacgtgc tggtgggcag
     2221 cttcgcccgc accttcagcc tgcgcgacgg cggctactac agcttcgtgg tggacagcca
     2281 catgcacttc aagagcgcca tccaccccag catcctgcag aacgggggcc ccatgttcgc
     2341 cttccgccgc gtggaggagc tgcacagcaa caccgagctg ggcatcgtgg agtaccagca
     2401 cgccttcaag acccccatcg ccttcgccag atcccgcgct cagtcgtcca attctgccgt
     2461 ggacggcacc gccggacccg gctccaccgg atctcgctga gaagttccta ttctctagaa
     2521 agtataggaa cttcgtcgag ttaattaatc aacctctgga ttacaaaatt tgtgaaagat
     2581 tgactggtat tcttaactat gttgctcctt ttacgctatg tggatacgct gctttaatgc
     2641 ctttgtatca tgctattgct tcccgtatgg ctttcatttt ctcctccttg tataaatcct
     2701 ggttgctgtc tctttatgag gagttgtggc ccgttgtcag gcaacgtggc gtggtgtgca
     2761 ctgtgtttgc tgacgcaacc cccactggtt ggggcattgc caccacctgt cagctccttt
     2821 ccgggacttt cgctttcccc ctccctattg ccacggcgga actcatcgcc gcctgccttg
     2881 cccgctgctg gacaggggct cggctgttgg gcactgacaa ttccgtggtg ttgtcgggga
     2941 aatcatcgtc ctttccttgg ctgctcgcct gtgttgccac ctggattctg cgcgggacgt
     3001 ccttctgcta cgtcccttcg gccctcaatc cagcggacct tccttcccgc ggcctgctgc
     3061 cggctctgcg gcctcttccg cgtcttcgcc ttcgccctca gacgagtcgg atctcccttt
     3121 gggccgcctc cccgcctaac ttgtttattg cagcttataa tggttacaaa taaagcaata
     3181 gcatcacaaa tttcacaaat aaagcatttt tttcactgca ttctagttgt ggtttgtcca
     3241 aactcatcaa tgtatcttat catgtctgcc tcgggtaagg ggcccacgta cattcgcgtg
     3301 gccgacagcg agctgtacgc ggactggagc tcggacagcg agggagaacg cgaccaggtc
     3361 ccgtggctgg cccccccgga gagacccgac tctccctcca ccaatggatc cggctttgag
     3421 atcttatcac caacggctcc gtctgtatac ccccgtagcg atgggcatca atctcgccgc
     3481 cagctcacaa cctttggatc cggaaggccc gatcgccgtt actcccaggc ctccgattcg
     3541 tccgtcttct ggtaaggcgc cccatcccga ggccccacgt cggtcgccga actgggcgac
     3601 cgccggcgag gtggacgtcg gagacgagct aatcgcgatt tccgacgaac gcggaccccc
     3661 ccgacatgac cgcccgcccc tcgccacgtc gaccgcgccc tcgccacacc cgcgaccccc
     3721 gggctacacg gccgttgtct ccccgatggc cctccaggct gtcgacgccc cctccctgtt
     3781 tgtcgcctgg ctggccgctc ggtggctccg gggggcttcc ggcctggggg ccgtcctgtg
     3841 tgggattgcg tggtatgtga cgtcaattgc ccgaggcgca taaagggccg gtggtccgcc
     3901 tagccgcagc aaattaaaaa tcgtgagtca ctgcgaccgc aacttcccac ccggagcttt
     3961 cttccggcct cgatgacgtc ccggctctcc gatcccaact cctcagcgcg atccgacatg
     4021 tccgtgccgc tttatcccac ggcctcgcca gtttcggtcg aagcctacta ctcggaaagc
     4081 gaagacgagg cggccaacga cttcctcgta cgcatgggcc gccaacagtc ggtattaagg
     4141 cgtcgacgca gacgcacccg ctgcgtcggc atggtgatcg cctgtctcct cgtggccgtt
     4201 ctgtcgggcg gatttggggc gctcctgatg tggctgctcc gctaaaagac cgcatcgaca
     4261 cgcgcgtcct tcttgtcgtc tctcttcccc cc
//

Plasmid： V15(Cas9 Us7-Us9 gene drive)        8583 bp DNA
FEATURES             Location/Qualifiers
     source          1..8583
                     /mol_type="other DNA"
                     /organism="synthetic DNA construct"
     source          7632..8583
                     /label=Source_1
                     /note="parental virus stock; acronym: HHV-1; acronym:
                     HSV-1"
                     /organism="Human herpesvirus 1"
     CDS             9..863
                     /codon_start=1
                     /label=Us7
                     /translation="LCRATDSTHSPAYPTLELNLAQQPLLRVRRATRDYAGVYVLRVWV
                     GDAPNASLFVLGMAIAAEGTLAYNGSAHGSCDPKLLPYSAPRLAPASVYQPAPNPASTP
                     STTTSTPSTTTSTPSTTIPAPQASTTPFPTGDPKPQPHGVNHEPPSNATRATRDSRYAL
                     TVTQIIQIAIPASIIALVFLGSCICFIHRCQRRYRRSRRPIYNPQIPTGISCAVNEAAM
                     ARLGAELKSHPSTPPKSRRRSSRTPMPSLTAIAEESEPAGAAGLPTPPVDPTTSTPTPP
                     LLV"
     promoter        1135..1382
                     /label=U6 promoter
                     /note="RNA polymerase III promoter for human U6 snRNA"
     misc_RNA        1403..1478
                     /label=gRNA scaffold
                     /note="guide RNA scaffold for the Streptococcus pyogenes
                     CRISPR/Cas9 system"
     enhancer        1498..1783
                     /label=CMV enhancer
                     /note="human cytomegalovirus immediate early enhancer;
                     contains an 18-bp deletion relative to the standard CMV
                     enhancer"
     promoter        1785..2063
                     /label=chicken beta-actin promoter
     intron          2064..2291
                     /label=hybrid intron
                     /note="hybrid between chicken beta-actin (CBA) and minute
                     virus of mice (MMV) introns (Gray et al., 2011)"
     CDS             2315..2380
                     /codon_start=1
                     /product="three tandem FLAG(R) epitope tags, followed by an
                     enterokinase cleavage site"
                     /label=3xFLAG
                     /translation="DYKDHDGDYKDHDIDYKDDDDK"
     CDS             2387..2407
                     /codon_start=1
                     /product="nuclear localization signal of SV40 (simian virus
                     40) large T antigen"
                     /label=SV40 NLS
                     /translation="PKKKRKV"
     CDS             2432..6532
                     /codon_start=1
                     /product="Cas9 (Csn1) endonuclease from the Streptococcus
                     pyogenes Type II CRISPR/Cas system"
                     /label=Cas9
                     /note="generates RNA-guided double strand breaks in DNA"
                     /translation="DKKYSIGLDIGTNSVGWAVITDEYKVPSKKFKVLGNTDRHSIKKN
                     LIGALLFDSGETAEATRLKRTARRRYTRRKNRICYLQEIFSNEMAKVDDSFFHRLEESF
                     LVEEDKKHERHPIFGNIVDEVAYHEKYPTIYHLRKKLVDSTDKADLRLIYLALAHMIKF
                     RGHFLIEGDLNPDNSDVDKLFIQLVQTYNQLFEENPINASGVDAKAILSARLSKSRRLE
                     NLIAQLPGEKKNGLFGNLIALSLGLTPNFKSNFDLAEDAKLQLSKDTYDDDLDNLLAQI
                     GDQYADLFLAAKNLSDAILLSDILRVNTEITKAPLSASMIKRYDEHHQDLTLLKALVRQ
                     QLPEKYKEIFFDQSKNGYAGYIDGGASQEEFYKFIKPILEKMDGTEELLVKLNREDLLR
                     KQRTFDNGSIPHQIHLGELHAILRRQEDFYPFLKDNREKIEKILTFRIPYYVGPLARGN
                     SRFAWMTRKSEETITPWNFEEVVDKGASAQSFIERMTNFDKNLPNEKVLPKHSLLYEYF
                     TVYNELTKVKYVTEGMRKPAFLSGEQKKAIVDLLFKTNRKVTVKQLKEDYFKKIECFDS
                     VEISGVEDRFNASLGTYHDLLKIIKDKDFLDNEENEDILEDIVLTLTLFEDREMIEERL
                     KTYAHLFDDKVMKQLKRRRYTGWGRLSRKLINGIRDKQSGKTILDFLKSDGFANRNFMQ
                     LIHDDSLTFKEDIQKAQVSGQGDSLHEHIANLAGSPAIKKGILQTVKVVDELVKVMGRH
                     KPENIVIEMARENQTTQKGQKNSRERMKRIEEGIKELGSQILKEHPVENTQLQNEKLYL
                     YYLQNGRDMYVDQELDINRLSDYDVDHIVPQSFLKDDSIDNKVLTRSDKNRGKSDNVPS
                     EEVVKKMKNYWRQLLNAKLITQRKFDNLTKAERGGLSELDKAGFIKRQLVETRQITKHV
                     AQILDSRMNTKYDENDKLIREVKVITLKSKLVSDFRKDFQFYKVREINNYHHAHDAYLN
                     AVVGTALIKKYPKLESEFVYGDYKVYDVRKMIAKSEQEIGKATAKYFFYSNIMNFFKTE
                     ITLANGEIRKRPLIETNGETGEIVWDKGRDFATVRKVLSMPQVNIVKKTEVQTGGFSKE
                     SILPKRNSDKLIARKKDWDPKKYGGFDSPTVAYSVLVVAKVEKGKSKKLKSVKELLGIT
                     IMERSSFEKNPIDFLEAKGYKEVKKDLIIKLPKYSLFELENGRKRMLASAGELQKGNEL
                     ALPSKYVNFLYLASHYEKLKGSPEDNEQKQLFVEQHKHYLDEIIEQISEFSKRVILADA
                     NLDKVLSAYNKHRDKPIREQAENIIHLFTLTNLGAPAAFKYFDTTIDRKRYTSTKEVLD
                     ATLIHQSITGLYETRIDLSQLGGD"
     CDS             6533..6580
                     /codon_start=1
                     /product="bipartite nuclear localization signal from
                     nucleoplasmin"
                     /label=nucleoplasmin NLS
                     /translation="KRPAATKKAGQAKKKK"
     CDS             6581..6634
                     /codon_start=1
                     /product="2A peptide from Thosea asigna virus capsid
                     protein"
                     /label=T2A
                     /note="Eukaryotic ribosomes fail to insert a peptide bond
                     between the Gly and Pro residues, yielding separate
                     polypeptides."
                     /translation="EGRGSLLTCGDVEENPGP"
     CDS             6659..7321
                     /codon_start=1
                     /product="green fluorescent protein 2 from Pontellina
                     plumata, also known as ppluGFP2 (Shagin et al., 2004)"
                     /label=CopGFP
                     /translation="PAMEIECRITGTLNGVEFELVGGGEGTPKQGRMTNKMKSTKGALT
                     FSPYLLSHVMGYGFYHFGTYPSGYENPFLHAINNGGYTNTRIEKYEDGGVLHVSFSYRY
                     EAGRVIGDFKVVGTGFPEDSVIFTDKIIRSNATVEHLHPMGDNVLVGSFARTFSLRDGG
                     YYSFVVDSHMHFKSAIHPSILQNGGPMFAFRRVEELHSNTELGIVEYQHAFKTPIAFA"
     polyA_signal    7424..7631
                     /label=bGH poly(A) signal
                     /note="bovine growth hormone polyadenylation signal"
     CDS             7632..7788
                     /codon_start=1
                     /gene="US8"
                     /label=US8
                     /translation="TNGSGFEILSPTAPSVYPRSDGHQSRRQLTTFGSGRPDRRYSQAS
                     DSSVFW"
     CDS             7637..8116
                     /gene="US8A"
                     /label=US8A
     CDS             8206..8478
                     /gene="US9"
                     /label=US9
ORIGIN
        1 actagtccct gtgtcgcgca accgacagca ctcacagccc cgcatatccc accctggagc
       61 tgaatctggc ccaacagccg cttttgcggg tccggagggc gacgcgtgac tatgccgggg
      121 tgtacgtgtt acgcgtatgg gtcggggacg caccaaacgc cagcctgttt gtcctgggga
      181 tggccatagc cgccgaaggg actctggcgt acaacggctc ggcccatggc tcctgcgacc
      241 cgaaactgct tccgtattcg gccccgcgtc tggccccggc gagcgtatac caacccgccc
      301 ctaacccggc ctccaccccc tcgaccacca cctccacccc ctcgaccacc acctccaccc
      361 cctcgaccac catccccgct ccccaagcat cgaccacacc cttccccacg ggagacccaa
      421 aaccccaacc tcacggggtc aaccacgaac ccccatcgaa tgccacgcga gcgacccgcg
      481 actcgcgata cgcgctaacg gtgacccaga taatccagat agccatcccc gcgtccatta
      541 tagccctggt gtttctgggg agctgtattt gctttataca cagatgtcaa cgccgctacc
      601 gacgctcccg ccgcccgatt tacaaccccc agatacccac tggcatctca tgcgcggtga
      661 acgaagcggc catggcccgc ctcggagccg agctcaaatc gcatccgagc acccccccca
      721 aatcccggcg ccggtcgtca cgcacaccaa tgccctccct gacggccatc gccgaagagt
      781 cggagcccgc gggggcggct gggcttccga cgccccccgt ggaccccacg acatccaccc
      841 caacgcctcc cctgttggta taggtccacg gccactggcc gggggcacca cataaccgac
      901 cgcagtcact gagttgggaa taaaccggta ttatttacct atatacgtgt atgtccattt
      961 cttccccccc ccccccggaa accaaagaag gaaacaaaga atggatggga ggagttcagg
     1021 aagccgggga gagggcccgc ggcgcattta aggcgttgtt gtgttgactt tggctcttct
     1081 ggcgggttgg tgcggtgctg tttgttgggc tcccatttta cccgaagatc ggcgagggcc
     1141 tatttcccat gattccttca tatttgcata tacgatacaa ggctgttaga gagataattg
     1201 gaattaattt gactgtaaac acaaagatat tagtacaaaa tacgtgacgt agaaagtaat
     1261 aatttcttgg gtagtttgca gttttaaaat tatgttttaa aatggactat catatgctta
     1321 ccgtaacttg aaagtatttc gatttcttgg ctttatatat cttgtggaaa ggacgaaaca
     1381 ccgacgcgtc cccacaacac tcgttttaga gctagaaata gcaagttaaa ataaggctag
     1441 tccgttatca acttgaaaaa gtggcaccga gtcggtgctt ttttgtctag aggtacccgt
     1501 tacataactt acggtaaatg gcccgcctgg ctgaccgccc aacgaccccc gcccattgac
     1561 gtcaatagta acgccaatag ggactttcca ttgacgtcaa tgggtggagt atttacggta
     1621 aactgcccac ttggcagtac atcaagtgta tcatatgcca agtacgcccc ctattgacgt
     1681 caatgacggt aaatggcccg cctggcattg tgcccagtac atgaccttat gggactttcc
     1741 tacttggcag tacatctacg tattagtcat cgctattacc atggtcgagg tgagccccac
     1801 gttctgcttc actctcccca tctccccccc ctccccaccc ccaattttgt atttatttat
     1861 tttttaatta ttttgtgcag cgatgggggc gggggggggg ggggggcgcg cgccaggcgg
     1921 ggcggggcgg gggcgagggg cggggcgggg cgaggcggag aggtgcggcg gcagccaatc
     1981 agagcggcgc gctccgaaag tttcctttta tggcgaggcg gcggcggcgg cggccctata
     2041 aaaagcgaag cgcgcggcgg gcgggagtcg ctgcgcgctg ccttcgcccc gtgccccgct
     2101 ccgccgccgc ctcgcgccgc ccgccccggc tctgactgac cgcgttactc ccacaggtga
     2161 gcgggcggga cggcccttct cctccgggct gtaattagct gagcaagagg taagggttta
     2221 agggatggtt ggttggtggg gtattaatgt ttaattacct ggagcacctg cctgaaatca
     2281 ctttttttca ggttggaccg gtgccaccat gatggactat aaggaccacg acggagacta
     2341 caaggatcat gatattgatt acaaagacga tgacgataag atggccccaa agaagaagcg
     2401 gaaggtcggt atccacggag tcccagcagc cgacaagaag tacagcatcg gcctggacat
     2461 cggcaccaac tctgtgggct gggccgtgat caccgacgag tacaaggtgc ccagcaagaa
     2521 attcaaggtg ctgggcaaca ccgaccggca cagcatcaag aagaacctga tcggagccct
     2581 gctgttcgac agcggcgaaa cagccgaggc cacccggctg aagagaaccg ccagaagaag
     2641 atacaccaga cggaagaacc ggatctgcta tctgcaagag atcttcagca acgagatggc
     2701 caaggtggac gacagcttct tccacagact ggaagagtcc ttcctggtgg aagaggataa
     2761 gaagcacgag cggcacccca tcttcggcaa catcgtggac gaggtggcct accacgagaa
     2821 gtaccccacc atctaccacc tgagaaagaa actggtggac agcaccgaca aggccgacct
     2881 gcggctgatc tatctggccc tggcccacat gatcaagttc cggggccact tcctgatcga
     2941 gggcgacctg aaccccgaca acagcgacgt ggacaagctg ttcatccagc tggtgcagac
     3001 ctacaaccag ctgttcgagg aaaaccccat caacgccagc ggcgtggacg ccaaggccat
     3061 cctgtctgcc agactgagca agagcagacg gctggaaaat ctgatcgccc agctgcccgg
     3121 cgagaagaag aatggcctgt tcggaaacct gattgccctg agcctgggcc tgacccccaa
     3181 cttcaagagc aacttcgacc tggccgagga tgccaaactg cagctgagca aggacaccta
     3241 cgacgacgac ctggacaacc tgctggccca gatcggcgac cagtacgccg acctgtttct
     3301 ggccgccaag aacctgtccg acgccatcct gctgagcgac atcctgagag tgaacaccga
     3361 gatcaccaag gcccccctga gcgcctctat gatcaagaga tacgacgagc accaccagga
     3421 cctgaccctg ctgaaagctc tcgtgcggca gcagctgcct gagaagtaca aagagatttt
     3481 cttcgaccag agcaagaacg gctacgccgg ctacattgac ggcggagcca gccaggaaga
     3541 gttctacaag ttcatcaagc ccatcctgga aaagatggac ggcaccgagg aactgctcgt
     3601 gaagctgaac agagaggacc tgctgcggaa gcagcggacc ttcgacaacg gcagcatccc
     3661 ccaccagatc cacctgggag agctgcacgc cattctgcgg cggcaggaag atttttaccc
     3721 attcctgaag gacaaccggg aaaagatcga gaagatcctg accttccgca tcccctacta
     3781 cgtgggccct ctggccaggg gaaacagcag attcgcctgg atgaccagaa agagcgagga
     3841 aaccatcacc ccctggaact tcgaggaagt ggtggacaag ggcgcttccg cccagagctt
     3901 catcgagcgg atgaccaact tcgataagaa cctgcccaac gagaaggtgc tgcccaagca
     3961 cagcctgctg tacgagtact tcaccgtgta taacgagctg accaaagtga aatacgtgac
     4021 cgagggaatg agaaagcccg ccttcctgag cggcgagcag aaaaaggcca tcgtggacct
     4081 gctgttcaag accaaccgga aagtgaccgt gaagcagctg aaagaggact acttcaagaa
     4141 aatcgagtgc ttcgactccg tggaaatctc cggcgtggaa gatcggttca acgcctccct
     4201 gggcacatac cacgatctgc tgaaaattat caaggacaag gacttcctgg acaatgagga
     4261 aaacgaggac attctggaag atatcgtgct gaccctgaca ctgtttgagg acagagagat
     4321 gatcgaggaa cggctgaaaa cctatgccca cctgttcgac gacaaagtga tgaagcagct
     4381 gaagcggcgg agatacaccg gctggggcag gctgagccgg aagctgatca acggcatccg
     4441 ggacaagcag tccggcaaga caatcctgga tttcctgaag tccgacggct tcgccaacag
     4501 aaacttcatg cagctgatcc acgacgacag cctgaccttt aaagaggaca tccagaaagc
     4561 ccaggtgtcc ggccagggcg atagcctgca cgagcacatt gccaatctgg ccggcagccc
     4621 cgccattaag aagggcatcc tgcagacagt gaaggtggtg gacgagctcg tgaaagtgat
     4681 gggccggcac aagcccgaga acatcgtgat cgaaatggcc agagagaacc agaccaccca
     4741 gaagggacag aagaacagcc gcgagagaat gaagcggatc gaagagggca tcaaagagct
     4801 gggcagccag atcctgaaag aacaccccgt ggaaaacacc cagctgcaga acgagaagct
     4861 gtacctgtac tacctgcaga atgggcggga tatgtacgtg gaccaggaac tggacatcaa
     4921 ccggctgtcc gactacgatg tggaccatat cgtgcctcag agctttctga aggacgactc
     4981 catcgacaac aaggtgctga ccagaagcga caagaaccgg ggcaagagcg acaacgtgcc
     5041 ctccgaagag gtcgtgaaga agatgaagaa ctactggcgg cagctgctga acgccaagct
     5101 gattacccag agaaagttcg acaatctgac caaggccgag agaggcggcc tgagcgaact
     5161 ggataaggcc ggcttcatca agagacagct ggtggaaacc cggcagatca caaagcacgt
     5221 ggcacagatc ctggactccc ggatgaacac taagtacgac gagaatgaca agctgatccg
     5281 ggaagtgaaa gtgatcaccc tgaagtccaa gctggtgtcc gatttccgga aggatttcca
     5341 gttttacaaa gtgcgcgaga tcaacaacta ccaccacgcc cacgacgcct acctgaacgc
     5401 cgtcgtggga accgccctga tcaaaaagta ccctaagctg gaaagcgagt tcgtgtacgg
     5461 cgactacaag gtgtacgacg tgcggaagat gatcgccaag agcgagcagg aaatcggcaa
     5521 ggctaccgcc aagtacttct tctacagcaa catcatgaac tttttcaaga ccgagattac
     5581 cctggccaac ggcgagatcc ggaagcggcc tctgatcgag acaaacggcg aaaccgggga
     5641 gatcgtgtgg gataagggcc gggattttgc caccgtgcgg aaagtgctga gcatgcccca
     5701 agtgaatatc gtgaaaaaga ccgaggtgca gacaggcggc ttcagcaaag agtctatcct
     5761 gcccaagagg aacagcgata agctgatcgc cagaaagaag gactgggacc ctaagaagta
     5821 cggcggcttc gacagcccca ccgtggccta ttctgtgctg gtggtggcca aagtggaaaa
     5881 gggcaagtcc aagaaactga agagtgtgaa agagctgctg gggatcacca tcatggaaag
     5941 aagcagcttc gagaagaatc ccatcgactt tctggaagcc aagggctaca aagaagtgaa
     6001 aaaggacctg atcatcaagc tgcctaagta ctccctgttc gagctggaaa acggccggaa
     6061 gagaatgctg gcctctgccg gcgaactgca gaagggaaac gaactggccc tgccctccaa
     6121 atatgtgaac ttcctgtacc tggccagcca ctatgagaag ctgaagggct cccccgagga
     6181 taatgagcag aaacagctgt ttgtggaaca gcacaagcac tacctggacg agatcatcga
     6241 gcagatcagc gagttctcca agagagtgat cctggccgac gctaatctgg acaaagtgct
     6301 gtccgcctac aacaagcacc gggataagcc catcagagag caggccgaga atatcatcca
     6361 cctgtttacc ctgaccaatc tgggagcccc tgccgccttc aagtactttg acaccaccat
     6421 cgaccggaag aggtacacca gcaccaaaga ggtgctggac gccaccctga tccaccagag
     6481 catcaccggc ctgtacgaga cacggatcga cctgtctcag ctgggaggcg acaaaaggcc
     6541 ggcggccacg aaaaaggccg gccaggcaaa aaagaaaaag gagggcagag gaagtctact
     6601 aacatgcggt gacgtggagg agaatcccgg ccctatggag agcgacgaga gcggcctgcc
     6661 cgccatggag atcgagtgcc gcatcaccgg caccctgaac ggcgtggagt tcgagctggt
     6721 gggcggcgga gagggcaccc ccaagcaggg ccgcatgacc aacaagatga agagcaccaa
     6781 aggcgccctg accttcagcc cctacctgct gagccacgtg atgggctacg gcttctacca
     6841 cttcggcacc taccccagcg gctacgagaa ccccttcctg cacgccatca acaacggcgg
     6901 ctacaccaac acccgcatcg agaagtacga ggacggcggc gtgctgcacg tgagcttcag
     6961 ctaccgctac gaggccggcc gcgtgatcgg cgacttcaag gtggtgggca ccggcttccc
     7021 cgaggacagc gtgatcttca ccgacaagat catccgcagc aacgccaccg tggagcacct
     7081 gcaccccatg ggcgataacg tgctggtggg cagcttcgcc cgcaccttca gcctgcgcga
     7141 cggcggctac tacagcttcg tggtggacag ccacatgcac ttcaagagcg ccatccaccc
     7201 cagcatcctg cagaacgggg gccccatgtt cgccttccgc cgcgtggagg agctgcacag
     7261 caacaccgag ctgggcatcg tggagtacca gcacgccttc aagaccccca tcgccttcgc
     7321 cagatcccgc gctcagtcgt ccaattctgc cgtggacggc accgccggac ccggctccac
     7381 cggatctcgc taagaattcc tagagctcgc tgatcagcct cgactgtgcc ttctagttgc
     7441 cagccatctg ttgtttgccc ctcccccgtg ccttccttga ccctggaagg tgccactccc
     7501 actgtccttt cctaataaaa tgaggaaatt gcatcgcatt gtctgagtag gtgtcattct
     7561 attctggggg gtggggtggg gcaggacagc aagggggagg attgggaaga gaatagcagg
     7621 catgctgggg acaccaatgg atccggcttt gagatcttat caccaacggc tccgtctgta
     7681 tacccccgta gcgatgggca tcaatctcgc cgccagctca caacctttgg atccggaagg
     7741 cccgatcgcc gttactccca ggcctccgat tcgtccgtct tctggtaagg cgccccatcc
     7801 cgaggcccca cgtcggtcgc cgaactgggc gaccgccggc gaggtggacg tcggagacga
     7861 gctaatcgcg atttccgacg aacgcggacc cccccgacat gaccgcccgc ccctcgccac
     7921 gtcgaccgcg ccctcgccac acccgcgacc cccgggctac acggccgttg tctccccgat
     7981 ggccctccag gctgtcgacg ccccctccct gtttgtcgcc tggctggccg ctcggtggct
     8041 ccggggggct tccggcctgg gggccgtcct gtgtgggatt gcgtggtatg tgacgtcaat
     8101 tgcccgaggc gcataaaggg ccggtggtcc gcctagccgc agcaaattaa aaatcgtgag
     8161 tcactgcgac cgcaacttcc cacccggagc tttcttccgg cctcgatgac gtcccggctc
     8221 tccgatccca actcctcagc gcgatccgac atgtccgtgc cgctttatcc cacggcctcg
     8281 ccagtttcgg tcgaagccta ctactcggaa agcgaagacg aggcggccaa cgacttcctc
     8341 gtacgcatgg gccgccaaca gtcggtatta aggcgtcgac gcagacgcac ccgctgcgtc
     8401 ggcatggtga tcgcctgtct cctcgtggcc gttctgtcgg gcggatttgg ggcgctcctg
     8461 atgtggctgc tccgctaaaa gaccgcatcg acacgcgcgt ccttcttgtc gtctctcttc
     8521 ccccccatca ccccgcaatt tgcacccagc ctttaactac attaaattgg gttcgattgg
     8581 caa
//

Plasmid： V19 (UL3-UL4 intergeneic mCherry)  3789 bp DNA
FEATURES             Location/Qualifiers
     source          1..3789
                     /mol_type="other DNA"
                     /organism="synthetic DNA construct"
     CDS             1..708
                     /label=UL3
     enhancer        759..1062
                     /label=Enhancer_1
                     /note="CMV enhancer human cytomegalovirus immediate early
                     enhancer"
     promoter        1063..1266
                     /label=CMV promoter
                     /note="CMV promoter human cytomegalovirus (CMV) immediate
                     early  promoter"
     CDS             1311..2018
                     /codon_start=1
                     /product="monomeric derivative of DsRed fluorescent protein
                     (Shaner et al., 2004)"
                     /label=mCherry
                     /note="mammalian codon-optimized"
                     /translation="MVSKGEEDNMAIIKEFMRFKVHMEGSVNGHEFEIEGEGEGRPYEG
                     TQTAKLKVTKGGPLPFAWDILSPQFMYGSKAYVKHPADIPDYLKLSFPEGFKWERVMNF
                     EDGGVVTVTQDSSLQDGEFIYKVKLRGTNFPSDGPVMQKKTMGWEASSERMYPEDGALK
                     GEIKQRLKLKDGGHYDAEVKTTYKAKKPVQLPGAYNVNIKLDITSHNEDYTIVEQYERA
                     EGRHSTGGMDELYK"
     polyA_signal    3000..3121
                     /label=PolyA_Signal_1
                     /note="SV40 poly(A) signal SV40 polyadenylation signal"
     CDS             complement(3190..3789)
                     /label=UL4
ORIGIN
        1 atggttaaac ctctggtctc atacgggtcg gtgatgtcgg gcgtcggggg agagggagtt
       61 ccctctgcgc ttgcgattct agcctcgtgg ggctggacgt tcgacacgcc aaaccacgag
      121 tcagggatat cgccagatac gactcccgca gattccattc ggggggccgc tgtggcctca
      181 cctgaccaac ctttacacgg gggcccggaa cgggaggcca cagcgccgtc tttctcccca
      241 acgcgcgcgg atgacggccc gccctgtacc gacgggccct acgtgacgtt tgataccctg
      301 tttatggtgt cgtcgatcga cgaattaggg cgtcgccagc tcacggacac catccgcaag
      361 gacctgcggt tgtcgctggc caagtttagc attgcgtgca ccaagacctc ctcgttttcg
      421 ggaaacgccc cgcgccacca cagacgcggg gcgttccagc gcggcacgcg ggcgccgcgc
      481 agcaacaaaa gccttcagat gtttgtgttg tgcaaacgca cccacgccgc tcgagtgcga
      541 gagcagcttc gggtcgttat tcagtcccgc aagccgcgca agtattacac gcgatcttcg
      601 gacgggcggc tctgccccgc cgtccccgtg ttcgtccacg agttcgtctc gtccgagcca
      661 atgcgcctcc accgagataa cgtcatgctg gcctcggggg ccgagtaacc gcccccccgc
      721 gccaccctca ctgcccgtcg cgcgtgtttg atgttaatcg ttacataact tacggtaaat
      781 ggcccgcctg gctgaccgcc caacgacccc cgcccattga cgtcaataat gacgtatgtt
      841 cccatagtaa cgccaatagg gactttccat tgacgtcaat gggtggagta tttacggtaa
      901 actgcccact tggcagtaca tcaagtgtat catatgccaa gtacgccccc tattgacgtc
      961 aatgacggta aatggcccgc ctggcattat gcccagtaca tgaccttatg ggactttcct
     1021 acttggcagt acatctacgt attagtcatc gctattacca tggtgatgcg gttttggcag
     1081 tacatcaatg ggcgtggata gcggtttgac tcacggggat ttccaagtct ccaccccatt
     1141 gacgtcaatg ggagtttgtt ttggcaccaa aatcaacggg actttccaaa atgtcgtaac
     1201 aactccgccc cattgacgca aatgggcggt aggcgtgtac ggtgggaggt ctatataagc
     1261 agagctggtt tagtgaaccg tcagatccgc tagcgctacc ggtcgccacc atggtgagca
     1321 agggcgagga ggataacatg gccatcatca aggagttcat gcgcttcaag gtgcacatgg
     1381 agggctccgt gaacggccac gagttcgaga tcgagggcga gggcgagggc cgcccctacg
     1441 agggcaccca gaccgccaag ctgaaggtga ccaagggtgg ccccctgccc ttcgcctggg
     1501 acatcctgtc ccctcagttc atgtacggct ccaaggccta cgtgaagcac cccgccgaca
     1561 tccccgacta cttgaagctg tccttccccg agggcttcaa gtgggagcgc gtgatgaact
     1621 tcgaggacgg cggcgtggtg accgtgaccc aggactcctc cctacaggac ggcgagttca
     1681 tctacaaggt gaagctgcgc ggcaccaact tcccctccga cggccccgta atgcagaaga
     1741 agaccatggg ctgggaggcc tcctccgagc ggatgtaccc cgaggacggc gccctgaagg
     1801 gcgagatcaa gcagaggctg aagctgaagg acggcggcca ctacgacgct gaggtcaaga
     1861 ccacctacaa ggccaagaag cccgtgcagc tgcccggcgc ctacaacgtc aacatcaagt
     1921 tggacatcac ctcccacaac gaggactaca ccatcgtgga acagtacgaa cgcgccgagg
     1981 gccgccactc caccggcggc atggacgagc tgtacaagtc cggagagggc agaggaagtc
     2041 ttctaacatg cggtgacgtg gaggagaatc ccggccctat gatgctgaag aagatcctga
     2101 agatcgagga actggacgag agagagctga tcgacatcga ggtgtccggc aaccacctgt
     2161 tctacgccaa cgacatcctc acccacaact cctcttggtt ctggaaaccc cgttataagt
     2221 gtgtcaacct gtctatcaag gacatcctgg agccggatgc cgcggaaccg gacgtgcagc
     2281 gtggcaggag cttccacttc tacgatgcca tggatgggca gatacagggc agcgtggagc
     2341 tggcagcccc aggacaggca aagatcgcag gcggggccgc ggtgtctgac agctccagca
     2401 cctcaatgaa tgtgtactcg ctgagtgtgg accctaacac ctggcagact ctgctccatg
     2461 agaggcacct gcggcagcca gaacacaaag tcctgcagca gctgcgcagc cgcggggaca
     2521 acgtgtacgt ggtgactgag gtgctgcaga cacagaagga ggtggaagtc acgcgcaccc
     2581 acaagcggga gggctcgggc cggttttccc tgcccggagc cacgtgcttg cagggtgagg
     2641 gccagggcca tctgagccag aagaagacgg tcaccatccc ctcaggcagc accctcgcat
     2701 tccgggtggc ccagctggtt attgactctg acttggacgt ccttctcttc ccggataaga
     2761 agcagaggac cttccagcca cccgcgacag gccacaagcg ttccacgagc gaaggcgcct
     2821 ggccacagct gccctctggc ctctccatga tgaggtgcct ccacaacttc ctgacagatt
     2881 gatctagata actgatcata atcagccata ccacatttgt agaggtttta cttgctttaa
     2941 aaaacctccc acacctcccc ctgaacctga aacataaaat gaatgcaatt gttgttgtta
     3001 acttgtttat tgcagcttat aatggttaca aataaagcaa tagcatcaca aatttcacaa
     3061 ataaagcatt tttttcactg cattctagtt gtggtttgtc caaactcatc aatgtatctt
     3121 atttaatgga ccgcccgcag ggggggtggc atttcagtgt cgggtgacga gcgcgatccg
     3181 gccgggatcc taggacccca aaagtttgtc tgcgtattcc agggcggggc tcagttgaat
     3241 ctcccgcagc acctctacca gcaggtccgc ggtgggctgg agaaactcgg ccgtcccggg
     3301 gcaggcggtc gtcgggggtg gaggcgcggc gcccaccccg tgtgccgcgc ctggcgtctc
     3361 ctctgggggc gacccgtaaa tggttgcagt gatgtaaatg gtgtccgcgg tccagaccac
     3421 ggtcaaaatg ccggccgtgg cgctccgggc gctttcgccg cgcgaggagc tgacccagga
     3481 gtcgaacgga tacgcgtaca tatgggcgtc ccacccgcgt tcgagcttct ggttgctgtc
     3541 ccggcctata aagcggtagg cacaaaattc ggcgcgacag tcgataatca ccaacagccc
     3601 aatgggggtg tgttggataa caacgcctcc gcgcggcagg cggtcctggc gctcccggcc
     3661 ccgtaccatg atcgcgcggg tgccgtactc aaaaacatgc accacctgcg cggcgtcggg
     3721 cagtgcgctg gtcagcgagg ccctggcgtg gcataggcta tacgcgatgg tcgtctgtgg
     3781 attggacat
//

Plasmid： V23 (Cas12f1 UL3-UL4 gene drive)    5092 bp DNA
FEATURES             Location/Qualifiers
     source          1..5092
                     /mol_type="other DNA"
                     /organism="synthetic DNA construct"
     CDS             1..708
                     /label=UL3
     misc_feature    748..996
                     /label=U6
     misc_feature    997..1125
                     /label=sgRNA
     terminator      1120..1125
                     /label=polIII terminator
     promoter        1132..1680
                     /label=EF1
     misc_feature    1693..1740
                     /label=NLS
     CDS             1741..3327
                     /label=Cas12f1
     misc_feature    3328..3381
                     /label=NLS
     misc_feature    3391..3444
                     /label=T2A
     CDS             3445..4203
                     /label=copGFP
     polyA_signal    4234..4441
                     /label=bGH polyA
                     /note="Unknown feature type:regulatory  bovine growth
                     hormone gene polyadenylation signal; bGH polyA"
     CDS             complement(4493..5092)
                     /label=UL4
ORIGIN
        1 atggttaaac ctctggtctc atacgggtcg gtgatgtcgg gcgtcggggg agagggagtt
       61 ccctctgcgc ttgcgattct agcctcgtgg ggctggacgt tcgacacgcc aaaccacgag
      121 tcagggatat cgccagatac gactcccgca gattccattc ggggggccgc tgtggcctca
      181 cctgaccaac ctttacacgg gggcccggaa cgggaggcca cagcgccgtc tttctcccca
      241 acgcgcgcgg atgacggccc gccctgtacc gacgggccct acgtgacgtt tgataccctg
      301 tttatggtgt cgtcgatcga cgaattaggg cgtcgccagc tcacggacac catccgcaag
      361 gacctgcggt tgtcgctggc caagtttagc attgcgtgca ccaagacctc ctcgttttcg
      421 ggaaacgccc cgcgccacca cagacgcggg gcgttccagc gcggcacgcg ggcgccgcgc
      481 agcaacaaaa gccttcagat gtttgtgttg tgcaaacgca cccacgccgc tcgagtgcga
      541 gagcagcttc gggtcgttat tcagtcccgc aagccgcgca agtattacac gcgatcttcg
      601 gacgggcggc tctgccccgc cgtccccgtg ttcgtccacg agttcgtctc gtccgagcca
      661 atgcgcctcc accgagataa cgtcatgctg gcctcggggg ccgagtaacc gcccccccgc
      721 gccaccctca ctgcccgtcg cggtaccgag ggcctatttc ccatgattcc ttcatatttg
      781 catatacgat acaaggctgt tagagagata attggaatta atttgactgt aaacacaaag
      841 atattagtac aaaatacgtg acgtagaaag taataatttc ttgggtagtt tgcagtttta
      901 aaattatgtt ttaaaatgga ctatcatatg cttaccgtaa cttgaaagta tttcgatttc
      961 ttggctttat atatcttgtg gaaaggacga aacaccaccg cttcaccgag tgaaggtggg
     1021 ctgcttgcat cagcctaatg tcgagaagtg ctttcttcgg aaagtaaccc tcgaaacaaa
     1081 gaaaggaatg caacatgtta ataaataacg catattttat ttttttctag ataaaaggat
     1141 ctgcgatcgc tccggtgccc gtcagtgggc agagcgcaca tcgcccacag tccccgagaa
     1201 gttgggggga ggggtcggca attgaacggg tgcctagaga aggtggcgcg gggtaaactg
     1261 ggaaagtgat gtcgtgtact ggctccgcct ttttcccgag ggtgggggag aaccgtatat
     1321 aagtgcagta gtcgccgtga acgttctttt tcgcaacggg tttgccgcca gaacacagct
     1381 gaagcttcga ggggctcgca tctctccttc acgcgcccgc cgccctacct gaggccgcca
     1441 tccacgccgg ttgagtcgcg ttctgccgcc tcccgcctgt ggtgcctcct gaactgcgtc
     1501 cgccgtctag gtaagtttaa agctcaggtc gagaccgggc ctttgtccgg cgctcccttg
     1561 gagcctacct agactcagcc ggctctccac gctttgcctg accctgcttg ctcaactcta
     1621 cgtctttgtt tcgttttctg ttctgcgccg ttacagatcc aagctgtgac cggcgcctac
     1681 accggtgcca ccatgccaaa gaagaagcgg aaggtcggta tccacggagt cccagcagcc
     1741 atggccaaga acacaattac aaagacactg aagctgagga tcgtgagacc atacaacagc
     1801 gctgaggtcg agaagattgt ggctgatgaa aagaacaaca gggaaaagat cgccctcgag
     1861 aagaacaagg ataaggtgaa ggaggcctgc tctaagcacc tgaaagtggc cgcctactgc
     1921 accacacagg tggagaggaa cgcctgtctg ttttgtaaag ctcggaagct ggatgataag
     1981 ttttaccaga agctgcgggg ccagttcccc gatgccgtct tttggcagga gattagcgag
     2041 atcttcagac agctgcagaa gcaggccgcc gagatctaca accagagcct gatcgagctc
     2101 tactacgaga tcttcatcaa gggcaagggc attgccaacg cctcctccgt ggagcactac
     2161 ctgagcgacg tgtgctacac aagagccgcc gagctcttta agaacgccgc tatcgcttcc
     2221 gggctgagga gcaagattaa gagtaacttc cggctcaagg agctgaagaa catgaagagc
     2281 ggcctgccca ctacaaagag cgacaacttc ccaattccac tggtgaagca gaaggggggc
     2341 cagtacacag ggttcgagat ttccaaccac aacagcgact ttattattaa gatccccttt
     2401 ggcaggtggc aggtcaagaa ggagattgac aagtacaggc cctgggagaa gtttgatttc
     2461 gagcaggtgc agaagagccc caagcctatt tccctgctgc tgtccacaca gcggcggaag
     2521 aggaacaagg ggtggtctaa ggatgagggg accgaggccg agattaagaa agtgatgaac
     2581 ggcgactacc agacaagcta catcgaggtc aagcggggca gtaagattgg cgagaagagc
     2641 gcctggatgc tgaacctgag cattgacgtg ccaaagattg ataagggcgt ggatcccagc
     2701 atcatcggag ggatcgatgt gggggtcaag agccccctcg tgtgcgccat caacaacgcc
     2761 ttcagcaggt acagcatctc cgataacgac ctgttccact ttaacaagaa gatgttcgcc
     2821 cggcggagga ttttgctcaa gaagaaccgg cacaagcggg ccggacacgg ggccaagaac
     2881 aagctcaagc ccatcactat cctgaccgag aagagcgaga ggttcaggaa gaagctcatc
     2941 gagagatggg cctgcgagat cgccgatttc tttattaaga acaaggtcgg aacagtgcag
     3001 atggagaacc tcgagagcat gaagaggaag gaggattcct acttcaacat tcggctgagg
     3061 gggttctggc cctacgctga gatgcagaac aagattgagt ttaagctgaa gcagtacggg
     3121 attgagatcc ggaaggtggc ccccaacaac accagcaaga cctgcagcaa gtgcgggcac
     3181 ctcaacaact acttcaactt cgagtaccgg aagaagaaca agttcccaca cttcaagtgc
     3241 gagaagtgca actttaagga gaacgccgat tacaacgccg ccctgaacat cagcaaccct
     3301 aagctgaaga gcactaagga ggagcccaaa aggccggcgg ccacgaaaaa ggccggccag
     3361 gcaaaaaaga aaaaggaatt cggcagtgga gagggcagag gaagtctgct aacatgcggt
     3421 gacgtcgagg agaatcctgg cccaatggag agcgacgaga gcggcctgcc cgccatggag
     3481 atcgagtgcc gcatcaccgg caccctgaac ggcgtggagt tcgagctggt gggcggcgga
     3541 gagggcaccc ccaagcaggg ccgcatgacc aacaagatga agagcaccaa aggcgccctg
     3601 accttcagcc cctacctgct gagccacgtg atgggctacg gcttctacca cttcggcacc
     3661 taccccagcg gctacgagaa ccccttcctg cacgccatca acaacggcgg ctacaccaac
     3721 acccgcatcg agaagtacga ggacggcggc gtgctgcacg tgagcttcag ctaccgctac
     3781 gaggccggcc gcgtgatcgg cgacttcaag gtggtgggca ccggcttccc cgaggacagc
     3841 gtgatcttca ccgacaagat catccgcagc aacgccaccg tggagcacct gcaccccatg
     3901 ggcgataacg tgctggtggg cagcttcgcc cgcaccttca gcctgcgcga cggcggctac
     3961 tacagcttcg tggtggacag ccacatgcac ttcaagagcg ccatccaccc cagcatcctg
     4021 cagaacgggg gccccatgtt cgccttccgc cgcgtggagg agctgcacag caacaccgag
     4081 ctgggcatcg tggagtacca gcacgccttc aagaccccca tcgccttcgc cagatcccgc
     4141 gctcagtcgt ccaattctgc cgtggacggc accgccggac ccggctccac cggatctcgc
     4201 taagaattcc tagagctcgc tgatcagcct cgactgtgcc ttctagttgc cagccatctg
     4261 ttgtttgccc ctcccccgtg ccttccttga ccctggaagg tgccactccc actgtccttt
     4321 cctaataaaa tgaggaaatt gcatcgcatt gtctgagtag gtgtcattct attctggggg
     4381 gtggggtggg gcaggacagc aagggggagg attgggaaga gaatagcagg catgctgggg
     4441 agcggccgct ggcatttcag tgtcgggtga cgagcgcgat ccggccggga tcctaggacc
     4501 ccaaaagttt gtctgcgtat tccagggcgg ggctcagttg aatctcccgc agcacctcta
     4561 ccagcaggtc cgcggtgggc tggagaaact cggccgtccc ggggcaggcg gtcgtcgggg
     4621 gtggaggcgc ggcgcccacc ccgtgtgccg cgcctggcgt ctcctctggg ggcgacccgt
     4681 aaatggttgc agtgatgtaa atggtgtccg cggtccagac cacggtcaaa atgccggccg
     4741 tggcgctccg ggcgctttcg ccgcgcgagg agctgaccca ggagtcgaac ggatacgcgt
     4801 acatatgggc gtcccacccg cgttcgagct tctggttgct gtcccggcct ataaagcggt
     4861 aggcacaaaa ttcggcgcga cagtcgataa tcaccaacag cccaatgggg gtgtgttgga
     4921 taacaacgcc tccgcgcggc aggcggtcct ggcgctcccg gccccgtacc atgatcgcgc
     4981 gggtgccgta ctcaaaaaca tgcaccacct gcgcggcgtc gggcagtgcg ctggtcagcg
     5041 aggccctggc gtggcatagg ctatacgcga tggtcgtctg tggattggac at
//

Plasmid： V24 (Cas12f1 UL3-UL4 gene drive)      5092 bp DNA
FEATURES             Location/Qualifiers
     source          1..5092
                     /mol_type="other DNA"
                     /organism="synthetic DNA construct"
     CDS             1..708
                     /label=UL3
     misc_feature    748..996
                     /label=U6
     misc_feature    875..1119
                     /label=v24 pcr
     misc_feature    997..1125
                     /label=sgRNA
     misc_feature    1095..1114
                     /label=UL3 sgRNA (2)
     terminator      1120..1125
                     /label=polIII terminator
     promoter        1132..1680
                     /label=EF1
     misc_feature    1693..1740
                     /label=NLS
     CDS             1741..3327
                     /label=Cas12f1
     misc_feature    3328..3381
                     /label=NLS
     CDS             3328..3375
                     /codon_start=1
                     /product="bipartite nuclear localization signal from
                     nucleoplasmin"
                     /label=nucleoplasmin NLS
                     /translation="KRPAATKKAGQAKKKK"
     misc_feature    3391..3444
                     /label=T2A
     CDS             3445..4203
                     /label=copGFP
     polyA_signal    4234..4441
                     /label=bGH polyA
                     /note="Unknown feature type:regulatory  bovine growth
                     hormone gene polyadenylation signal; bGH polyA"
     CDS             complement(4493..5092)
                     /label=UL4
ORIGIN
        1 atggttaaac ctctggtctc atacgggtcg gtgatgtcgg gcgtcggggg agagggagtt
       61 ccctctgcgc ttgcgattct agcctcgtgg ggctggacgt tcgacacgcc aaaccacgag
      121 tcagggatat cgccagatac gactcccgca gattccattc ggggggccgc tgtggcctca
      181 cctgaccaac ctttacacgg gggcccggaa cgggaggcca cagcgccgtc tttctcccca
      241 acgcgcgcgg atgacggccc gccctgtacc gacgggccct acgtgacgtt tgataccctg
      301 tttatggtgt cgtcgatcga cgaattaggg cgtcgccagc tcacggacac catccgcaag
      361 gacctgcggt tgtcgctggc caagtttagc attgcgtgca ccaagacctc ctcgttttcg
      421 ggaaacgccc cgcgccacca cagacgcggg gcgttccagc gcggcacgcg ggcgccgcgc
      481 agcaacaaaa gccttcagat gtttgtgttg tgcaaacgca cccacgccgc tcgagtgcga
      541 gagcagcttc gggtcgttat tcagtcccgc aagccgcgca agtattacac gcgatcttcg
      601 gacgggcggc tctgccccgc cgtccccgtg ttcgtccacg agttcgtctc gtccgagcca
      661 atgcgcctcc accgagataa cgtcatgctg gcctcggggg ccgagtaacc gcccccccgc
      721 gccaccctca ctgcccgtcg cggtaccgag ggcctatttc ccatgattcc ttcatatttg
      781 catatacgat acaaggctgt tagagagata attggaatta atttgactgt aaacacaaag
      841 atattagtac aaaatacgtg acgtagaaag taataatttc ttgggtagtt tgcagtttta
      901 aaattatgtt ttaaaatgga ctatcatatg cttaccgtaa cttgaaagta tttcgatttc
      961 ttggctttat atatcttgtg gaaaggacga aacaccaccg cttcaccgag tgaaggtggg
     1021 ctgcttgcat cagcctaatg tcgagaagtg ctttcttcgg aaagtaaccc tcgaaacaaa
     1081 gaaaggaatg caacttaaca tcaaacacgc gcgattttat ttttttctag ataaaaggat
     1141 ctgcgatcgc tccggtgccc gtcagtgggc agagcgcaca tcgcccacag tccccgagaa
     1201 gttgggggga ggggtcggca attgaacggg tgcctagaga aggtggcgcg gggtaaactg
     1261 ggaaagtgat gtcgtgtact ggctccgcct ttttcccgag ggtgggggag aaccgtatat
     1321 aagtgcagta gtcgccgtga acgttctttt tcgcaacggg tttgccgcca gaacacagct
     1381 gaagcttcga ggggctcgca tctctccttc acgcgcccgc cgccctacct gaggccgcca
     1441 tccacgccgg ttgagtcgcg ttctgccgcc tcccgcctgt ggtgcctcct gaactgcgtc
     1501 cgccgtctag gtaagtttaa agctcaggtc gagaccgggc ctttgtccgg cgctcccttg
     1561 gagcctacct agactcagcc ggctctccac gctttgcctg accctgcttg ctcaactcta
     1621 cgtctttgtt tcgttttctg ttctgcgccg ttacagatcc aagctgtgac cggcgcctac
     1681 accggtgcca ccatgccaaa gaagaagcgg aaggtcggta tccacggagt cccagcagcc
     1741 atggccaaga acacaattac aaagacactg aagctgagga tcgtgagacc atacaacagc
     1801 gctgaggtcg agaagattgt ggctgatgaa aagaacaaca gggaaaagat cgccctcgag
     1861 aagaacaagg ataaggtgaa ggaggcctgc tctaagcacc tgaaagtggc cgcctactgc
     1921 accacacagg tggagaggaa cgcctgtctg ttttgtaaag ctcggaagct ggatgataag
     1981 ttttaccaga agctgcgggg ccagttcccc gatgccgtct tttggcagga gattagcgag
     2041 atcttcagac agctgcagaa gcaggccgcc gagatctaca accagagcct gatcgagctc
     2101 tactacgaga tcttcatcaa gggcaagggc attgccaacg cctcctccgt ggagcactac
     2161 ctgagcgacg tgtgctacac aagagccgcc gagctcttta agaacgccgc tatcgcttcc
     2221 gggctgagga gcaagattaa gagtaacttc cggctcaagg agctgaagaa catgaagagc
     2281 ggcctgccca ctacaaagag cgacaacttc ccaattccac tggtgaagca gaaggggggc
     2341 cagtacacag ggttcgagat ttccaaccac aacagcgact ttattattaa gatccccttt
     2401 ggcaggtggc aggtcaagaa ggagattgac aagtacaggc cctgggagaa gtttgatttc
     2461 gagcaggtgc agaagagccc caagcctatt tccctgctgc tgtccacaca gcggcggaag
     2521 aggaacaagg ggtggtctaa ggatgagggg accgaggccg agattaagaa agtgatgaac
     2581 ggcgactacc agacaagcta catcgaggtc aagcggggca gtaagattgg cgagaagagc
     2641 gcctggatgc tgaacctgag cattgacgtg ccaaagattg ataagggcgt ggatcccagc
     2701 atcatcggag ggatcgatgt gggggtcaag agccccctcg tgtgcgccat caacaacgcc
     2761 ttcagcaggt acagcatctc cgataacgac ctgttccact ttaacaagaa gatgttcgcc
     2821 cggcggagga ttttgctcaa gaagaaccgg cacaagcggg ccggacacgg ggccaagaac
     2881 aagctcaagc ccatcactat cctgaccgag aagagcgaga ggttcaggaa gaagctcatc
     2941 gagagatggg cctgcgagat cgccgatttc tttattaaga acaaggtcgg aacagtgcag
     3001 atggagaacc tcgagagcat gaagaggaag gaggattcct acttcaacat tcggctgagg
     3061 gggttctggc cctacgctga gatgcagaac aagattgagt ttaagctgaa gcagtacggg
     3121 attgagatcc ggaaggtggc ccccaacaac accagcaaga cctgcagcaa gtgcgggcac
     3181 ctcaacaact acttcaactt cgagtaccgg aagaagaaca agttcccaca cttcaagtgc
     3241 gagaagtgca actttaagga gaacgccgat tacaacgccg ccctgaacat cagcaaccct
     3301 aagctgaaga gcactaagga ggagcccaaa aggccggcgg ccacgaaaaa ggccggccag
     3361 gcaaaaaaga aaaaggaatt cggcagtgga gagggcagag gaagtctgct aacatgcggt
     3421 gacgtcgagg agaatcctgg cccaatggag agcgacgaga gcggcctgcc cgccatggag
     3481 atcgagtgcc gcatcaccgg caccctgaac ggcgtggagt tcgagctggt gggcggcgga
     3541 gagggcaccc ccaagcaggg ccgcatgacc aacaagatga agagcaccaa aggcgccctg
     3601 accttcagcc cctacctgct gagccacgtg atgggctacg gcttctacca cttcggcacc
     3661 taccccagcg gctacgagaa ccccttcctg cacgccatca acaacggcgg ctacaccaac
     3721 acccgcatcg agaagtacga ggacggcggc gtgctgcacg tgagcttcag ctaccgctac
     3781 gaggccggcc gcgtgatcgg cgacttcaag gtggtgggca ccggcttccc cgaggacagc
     3841 gtgatcttca ccgacaagat catccgcagc aacgccaccg tggagcacct gcaccccatg
     3901 ggcgataacg tgctggtggg cagcttcgcc cgcaccttca gcctgcgcga cggcggctac
     3961 tacagcttcg tggtggacag ccacatgcac ttcaagagcg ccatccaccc cagcatcctg
     4021 cagaacgggg gccccatgtt cgccttccgc cgcgtggagg agctgcacag caacaccgag
     4081 ctgggcatcg tggagtacca gcacgccttc aagaccccca tcgccttcgc cagatcccgc
     4141 gctcagtcgt ccaattctgc cgtggacggc accgccggac ccggctccac cggatctcgc
     4201 taagaattcc tagagctcgc tgatcagcct cgactgtgcc ttctagttgc cagccatctg
     4261 ttgtttgccc ctcccccgtg ccttccttga ccctggaagg tgccactccc actgtccttt
     4321 cctaataaaa tgaggaaatt gcatcgcatt gtctgagtag gtgtcattct attctggggg
     4381 gtggggtggg gcaggacagc aagggggagg attgggaaga gaatagcagg catgctgggg
     4441 agcggccgct ggcatttcag tgtcgggtga cgagcgcgat ccggccggga tcctaggacc
     4501 ccaaaagttt gtctgcgtat tccagggcgg ggctcagttg aatctcccgc agcacctcta
     4561 ccagcaggtc cgcggtgggc tggagaaact cggccgtccc ggggcaggcg gtcgtcgggg
     4621 gtggaggcgc ggcgcccacc ccgtgtgccg cgcctggcgt ctcctctggg ggcgacccgt
     4681 aaatggttgc agtgatgtaa atggtgtccg cggtccagac cacggtcaaa atgccggccg
     4741 tggcgctccg ggcgctttcg ccgcgcgagg agctgaccca ggagtcgaac ggatacgcgt
     4801 acatatgggc gtcccacccg cgttcgagct tctggttgct gtcccggcct ataaagcggt
     4861 aggcacaaaa ttcggcgcga cagtcgataa tcaccaacag cccaatgggg gtgtgttgga
     4921 taacaacgcc tccgcgcggc aggcggtcct ggcgctcccg gccccgtacc atgatcgcgc
     4981 gggtgccgta ctcaaaaaca tgcaccacct gcgcggcgtc gggcagtgcg ctggtcagcg
     5041 aggccctggc gtggcatagg ctatacgcga tggtcgtctg tggattggac at
//

Plasmid： V25 (Cas9 UL3-UL4 gene drive)     8384 bp DNA
FEATURES             Location/Qualifiers
     source          1..8384
                     /mol_type="genomic DNA"
                     /organism="unspecified"
     misc_feature    1..1016
                     /label=UL3
     promoter        1025..1264
                     /label=U6 promoter
                     /note="RNA polymerase III promoter for human U6 snRNA"
     misc_feature    1274..1293
                     /label=sgRNA
     misc_RNA        1294..1369
                     /label=gRNA scaffold
                     /note="guide RNA scaffold for the Streptococcus pyogenes
                     CRISPR/Cas9 system"
     enhancer        1419..1704
                     /label=CMV enhancer
                     /note="human cytomegalovirus immediate early enhancer;
                     contains an 18-bp deletion relative to the standard CMV
                     enhancer"
     promoter        1706..1984
                     /label=chicken beta-actin promoter
     intron          1985..2212
                     /label=hybrid intron
                     /note="hybrid between chicken beta-actin (CBA) and minute
                     virus of mice (MMV) introns (Gray et al., 2011)"
     CDS             2236..2301
                     /codon_start=1
                     /product="three tandem FLAG(R) epitope tags, followed by an
                     enterokinase cleavage site"
                     /label=3xFLAG
                     /translation="DYKDHDGDYKDHDIDYKDDDDK"
     CDS             2308..2328
                     /codon_start=1
                     /product="nuclear localization signal of SV40 (simian virus
                     40) large T antigen"
                     /label=SV40 NLS
                     /translation="PKKKRKV"
     CDS             2353..6453
                     /codon_start=1
                     /product="Cas9 (Csn1) endonuclease from the Streptococcus
                     pyogenes Type II CRISPR/Cas system"
                     /label=Cas9
                     /note="generates RNA-guided double strand breaks in DNA"
                     /translation="DKKYSIGLDIGTNSVGWAVITDEYKVPSKKFKVLGNTDRHSIKKN
                     LIGALLFDSGETAEATRLKRTARRRYTRRKNRICYLQEIFSNEMAKVDDSFFHRLEESF
                     LVEEDKKHERHPIFGNIVDEVAYHEKYPTIYHLRKKLVDSTDKADLRLIYLALAHMIKF
                     RGHFLIEGDLNPDNSDVDKLFIQLVQTYNQLFEENPINASGVDAKAILSARLSKSRRLE
                     NLIAQLPGEKKNGLFGNLIALSLGLTPNFKSNFDLAEDAKLQLSKDTYDDDLDNLLAQI
                     GDQYADLFLAAKNLSDAILLSDILRVNTEITKAPLSASMIKRYDEHHQDLTLLKALVRQ
                     QLPEKYKEIFFDQSKNGYAGYIDGGASQEEFYKFIKPILEKMDGTEELLVKLNREDLLR
                     KQRTFDNGSIPHQIHLGELHAILRRQEDFYPFLKDNREKIEKILTFRIPYYVGPLARGN
                     SRFAWMTRKSEETITPWNFEEVVDKGASAQSFIERMTNFDKNLPNEKVLPKHSLLYEYF
                     TVYNELTKVKYVTEGMRKPAFLSGEQKKAIVDLLFKTNRKVTVKQLKEDYFKKIECFDS
                     VEISGVEDRFNASLGTYHDLLKIIKDKDFLDNEENEDILEDIVLTLTLFEDREMIEERL
                     KTYAHLFDDKVMKQLKRRRYTGWGRLSRKLINGIRDKQSGKTILDFLKSDGFANRNFMQ
                     LIHDDSLTFKEDIQKAQVSGQGDSLHEHIANLAGSPAIKKGILQTVKVVDELVKVMGRH
                     KPENIVIEMARENQTTQKGQKNSRERMKRIEEGIKELGSQILKEHPVENTQLQNEKLYL
                     YYLQNGRDMYVDQELDINRLSDYDVDHIVPQSFLKDDSIDNKVLTRSDKNRGKSDNVPS
                     EEVVKKMKNYWRQLLNAKLITQRKFDNLTKAERGGLSELDKAGFIKRQLVETRQITKHV
                     AQILDSRMNTKYDENDKLIREVKVITLKSKLVSDFRKDFQFYKVREINNYHHAHDAYLN
                     AVVGTALIKKYPKLESEFVYGDYKVYDVRKMIAKSEQEIGKATAKYFFYSNIMNFFKTE
                     ITLANGEIRKRPLIETNGETGEIVWDKGRDFATVRKVLSMPQVNIVKKTEVQTGGFSKE
                     SILPKRNSDKLIARKKDWDPKKYGGFDSPTVAYSVLVVAKVEKGKSKKLKSVKELLGIT
                     IMERSSFEKNPIDFLEAKGYKEVKKDLIIKLPKYSLFELENGRKRMLASAGELQKGNEL
                     ALPSKYVNFLYLASHYEKLKGSPEDNEQKQLFVEQHKHYLDEIIEQISEFSKRVILADA
                     NLDKVLSAYNKHRDKPIREQAENIIHLFTLTNLGAPAAFKYFDTTIDRKRYTSTKEVLD
                     ATLIHQSITGLYETRIDLSQLGGD"
     CDS             6454..6501
                     /codon_start=1
                     /product="bipartite nuclear localization signal from
                     nucleoplasmin"
                     /label=nucleoplasmin NLS
                     /translation="KRPAATKKAGQAKKKK"
     CDS             6502..6555
                     /codon_start=1
                     /product="2A peptide from Thosea asigna virus capsid
                     protein"
                     /label=T2A
                     /note="Eukaryotic ribosomes fail to insert a peptide bond
                     between the Gly and Pro residues, yielding separate
                     polypeptides."
                     /translation="EGRGSLLTCGDVEENPGP"
     CDS             6580..7242
                     /codon_start=1
                     /product="green fluorescent protein 2 from Pontellina
                     plumata, also known as ppluGFP2 (Shagin et al., 2004)"
                     /label=CopGFP
                     /translation="PAMEIECRITGTLNGVEFELVGGGEGTPKQGRMTNKMKSTKGALT
                     FSPYLLSHVMGYGFYHFGTYPSGYENPFLHAINNGGYTNTRIEKYEDGGVLHVSFSYRY
                     EAGRVIGDFKVVGTGFPEDSVIFTDKIIRSNATVEHLHPMGDNVLVGSFARTFSLRDGG
                     YYSFVVDSHMHFKSAIHPSILQNGGPMFAFRRVEELHSNTELGIVEYQHAFKTPIAFA"
     polyA_signal    7345..7552
                     /label=bGH poly(A) signal
                     /note="bovine growth hormone polyadenylation signal"
     misc_feature    7553..8384
                     /label=UL4
ORIGIN
        1 tagaatcggt tgggaccgct tcgtgggcgg agttatccgc cggttggccg cgcgccgccc
       61 cggcctggtg tttatgctct ggggcgcaca tgcccagaat gccatcaggc cggaccctcg
      121 ggtccattgc gtcctcaagt tttcgcaccc gtcgcccctc tccaaggttc cgttcggaac
      181 atgccagcat ttcctcgtgg cgaatcgata tctcgagacc cggtcgattt cacccatcga
      241 ctggtcggtt tgaaaggcat cgacgtccgg ggttttcgtc tgtgggggct tttgggtatt
      301 tccgatgaat aaagacggtt aatggttaaa cctctggtct catacgggtc ggtgatgtcg
      361 ggcgtcgggg gagagggagt tccctctgcg cttgcgattc tagcctcgtg gggctggacg
      421 ttcgacacgc caaaccacga gtcagggata tcgccagata cgactcccgc agattccatt
      481 cggggggccg ctgtggcctc acctgaccaa cctttacacg ggggcccgga acgggaggcc
      541 acagcgccgt ctttctcccc aacgcgcgcg gatgacggcc cgccctgtac cgacgggccc
      601 tacgtgacgt ttgataccct gtttatggtg tcgtcgatcg acgaattagg gcgtcgccag
      661 ctcacggaca ccatccgcaa ggacctgcgg ttgtcgctgg ccaagtttag cattgcgtgc
      721 accaagacct cctcgttttc gggaaacgcc ccgcgccacc acagacgcgg ggcgttccag
      781 cgcggcacgc gggcgccgcg cagcaacaaa agccttcaga tgtttgtgtt gtgcaaacgc
      841 acccacgccg ctcgagtgcg agagcagctt cgggtcgtta ttcagtcccg caagccgcgc
      901 aagtattaca cgcgatcttc ggacgggcgg ctctgccccg ccgtccccgt gttcgtccac
      961 gagttcgtct cgtccgagcc aatgcgcctc caccgagata acgtcatgct ggcctccaca
     1021 tgtgagggcc tatttcccat gattccttca tatttgcata tacgatacaa ggctgttaga
     1081 gagataattg gaattaattt gactgtaaac acaaagatat tagtacaaaa tacgtgacgt
     1141 agaaagtaat aatttcttgg gtagtttgca gttttaaaat tatgttttaa aatggactat
     1201 catatgctta ccgtaacttg aaagtatttc gatttcttgg ctttatatat cttgtggaaa
     1261 ggacgaaaca ccgaaataac gcataaattt ggcgttttag agctagaaat agcaagttaa
     1321 aataaggcta gtccgttatc aacttgaaaa agtggcaccg agtcggtgct tttttgtcta
     1381 gaatggttgc agtgatgtaa atggtgtccg cgggtacccg ttacataact tacggtaaat
     1441 ggcccgcctg gctgaccgcc caacgacccc cgcccattga cgtcaatagt aacgccaata
     1501 gggactttcc attgacgtca atgggtggag tatttacggt aaactgccca cttggcagta
     1561 catcaagtgt atcatatgcc aagtacgccc cctattgacg tcaatgacgg taaatggccc
     1621 gcctggcatt gtgcccagta catgacctta tgggactttc ctacttggca gtacatctac
     1681 gtattagtca tcgctattac catggtcgag gtgagcccca cgttctgctt cactctcccc
     1741 atctcccccc cctccccacc cccaattttg tatttattta ttttttaatt attttgtgca
     1801 gcgatggggg cggggggggg gggggggcgc gcgccaggcg gggcggggcg ggggcgaggg
     1861 gcggggcggg gcgaggcgga gaggtgcggc ggcagccaat cagagcggcg cgctccgaaa
     1921 gtttcctttt atggcgaggc ggcggcggcg gcggccctat aaaaagcgaa gcgcgcggcg
     1981 ggcgggagtc gctgcgcgct gccttcgccc cgtgccccgc tccgccgccg cctcgcgccg
     2041 cccgccccgg ctctgactga ccgcgttact cccacaggtg agcgggcggg acggcccttc
     2101 tcctccgggc tgtaattagc tgagcaagag gtaagggttt aagggatggt tggttggtgg
     2161 ggtattaatg tttaattacc tggagcacct gcctgaaatc actttttttc aggttggacc
     2221 ggtgccacca tgatggacta taaggaccac gacggagact acaaggatca tgatattgat
     2281 tacaaagacg atgacgataa gatggcccca aagaagaagc ggaaggtcgg tatccacgga
     2341 gtcccagcag ccgacaagaa gtacagcatc ggcctggaca tcggcaccaa ctctgtgggc
     2401 tgggccgtga tcaccgacga gtacaaggtg cccagcaaga aattcaaggt gctgggcaac
     2461 accgaccggc acagcatcaa gaagaacctg atcggagccc tgctgttcga cagcggcgaa
     2521 acagccgagg ccacccggct gaagagaacc gccagaagaa gatacaccag acggaagaac
     2581 cggatctgct atctgcaaga gatcttcagc aacgagatgg ccaaggtgga cgacagcttc
     2641 ttccacagac tggaagagtc cttcctggtg gaagaggata agaagcacga gcggcacccc
     2701 atcttcggca acatcgtgga cgaggtggcc taccacgaga agtaccccac catctaccac
     2761 ctgagaaaga aactggtgga cagcaccgac aaggccgacc tgcggctgat ctatctggcc
     2821 ctggcccaca tgatcaagtt ccggggccac ttcctgatcg agggcgacct gaaccccgac
     2881 aacagcgacg tggacaagct gttcatccag ctggtgcaga cctacaacca gctgttcgag
     2941 gaaaacccca tcaacgccag cggcgtggac gccaaggcca tcctgtctgc cagactgagc
     3001 aagagcagac ggctggaaaa tctgatcgcc cagctgcccg gcgagaagaa gaatggcctg
     3061 ttcggaaacc tgattgccct gagcctgggc ctgaccccca acttcaagag caacttcgac
     3121 ctggccgagg atgccaaact gcagctgagc aaggacacct acgacgacga cctggacaac
     3181 ctgctggccc agatcggcga ccagtacgcc gacctgtttc tggccgccaa gaacctgtcc
     3241 gacgccatcc tgctgagcga catcctgaga gtgaacaccg agatcaccaa ggcccccctg
     3301 agcgcctcta tgatcaagag atacgacgag caccaccagg acctgaccct gctgaaagct
     3361 ctcgtgcggc agcagctgcc tgagaagtac aaagagattt tcttcgacca gagcaagaac
     3421 ggctacgccg gctacattga cggcggagcc agccaggaag agttctacaa gttcatcaag
     3481 cccatcctgg aaaagatgga cggcaccgag gaactgctcg tgaagctgaa cagagaggac
     3541 ctgctgcgga agcagcggac cttcgacaac ggcagcatcc cccaccagat ccacctggga
     3601 gagctgcacg ccattctgcg gcggcaggaa gatttttacc cattcctgaa ggacaaccgg
     3661 gaaaagatcg agaagatcct gaccttccgc atcccctact acgtgggccc tctggccagg
     3721 ggaaacagca gattcgcctg gatgaccaga aagagcgagg aaaccatcac cccctggaac
     3781 ttcgaggaag tggtggacaa gggcgcttcc gcccagagct tcatcgagcg gatgaccaac
     3841 ttcgataaga acctgcccaa cgagaaggtg ctgcccaagc acagcctgct gtacgagtac
     3901 ttcaccgtgt ataacgagct gaccaaagtg aaatacgtga ccgagggaat gagaaagccc
     3961 gccttcctga gcggcgagca gaaaaaggcc atcgtggacc tgctgttcaa gaccaaccgg
     4021 aaagtgaccg tgaagcagct gaaagaggac tacttcaaga aaatcgagtg cttcgactcc
     4081 gtggaaatct ccggcgtgga agatcggttc aacgcctccc tgggcacata ccacgatctg
     4141 ctgaaaatta tcaaggacaa ggacttcctg gacaatgagg aaaacgagga cattctggaa
     4201 gatatcgtgc tgaccctgac actgtttgag gacagagaga tgatcgagga acggctgaaa
     4261 acctatgccc acctgttcga cgacaaagtg atgaagcagc tgaagcggcg gagatacacc
     4321 ggctggggca ggctgagccg gaagctgatc aacggcatcc gggacaagca gtccggcaag
     4381 acaatcctgg atttcctgaa gtccgacggc ttcgccaaca gaaacttcat gcagctgatc
     4441 cacgacgaca gcctgacctt taaagaggac atccagaaag cccaggtgtc cggccagggc
     4501 gatagcctgc acgagcacat tgccaatctg gccggcagcc ccgccattaa gaagggcatc
     4561 ctgcagacag tgaaggtggt ggacgagctc gtgaaagtga tgggccggca caagcccgag
     4621 aacatcgtga tcgaaatggc cagagagaac cagaccaccc agaagggaca gaagaacagc
     4681 cgcgagagaa tgaagcggat cgaagagggc atcaaagagc tgggcagcca gatcctgaaa
     4741 gaacaccccg tggaaaacac ccagctgcag aacgagaagc tgtacctgta ctacctgcag
     4801 aatgggcggg atatgtacgt ggaccaggaa ctggacatca accggctgtc cgactacgat
     4861 gtggaccata tcgtgcctca gagctttctg aaggacgact ccatcgacaa caaggtgctg
     4921 accagaagcg acaagaaccg gggcaagagc gacaacgtgc cctccgaaga ggtcgtgaag
     4981 aagatgaaga actactggcg gcagctgctg aacgccaagc tgattaccca gagaaagttc
     5041 gacaatctga ccaaggccga gagaggcggc ctgagcgaac tggataaggc cggcttcatc
     5101 aagagacagc tggtggaaac ccggcagatc acaaagcacg tggcacagat cctggactcc
     5161 cggatgaaca ctaagtacga cgagaatgac aagctgatcc gggaagtgaa agtgatcacc
     5221 ctgaagtcca agctggtgtc cgatttccgg aaggatttcc agttttacaa agtgcgcgag
     5281 atcaacaact accaccacgc ccacgacgcc tacctgaacg ccgtcgtggg aaccgccctg
     5341 atcaaaaagt accctaagct ggaaagcgag ttcgtgtacg gcgactacaa ggtgtacgac
     5401 gtgcggaaga tgatcgccaa gagcgagcag gaaatcggca aggctaccgc caagtacttc
     5461 ttctacagca acatcatgaa ctttttcaag accgagatta ccctggccaa cggcgagatc
     5521 cggaagcggc ctctgatcga gacaaacggc gaaaccgggg agatcgtgtg ggataagggc
     5581 cgggattttg ccaccgtgcg gaaagtgctg agcatgcccc aagtgaatat cgtgaaaaag
     5641 accgaggtgc agacaggcgg cttcagcaaa gagtctatcc tgcccaagag gaacagcgat
     5701 aagctgatcg ccagaaagaa ggactgggac cctaagaagt acggcggctt cgacagcccc
     5761 accgtggcct attctgtgct ggtggtggcc aaagtggaaa agggcaagtc caagaaactg
     5821 aagagtgtga aagagctgct ggggatcacc atcatggaaa gaagcagctt cgagaagaat
     5881 cccatcgact ttctggaagc caagggctac aaagaagtga aaaaggacct gatcatcaag
     5941 ctgcctaagt actccctgtt cgagctggaa aacggccgga agagaatgct ggcctctgcc
     6001 ggcgaactgc agaagggaaa cgaactggcc ctgccctcca aatatgtgaa cttcctgtac
     6061 ctggccagcc actatgagaa gctgaagggc tcccccgagg ataatgagca gaaacagctg
     6121 tttgtggaac agcacaagca ctacctggac gagatcatcg agcagatcag cgagttctcc
     6181 aagagagtga tcctggccga cgctaatctg gacaaagtgc tgtccgccta caacaagcac
     6241 cgggataagc ccatcagaga gcaggccgag aatatcatcc acctgtttac cctgaccaat
     6301 ctgggagccc ctgccgcctt caagtacttt gacaccacca tcgaccggaa gaggtacacc
     6361 agcaccaaag aggtgctgga cgccaccctg atccaccaga gcatcaccgg cctgtacgag
     6421 acacggatcg acctgtctca gctgggaggc gacaaaaggc cggcggccac gaaaaaggcc
     6481 ggccaggcaa aaaagaaaaa ggagggcaga ggaagtctac taacatgcgg tgacgtggag
     6541 gagaatcccg gccctatgga gagcgacgag agcggcctgc ccgccatgga gatcgagtgc
     6601 cgcatcaccg gcaccctgaa cggcgtggag ttcgagctgg tgggcggcgg agagggcacc
     6661 cccaagcagg gccgcatgac caacaagatg aagagcacca aaggcgccct gaccttcagc
     6721 ccctacctgc tgagccacgt gatgggctac ggcttctacc acttcggcac ctaccccagc
     6781 ggctacgaga accccttcct gcacgccatc aacaacggcg gctacaccaa cacccgcatc
     6841 gagaagtacg aggacggcgg cgtgctgcac gtgagcttca gctaccgcta cgaggccggc
     6901 cgcgtgatcg gcgacttcaa ggtggtgggc accggcttcc ccgaggacag cgtgatcttc
     6961 accgacaaga tcatccgcag caacgccacc gtggagcacc tgcaccccat gggcgataac
     7021 gtgctggtgg gcagcttcgc ccgcaccttc agcctgcgcg acggcggcta ctacagcttc
     7081 gtggtggaca gccacatgca cttcaagagc gccatccacc ccagcatcct gcagaacggg
     7141 ggccccatgt tcgccttccg ccgcgtggag gagctgcaca gcaacaccga gctgggcatc
     7201 gtggagtacc agcacgcctt caagaccccc atcgccttcg ccagatcccg cgctcagtcg
     7261 tccaattctg ccgtggacgg caccgccgga cccggctcca ccggatctcg ctaagaattc
     7321 ctagagctcg ctgatcagcc tcgactgtgc cttctagttg ccagccatct gttgtttgcc
     7381 cctcccccgt gccttccttg accctggaag gtgccactcc cactgtcctt tcctaataaa
     7441 atgaggaaat tgcatcgcat tgtctgagta ggtgtcattc tattctgggg ggtggggtgg
     7501 ggcaggacag caagggggag gattgggaag agaatagcag gcatgctggg gaaccacggt
     7561 caaaatgccg gccgtggcgc tccgggcgct ttcgccgcgc gaggagctga cccaggagtc
     7621 gaacggatac gcgtacatat gggcgtccca cccgcgttcg agcttctggt tgctgtcccg
     7681 gcctataaag cggtaggcac aaaattcggc gcgacagtcg ataatcacca acagcccaat
     7741 gggggtgtgt tggataacaa cgcctccgcg cggcaggcgg tcctggcgct cccggccccg
     7801 taccatgatc gcgcgggtgc cgtactcaaa aacatgcacc acctgcgcgg cgtcgggcag
     7861 tgcgctggtc agcgaggccc tggcgtggca taggctatac gcgatggtcg tctgtggatt
     7921 ggacatctcg cggtgggtag tgagtccccc gggccgggtt cggtggaact gtaaggggac
     7981 ggcgggttaa tatacaatga ccacgttcgg atcgcgcaga gccgatagta tgtgcttact
     8041 aatgacgtca tcgcgctcgt ggcgctcccg gagcggattt aagttcatgc gaaggaattc
     8101 ggaggaggtg gtgcgggaca tggccacgta cgcgctgttg aggcgcaggt tgccgggcgt
     8161 aaagcagatg gcgaccttgt ccaggctaag gccctgggag cgcgtgatgg tcatggcaag
     8221 cttggagctg atgccgtagt cggcgtttat ggccatggcc agctccgtag agtcaatgga
     8281 ctcgacaaac tcgctgatgt tggtgttgac gacggacatg aagccgtgtt ggtcccgcaa
     8341 gaccacgtaa ggcagggggg cctcttccag taactcggcc acgt
//

Plasmid： V28 (Cas9 UL3-UL4 gene drive)    7644 bp DNA
FEATURES             Location/Qualifiers
     source          1..7644
                     /mol_type="other DNA"
                     /organism="synthetic DNA construct"
     CDS             1..708
                     /label=UL3
     misc_feature    748..996
                     /label=U6
                     /note="Unknown feature type:regulatory  human U6 promoter"
     misc_feature    997..1017
                     /label=gRNA
     misc_feature    1018..1093
                     /label=gRNA scaffold
                     /note="guide RNA scaffold for the Streptococcus pyogenes
                     CRISPR/Cas9 system; gRNA scaffold"
     terminator      1094..1099
                     /label=polIII terminator
                     /note="Unknown feature type:regulatory  RNA polymerase III
                     transcription terminator; polIII terminator"
     promoter        1113..1661
                     /label=EF1
     CDS             1674..5942
                     /codon_start=1
                     /label=NLS Cas9 NLS
                     /translation="MDYKDHDGDYKDHDIDYKDDDDKMAPKKKRKVGIHGVPAADKKYS
                     IGLDIGTNSVGWAVITDEYKVPSKKFKVLGNTDRHSIKKNLIGALLFDSGETAEATRLK
                     RTARRRYTRRKNRICYLQEIFSNEMAKVDDSFFHRLEESFLVEEDKKHERHPIFGNIVD
                     EVAYHEKYPTIYHLRKKLVDSTDKADLRLIYLALAHMIKFRGHFLIEGDLNPDNSDVDK
                     LFIQLVQTYNQLFEENPINASGVDAKAILSARLSKSRRLENLIAQLPGEKKNGLFGNLI
                     ALSLGLTPNFKSNFDLAEDAKLQLSKDTYDDDLDNLLAQIGDQYADLFLAAKNLSDAIL
                     LSDILRVNTEITKAPLSASMIKRYDEHHQDLTLLKALVRQQLPEKYKEIFFDQSKNGYA
                     GYIDGGASQEEFYKFIKPILEKMDGTEELLVKLNREDLLRKQRTFDNGSIPHQIHLGEL
                     HAILRRQEDFYPFLKDNREKIEKILTFRIPYYVGPLARGNSRFAWMTRKSEETITPWNF
                     EEVVDKGASAQSFIERMTNFDKNLPNEKVLPKHSLLYEYFTVYNELTKVKYVTEGMRKP
                     AFLSGEQKKAIVDLLFKTNRKVTVKQLKEDYFKKIECFDSVEISGVEDRFNASLGTYHD
                     LLKIIKDKDFLDNEENEDILEDIVLTLTLFEDREMIEERLKTYAHLFDDKVMKQLKRRR
                     YTGWGRLSRKLINGIRDKQSGKTILDFLKSDGFANRNFMQLIHDDSLTFKEDIQKAQVS
                     GQGDSLHEHIANLAGSPAIKKGILQTVKVVDELVKVMGRHKPENIVIEMARENQTTQKG
                     QKNSRERMKRIEEGIKELGSQILKEHPVENTQLQNEKLYLYYLQNGRDMYVDQELDINR
                     LSDYDVDHIVPQSFLKDDSIDNKVLTRSDKNRGKSDNVPSEEVVKKMKNYWRQLLNAKL
                     ITQRKFDNLTKAERGGLSELDKAGFIKRQLVETRQITKHVAQILDSRMNTKYDENDKLI
                     REVKVITLKSKLVSDFRKDFQFYKVREINNYHHAHDAYLNAVVGTALIKKYPKLESEFV
                     YGDYKVYDVRKMIAKSEQEIGKATAKYFFYSNIMNFFKTEITLANGEIRKRPLIETNGE
                     TGEIVWDKGRDFATVRKVLSMPQVNIVKKTEVQTGGFSKESILPKRNSDKLIARKKDWD
                     PKKYGGFDSPTVAYSVLVVAKVEKGKSKKLKSVKELLGITIMERSSFEKNPIDFLEAKG
                     YKEVKKDLIIKLPKYSLFELENGRKRMLASAGELQKGNELALPSKYVNFLYLASHYEKL
                     KGSPEDNEQKQLFVEQHKHYLDEIIEQISEFSKRVILADANLDKVLSAYNKHRDKPIRE
                     QAENIIHLFTLTNLGAPAAFKYFDTTIDRKRYTSTKEVLDATLIHQSITGLYETRIDLS
                     QLGGDKRPAATKKAGQAKKKK"
     CDS             5997..6755
                     /codon_start=1
                     /label=copGFP
                     /translation="MESDESGLPAMEIECRITGTLNGVEFELVGGGEGTPKQGRMTNKM
                     KSTKGALTFSPYLLSHVMGYGFYHFGTYPSGYENPFLHAINNGGYTNTRIEKYEDGGVL
                     HVSFSYRYEAGRVIGDFKVVGTGFPEDSVIFTDKIIRSNATVEHLHPMGDNVLVGSFAR
                     TFSLRDGGYYSFVVDSHMHFKSAIHPSILQNGGPMFAFRRVEELHSNTELGIVEYQHAF
                     KTPIAFARSRAQSSNSAVDGTAGPGSTGSR"
     polyA_signal    6786..6993
                     /label=bGH polyA
                     /note="Unknown feature type:regulatory  bovine growth
                     hormone gene polyadenylation signal; bGH polyA"
     CDS             complement(7045..7644)
                     /label=UL4
ORIGIN
        1 atggttaaac ctctggtctc atacgggtcg gtgatgtcgg gcgtcggggg agagggagtt
       61 ccctctgcgc ttgcgattct agcctcgtgg ggctggacgt tcgacacgcc aaaccacgag
      121 tcagggatat cgccagatac gactcccgca gattccattc ggggggccgc tgtggcctca
      181 cctgaccaac ctttacacgg gggcccggaa cgggaggcca cagcgccgtc tttctcccca
      241 acgcgcgcgg atgacggccc gccctgtacc gacgggccct acgtgacgtt tgataccctg
      301 tttatggtgt cgtcgatcga cgaattaggg cgtcgccagc tcacggacac catccgcaag
      361 gacctgcggt tgtcgctggc caagtttagc attgcgtgca ccaagacctc ctcgttttcg
      421 ggaaacgccc cgcgccacca cagacgcggg gcgttccagc gcggcacgcg ggcgccgcgc
      481 agcaacaaaa gccttcagat gtttgtgttg tgcaaacgca cccacgccgc tcgagtgcga
      541 gagcagcttc gggtcgttat tcagtcccgc aagccgcgca agtattacac gcgatcttcg
      601 gacgggcggc tctgccccgc cgtccccgtg ttcgtccacg agttcgtctc gtccgagcca
      661 atgcgcctcc accgagataa cgtcatgctg gcctcggggg ccgagtaacc gcccccccgc
      721 gccaccctca ctgcccgtcg cggtaccgag ggcctatttc ccatgattcc ttcatatttg
      781 catatacgat acaaggctgt tagagagata attggaatta atttgactgt aaacacaaag
      841 atattagtac aaaatacgtg acgtagaaag taataatttc ttgggtagtt tgcagtttta
      901 aaattatgtt ttaaaatgga ctatcatatg cttaccgtaa cttgaaagta tttcgatttc
      961 ttggctttat atatcttgtg gaaaggacga aacaccgaaa taacgcataa atttggcgtt
     1021 ttagagctag aaatagcaag ttaaaataag gctagtccgt tatcaacttg aaaaagtggc
     1081 accgagtcgg tgcttttttg tctagacagg tctaaaagga tctgcgatcg ctccggtgcc
     1141 cgtcagtggg cagagcgcac atcgcccaca gtccccgaga agttgggggg aggggtcggc
     1201 aattgaacgg gtgcctagag aaggtggcgc ggggtaaact gggaaagtga tgtcgtgtac
     1261 tggctccgcc tttttcccga gggtggggga gaaccgtata taagtgcagt agtcgccgtg
     1321 aacgttcttt ttcgcaacgg gtttgccgcc agaacacagc tgaagcttcg aggggctcgc
     1381 atctctcctt cacgcgcccg ccgccctacc tgaggccgcc atccacgccg gttgagtcgc
     1441 gttctgccgc ctcccgcctg tggtgcctcc tgaactgcgt ccgccgtcta ggtaagttta
     1501 aagctcaggt cgagaccggg cctttgtccg gcgctccctt ggagcctacc tagactcagc
     1561 cggctctcca cgctttgcct gaccctgctt gctcaactct acgtctttgt ttcgttttct
     1621 gttctgcgcc gttacagatc caagctgtga ccggcgccta caccggtgcc accatggact
     1681 ataaggacca cgacggagac tacaaggatc atgatattga ttacaaagac gatgacgata
     1741 agatggcccc aaagaagaag cggaaggtcg gtatccacgg agtcccagca gccgacaaga
     1801 agtacagcat cggcctggac atcggcacca actctgtggg ctgggccgtg atcaccgacg
     1861 agtacaaggt gcccagcaag aaattcaagg tgctgggcaa caccgaccgg cacagcatca
     1921 agaagaacct gatcggagcc ctgctgttcg acagcggcga aacagccgag gccacccggc
     1981 tgaagagaac cgccagaaga agatacacca gacggaagaa ccggatctgc tatctgcaag
     2041 agatcttcag caacgagatg gccaaggtgg acgacagctt cttccacaga ctggaagagt
     2101 ccttcctggt ggaagaggat aagaagcacg agcggcaccc catcttcggc aacatcgtgg
     2161 acgaggtggc ctaccacgag aagtacccca ccatctacca cctgagaaag aaactggtgg
     2221 acagcaccga caaggccgac ctgcggctga tctatctggc cctggcccac atgatcaagt
     2281 tccggggcca cttcctgatc gagggcgacc tgaaccccga caacagcgac gtggacaagc
     2341 tgttcatcca gctggtgcag acctacaacc agctgttcga ggaaaacccc atcaacgcca
     2401 gcggcgtgga cgccaaggcc atcctgtctg ccagactgag caagagcaga cggctggaaa
     2461 atctgatcgc ccagctgccc ggcgagaaga agaatggcct gttcggaaac ctgattgccc
     2521 tgagcctggg cctgaccccc aacttcaaga gcaacttcga cctggccgag gatgccaaac
     2581 tgcagctgag caaggacacc tacgacgacg acctggacaa cctgctggcc cagatcggcg
     2641 accagtacgc cgacctgttt ctggccgcca agaacctgtc cgacgccatc ctgctgagcg
     2701 acatcctgag agtgaacacc gagatcacca aggcccccct gagcgcctct atgatcaaga
     2761 gatacgacga gcaccaccag gacctgaccc tgctgaaagc tctcgtgcgg cagcagctgc
     2821 ctgagaagta caaagagatt ttcttcgacc agagcaagaa cggctacgcc ggctacattg
     2881 acggcggagc cagccaggaa gagttctaca agttcatcaa gcccatcctg gaaaagatgg
     2941 acggcaccga ggaactgctc gtgaagctga acagagagga cctgctgcgg aagcagcgga
     3001 ccttcgacaa cggcagcatc ccccaccaga tccacctggg agagctgcac gccattctgc
     3061 ggcggcagga agatttttac ccattcctga aggacaaccg ggaaaagatc gagaagatcc
     3121 tgaccttccg catcccctac tacgtgggcc ctctggccag gggaaacagc agattcgcct
     3181 ggatgaccag aaagagcgag gaaaccatca ccccctggaa cttcgaggaa gtggtggaca
     3241 agggcgcttc cgcccagagc ttcatcgagc ggatgaccaa cttcgataag aacctgccca
     3301 acgagaaggt gctgcccaag cacagcctgc tgtacgagta cttcaccgtg tataacgagc
     3361 tgaccaaagt gaaatacgtg accgagggaa tgagaaagcc cgccttcctg agcggcgagc
     3421 agaaaaaggc catcgtggac ctgctgttca agaccaaccg gaaagtgacc gtgaagcagc
     3481 tgaaagagga ctacttcaag aaaatcgagt gcttcgactc cgtggaaatc tccggcgtgg
     3541 aagatcggtt caacgcctcc ctgggcacat accacgatct gctgaaaatt atcaaggaca
     3601 aggacttcct ggacaatgag gaaaacgagg acattctgga agatatcgtg ctgaccctga
     3661 cactgtttga ggacagagag atgatcgagg aacggctgaa aacctatgcc cacctgttcg
     3721 acgacaaagt gatgaagcag ctgaagcggc ggagatacac cggctggggc aggctgagcc
     3781 ggaagctgat caacggcatc cgggacaagc agtccggcaa gacaatcctg gatttcctga
     3841 agtccgacgg cttcgccaac agaaacttca tgcagctgat ccacgacgac agcctgacct
     3901 ttaaagagga catccagaaa gcccaggtgt ccggccaggg cgatagcctg cacgagcaca
     3961 ttgccaatct ggccggcagc cccgccatta agaagggcat cctgcagaca gtgaaggtgg
     4021 tggacgagct cgtgaaagtg atgggccggc acaagcccga gaacatcgtg atcgaaatgg
     4081 ccagagagaa ccagaccacc cagaagggac agaagaacag ccgcgagaga atgaagcgga
     4141 tcgaagaggg catcaaagag ctgggcagcc agatcctgaa agaacacccc gtggaaaaca
     4201 cccagctgca gaacgagaag ctgtacctgt actacctgca gaatgggcgg gatatgtacg
     4261 tggaccagga actggacatc aaccggctgt ccgactacga tgtggaccat atcgtgcctc
     4321 agagctttct gaaggacgac tccatcgaca acaaggtgct gaccagaagc gacaagaacc
     4381 ggggcaagag cgacaacgtg ccctccgaag aggtcgtgaa gaagatgaag aactactggc
     4441 ggcagctgct gaacgccaag ctgattaccc agagaaagtt cgacaatctg accaaggccg
     4501 agagaggcgg cctgagcgaa ctggataagg ccggcttcat caagagacag ctggtggaaa
     4561 cccggcagat cacaaagcac gtggcacaga tcctggactc ccggatgaac actaagtacg
     4621 acgagaatga caagctgatc cgggaagtga aagtgatcac cctgaagtcc aagctggtgt
     4681 ccgatttccg gaaggatttc cagttttaca aagtgcgcga gatcaacaac taccaccacg
     4741 cccacgacgc ctacctgaac gccgtcgtgg gaaccgccct gatcaaaaag taccctaagc
     4801 tggaaagcga gttcgtgtac ggcgactaca aggtgtacga cgtgcggaag atgatcgcca
     4861 agagcgagca ggaaatcggc aaggctaccg ccaagtactt cttctacagc aacatcatga
     4921 actttttcaa gaccgagatt accctggcca acggcgagat ccggaagcgg cctctgatcg
     4981 agacaaacgg cgaaaccggg gagatcgtgt gggataaggg ccgggatttt gccaccgtgc
     5041 ggaaagtgct gagcatgccc caagtgaata tcgtgaaaaa gaccgaggtg cagacaggcg
     5101 gcttcagcaa agagtctatc ctgcccaaga ggaacagcga taagctgatc gccagaaaga
     5161 aggactggga ccctaagaag tacggcggct tcgacagccc caccgtggcc tattctgtgc
     5221 tggtggtggc caaagtggaa aagggcaagt ccaagaaact gaagagtgtg aaagagctgc
     5281 tggggatcac catcatggaa agaagcagct tcgagaagaa tcccatcgac tttctggaag
     5341 ccaagggcta caaagaagtg aaaaaggacc tgatcatcaa gctgcctaag tactccctgt
     5401 tcgagctgga aaacggccgg aagagaatgc tggcctctgc cggcgaactg cagaagggaa
     5461 acgaactggc cctgccctcc aaatatgtga acttcctgta cctggccagc cactatgaga
     5521 agctgaaggg ctcccccgag gataatgagc agaaacagct gtttgtggaa cagcacaagc
     5581 actacctgga cgagatcatc gagcagatca gcgagttctc caagagagtg atcctggccg
     5641 acgctaatct ggacaaagtg ctgtccgcct acaacaagca ccgggataag cccatcagag
     5701 agcaggccga gaatatcatc cacctgttta ccctgaccaa tctgggagcc cctgccgcct
     5761 tcaagtactt tgacaccacc atcgaccgga agaggtacac cagcaccaaa gaggtgctgg
     5821 acgccaccct gatccaccag agcatcaccg gcctgtacga gacacggatc gacctgtctc
     5881 agctgggagg cgacaaaagg ccggcggcca cgaaaaaggc cggccaggca aaaaagaaaa
     5941 aggagggcag aggaagtcta ctaacatgcg gtgacgtgga ggagaatccc ggccctatgg
     6001 agagcgacga gagcggcctg cccgccatgg agatcgagtg ccgcatcacc ggcaccctga
     6061 acggcgtgga gttcgagctg gtgggcggcg gagagggcac ccccaagcag ggccgcatga
     6121 ccaacaagat gaagagcacc aaaggcgccc tgaccttcag cccctacctg ctgagccacg
     6181 tgatgggcta cggcttctac cacttcggca cctaccccag cggctacgag aaccccttcc
     6241 tgcacgccat caacaacggc ggctacacca acacccgcat cgagaagtac gaggacggcg
     6301 gcgtgctgca cgtgagcttc agctaccgct acgaggccgg ccgcgtgatc ggcgacttca
     6361 aggtggtggg caccggcttc cccgaggaca gcgtgatctt caccgacaag atcatccgca
     6421 gcaacgccac cgtggagcac ctgcacccca tgggcgataa cgtgctggtg ggcagcttcg
     6481 cccgcacctt cagcctgcgc gacggcggct actacagctt cgtggtggac agccacatgc
     6541 acttcaagag cgccatccac cccagcatcc tgcagaacgg gggccccatg ttcgccttcc
     6601 gccgcgtgga ggagctgcac agcaacaccg agctgggcat cgtggagtac cagcacgcct
     6661 tcaagacccc catcgccttc gccagatccc gcgctcagtc gtccaattct gccgtggacg
     6721 gcaccgccgg acccggctcc accggatctc gctaagaatt cctagagctc gctgatcagc
     6781 ctcgactgtg ccttctagtt gccagccatc tgttgtttgc ccctcccccg tgccttcctt
     6841 gaccctggaa ggtgccactc ccactgtcct ttcctaataa aatgaggaaa ttgcatcgca
     6901 ttgtctgagt aggtgtcatt ctattctggg gggtggggtg gggcaggaca gcaaggggga
     6961 ggattgggaa gagaatagca ggcatgctgg ggagcggccg ctggcatttc agtgtcgggt
     7021 gacgagcgcg atccggccgg gatcctagga ccccaaaagt ttgtctgcgt attccagggc
     7081 ggggctcagt tgaatctccc gcagcacctc taccagcagg tccgcggtgg gctggagaaa
     7141 ctcggccgtc ccggggcagg cggtcgtcgg gggtggaggc gcggcgccca ccccgtgtgc
     7201 cgcgcctggc gtctcctctg ggggcgaccc gtaaatggtt gcagtgatgt aaatggtgtc
     7261 cgcggtccag accacggtca aaatgccggc cgtggcgctc cgggcgcttt cgccgcgcga
     7321 ggagctgacc caggagtcga acggatacgc gtacatatgg gcgtcccacc cgcgttcgag
     7381 cttctggttg ctgtcccggc ctataaagcg gtaggcacaa aattcggcgc gacagtcgat
     7441 aatcaccaac agcccaatgg gggtgtgttg gataacaacg cctccgcgcg gcaggcggtc
     7501 ctggcgctcc cggccccgta ccatgatcgc gcgggtgccg tactcaaaaa catgcaccac
     7561 ctgcgcggcg tcgggcagtg cgctggtcag cgaggccctg gcgtggcata ggctatacgc
     7621 gatggtcgtc tgtggattgg acat
//
```
